# Fasciclin 2 Cooperates with Discs Large to Maintain Epithelial Architecture

**DOI:** 10.64898/2026.05.24.727532

**Authors:** Tara M. Finegan, Michael W. Linhoff, Hannah Rice, Sahel Ghasemzadeh, Zach Wright, Kathryn E. Neville, Tyler J. Wilson, Evan W. Ost, Nicholas Lowe, Adrian Dunivan, Oscar Andrade Mendoza, Christian M. Cammarota, Dan T. Bergstralh

## Abstract

Cell adhesion molecules of the immunoglobulin superfamily (IgCAMs) coordinate adhesive interactions with intracellular organization during tissue morphogenesis. In the *Drosophila* follicular epithelium, epithelial maintenance depends on reintegration, a process in which mitotically displaced cells reincorporate into the epithelial monolayer. Previous work identified the IgCAMs Fasciclin 2 (Fas2) and Neuroglian (Nrg) as parallel, partially redundant regulators of reintegration, but the intracellular mechanisms linking adhesion to reintegration remained unclear. Here, we show that Fas2 supports reintegration through two mechanistically distinct modes: a transmembrane mode and a GPI-linked mode. Although both contribute to reintegration, the transmembrane mechanism is more effective and depends on stabilization of a cortical Fas2 pool through intracellular coupling. Using yeast two-hybrid screening, genetics, and fluorescence recovery after photobleaching (FRAP), we identify the scaffold protein Discs large (Dlg1) as a functional intracellular partner of transmembrane Fas2. Partial disruption of Dlg1 preferentially sensitizes epithelia in which the parallel Nrg-dependent reintegration mechanism is compromised, consistent with Dlg1 functioning primarily within the Fas2-dependent reintegration arm. While Dlg1 is not required for Fas2 membrane localization, Dlg1 disruption increases the mobile fraction of transmembrane Fas2, indicating that Dlg1 promotes retention of a stabilized cortical Fas2 pool. Together, these findings support a model in which epithelial reintegration depends on coordinated adhesion–scaffold coupling and reveal mechanistic parallels between epithelial reintegration and IgCAM-dependent processes in the developing nervous system.

## Introduction

Epithelial tissues must maintain cohesive architecture while simultaneously accommodating cellular remodeling. During development, cells divide, rearrange, change shape, and move relative to one another without compromising tissue integrity. How cell-cell adhesion systems remain mechanically and spatially coordinated during these dynamic events remains a central question in tissue morphogenesis. Cell adhesion molecules (CAMs) of the immunoglobulin superfamily are characterized by extracellular domains containing variable numbers of immunoglobulin-like and fibronectin type III domains, which mediate homophilic and heterophilic binding interactions essential for tissue organization and morphogenesis (Halaby et al. 1999). In the nervous system, IgCAMs direct axon pathfinding, fasciculation, and synapse formation through coordinated interactions with cortical scaffolds and the cytoskeleton (Sanes and Zipursky 2020; Maness and Schachner 2007). Analagous mechanisms also operate in epithelia, where cells must maintain adhesion while undergoing rearrangement, division, and shape change. How IgCAM-based adhesion is coupled to intracellular machinery to support epithelial organization remains incompletely understood.

The *Drosophila* follicular epithelium provides a tractable system for studying these mechanisms. During early oogenesis, follicle cells divide within a polarized epithelial monolayer surrounding the germline cyst. Some daughter cells are transiently displaced apically from the monolayer and subsequently reintegrate. Failure of reintegration produces “popped-out” cells positioned above the epithelium, providing a quantifiable readout of epithelial organization defects (Bergstralh et al. 2015). Previous work identified two IgCAMs, Neuroglian (Nrg) and Fasciclin 2 (Fas2), as central regulators of this process (Bergstralh et al. 2015; Cammarota et al. 2020). Simultaneous disruption of Nrg and Fas2 causes a supra-additive increase in reintegration failure, indicating that these proteins function through parallel, partially redundant mechanisms rather than within a single linear pathway (Cammarota et al. 2020).

Fas2 is the *Drosophila* ortholog of vertebrate Neural Cell Adhesion Molecule (NCAM) (Harrelson and Goodman 1988; Grenningloh et al. 1991; Finegan and Bergstralh 2020). Like NCAM, Fas2 is expressed as multiple alternatively spliced isoforms that differ in membrane attachment and intracellular organization. Five Fas2 isoforms contain transmembrane domains and intracellular regions capable of scaffold interaction, whereas two isoforms are linked to the membrane through glycosylphosphatidylinositol (GPI) anchors and lack intracellular domains (Neuert et al. 2020). This structural diversity makes Fas2 particularly useful for dissecting how adhesive interactions are coupled to intracellular machinery during epithelial morphogenesis.

Our previous work suggested that reintegration depends not only on Fas2-mediated adhesion, but also on intracellular coupling mediated by the Fas2 cytoplasmic domain (Cammarota et al. 2020). This raised two major questions. First, which Fas2 isoforms support reintegration in the follicular epithelium? Second, what intracellular factors interact with transmembrane Fas2 to organize reintegration behavior?

Here, we show that reintegration is supported by two partially redundant Fas2 mechanisms: a transmembrane mode and a GPI-linked mode. We further identify the scaffold protein Discs large (Dlg1) as a functional intracellular partner of transmembrane Fas2. Our findings indicate that Dlg1 is not required for Fas2 membrane localization, but instead promotes retention of a stabilized cortical Fas2 pool. Together, these results support a model in which epithelial reintegration depends on coordinated adhesion–scaffold coupling and reveal mechanistic parallels between epithelial reintegration and IgCAM-dependent processes in the developing nervous system.

## Results

### Fas2 expression and localization in the follicular epithelium

Because reintegrating cells must generate traction as they incorporate into the monolayer, reintegration is expected to require stable coupling between the adhesion molecule and cortex. Consistent with this, our previous rescue experiments suggested that Fas2 supports epithelial reintegration through both its adhesive ectodomain and its intracellular domain, leading to the prediction that the relevant reintegration factor should be a transmembrane isoform (Cammarota et al. 2020). To evaluate this prediction, we asked which Fas2 isoforms are present in proliferative-stage follicle cells, where reintegration occurs. The seven annotated isoforms are distinguished primarily by their C-termini: five have transmembrane (TM) domains and the remaining two (PB and PC) are linked to the membrane by a GPI-anchor (Figure 1A). Because only the transmembrane forms contain an intracellular domain, distinguishing between these classes is essential for interpreting mechanism.

**Figure 1.**
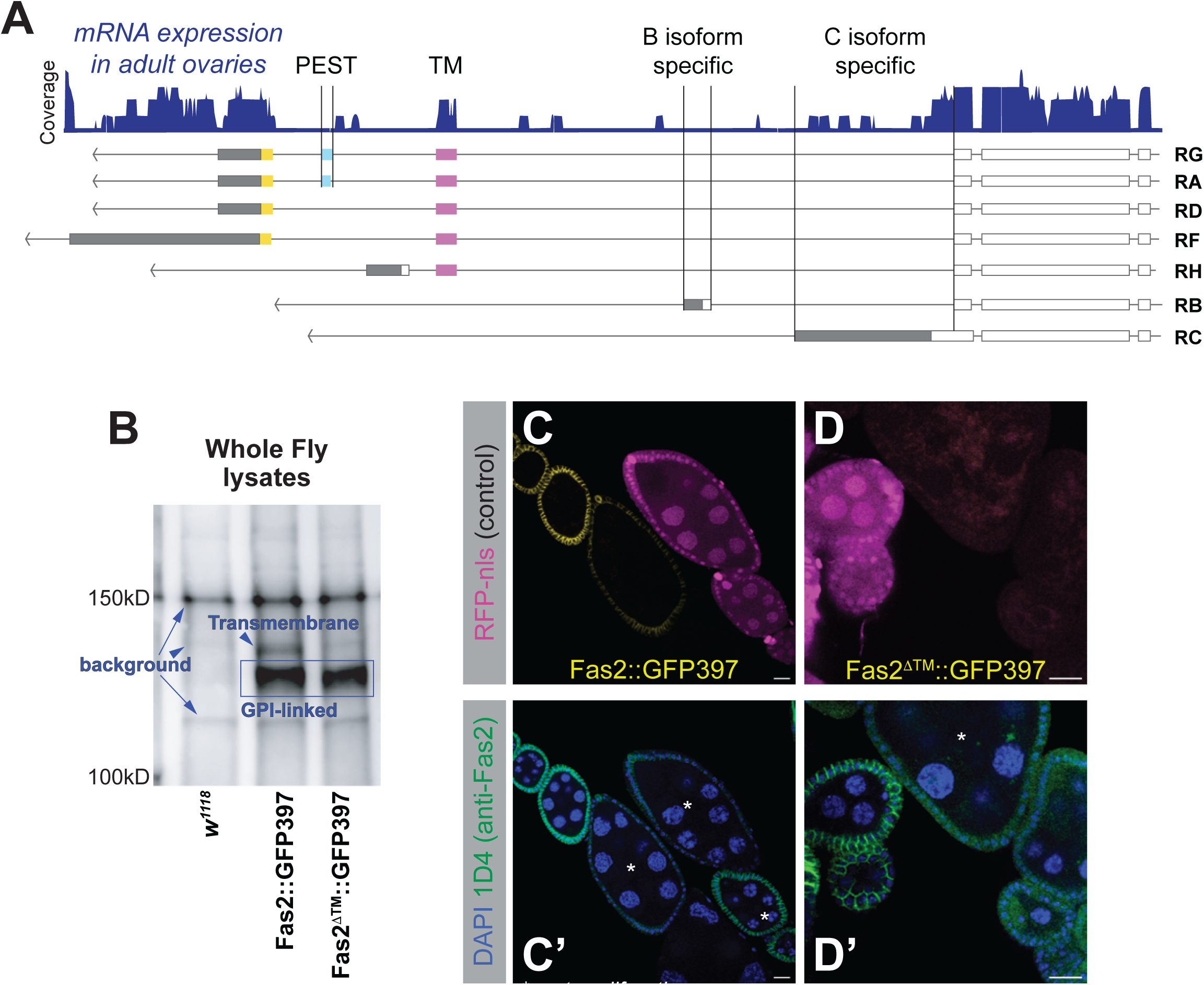
Fas2 protein trap alleles reveal distinct isoform populations. **A)** The exons encoding C-termini of the seven Fas2 isoforms and their expression in adult, mated flies. Expression data was mined from the FlyCellAtlas. **B)** Western blot analysis of whole-fly lysates probed with anti-GFP antibody. *Fas2::GFP397* reveals two bands consistent with TM and GPI-linked isoforms; *Fas2ΔTM::GFP397* displays a single band consistent with GPI-linked isoforms. **C and D)** Expression and localization of fluorescent Fas2 variants in proliferative-stage egg chambers. The 1D4 antibody, which only recognizes transmembrane isoforms, is shown for comparison. RFP-nls expressing egg chambers do not express tagged-Fas2 and provide a negative control. Post-proliferative egg chambers are marked with an asterisk. In these experiments we included egg chambers expressing RFP-nls, but not tagged Fas2, as a positive control for the antibody. Scale bars = 20μm.

We first examined mRNA expression to determine which isoforms are expressed in the ovary. Transmembrane isoforms are well supported; of three cDNA clones isolated from egg chambers at developmental stages 1-6 (the developmental window in which follicle cells are proliferative), two (GM05745 and GM25119) are sequences shared by the TM, but not GPI-linked, isoforms (Rubin et al. 2000). RNASeq data mined from the FlyCellAtlas (H. Li et al. 2022; Gramates et al. 2022) likewise indicate expression of transmembrane isoforms in ovaries, though expression of exons unique to isoforms PA and PG are not detected (Figure 1A). These exons encode a PEST (rich in proline, glutamic acid, serine, and threonine) sequence that targets proteins for degradation (Rechsteiner and Rogers 1996). Expression of the unusual PH isoform is also not supported. Expression of the exon unique to GPI-linked isoform PC is detected, whereas the exon unique to PB is not (Figure 1A).

We investigated further using the protein trap *Fas2::GFP397.* This allele carries a GFP insertion positioned immediately upstream of the transmembrane domain of TM isoforms or at the C-terminus of the GPI-linked PB isoform (Supplemental Figure 1A) (Morin et al. 2001). It does not report GPI-linked isoform PC. A derivative line, *Fas2^ΔTM^::GFP397*, introduces a premature stop codon within the shared TM domain (Neuert et al. 2020), eliminating expression of all TM isoforms. This is confirmed by western blot. In *Fas2::GFP397* lysates made from whole flies, two bands were detected at approximately 125kDa and 115kDa (Figure 1B). Because transmembrane (TM) Fas2 isoforms range from ∼93–98 kDa and GPI-linked isoforms are predicted to be 86 kDa (PB) and 90 kDa (PC), these bands are consistent with tagging of at least one TM isoform and the GPI-linked PB isoform. In agreement, *Fas2^ΔTM^::GFP397* lysates displayed a single band consistent with a GPI-linked isoform, as disruption of the shared TM domain eliminates all TM variants (Figure 1B).

The expression of at least one TM isoform can be surmised from previous work. The anti-Fas2 monoclonal antibody 1D4 recognizes a signal at epithelial cell-cell boundaries in follicle cells, and this signal drops rapidly after proliferation ceases at developmental stage 6 (Gomez et al. 2012; Bergstralh et al. 2015; Cammarota et al. 2020; Finegan et al. 2024). Consistent with expectation, Fas2::GFP397 localizes to follicle cell–cell borders in proliferative egg chambers (Figure 1C), whereas neither GFP signal nor 1D4 immunoreactivity is detected at borders in *Fas2^ΔTM^::GFP397* tissue (Figure 1D).

Together, these results indicate that the proliferative follicle epithelium expresses at least one transmembrane isoform, and that this isoform(s) does not contain the PEST sequence. Of the two GPI-linked isoforms, only expression of PC is supported.

### Fas2 supports reintegration through two partially redundant modes

Having established that both transmembrane and GPI-linked Fas2 isoforms are expressed in proliferative follicle cells, we next asked whether these isoforms differ in localization behavior and reintegration function. Another Fas2 protein trap line, *Fas2^CPTI000483^* (henceforth *Fas2::YFP483*), carries a Venus/YFP tag inserted into an early *Fas2* intron (Supplemental Figure 1A) (Lowe et al. 2014). Based on its position, this tag is expected to label all isoforms. However, western blot analysis of *Fas2::YFP483* lysates revealed only one clear band, corresponding in size to a GPI-linked isoform (Supplemental Figure 2A). This does not indicate a loss of TM expression, as staining with 1D4 revealed cortical signal in Fas2::YFP483 follicle cells (Supplemental Figure 2B). It is also not completely explained by exon skipping, as RT-PCR shows that the YFP exon is present in at least some TM isoform mRNAs (Supplemental Figure 2C). Another possibility is that the YFP tag is proteolytically cleaved. Western blot analysis using a gradient gel, which permits resolution of smaller proteins but not individual Fas2 isoforms, revealed a GFP-sized band in Fas2::YFP483 lysates that was absent from Fas2::GFP397 controls (Supplemental Figure S2E), consistent with proteolytic release of the fluorescent tag. This could impact the TM isoforms preferentially or affect all isoforms similarly, with the apparent absence of TM bands by western instead reflecting the substantially lower abundance of TM Fas2 relative to the dominant GPI-linked population. Consistent with the possibility of cleavage, AlphaFold modeling predicts that the YFP tag is positioned before an exposed linker extending from the Fas2 ectodomain (Supplemental Figure 2D).

Because Fas2::YFP483 preferentially reports the GPI-linked Fas2 pool by western blot, it provided an opportunity to compare the localization behavior of GPI-linked and TM Fas2 isoforms *in vivo*. Both Fas2::YFP483 and 1D4 signal were strongly reduced at *Fas2^EB112^* (null) clone borders (Figure 2A). In addition, despite their distinct modes of membrane attachment, both GPI-linked and TM Fas2 isoforms require trans interactions for stable enrichment at follicle cell borders.

**Figure 2.**
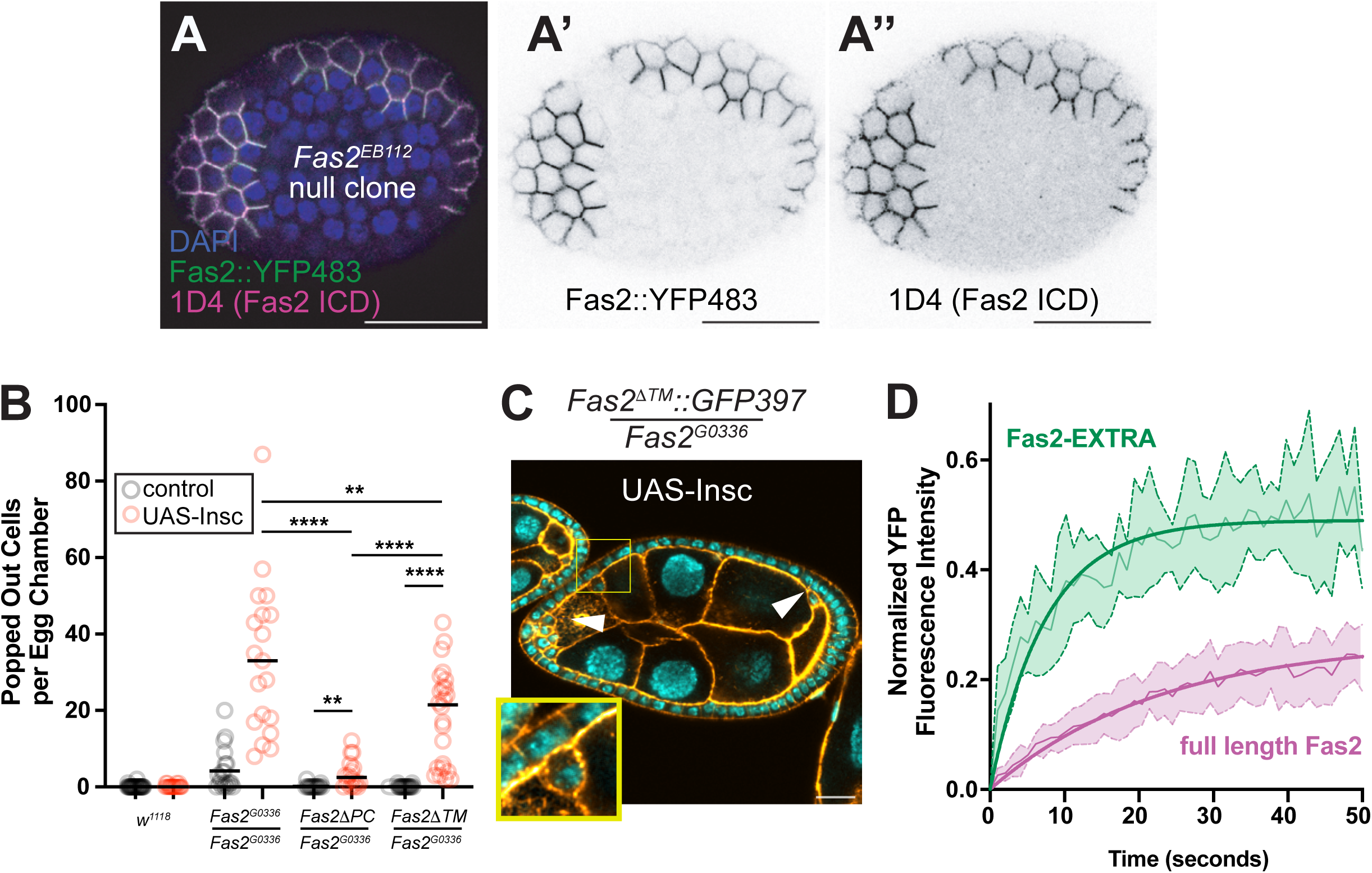
Fas2 isoforms expressed in proliferative follicle epithelium. **A)** Clonal expression of Fas2::YFP483 apposed to *Fas2^EB112/EB112^* cells in the follicle epithelium. Both Fas2::YFP483 signal and 1D4 immunoreactivity are lost from clone borders. **B)** Quantification of popped-out cell numbers (per egg chamber) in *Fas2* mutant conditions shows that Fas2^TM^ and Fas2^PB^ isoforms (expressed in Fas2^ΔPC^ and Fas2^ΔTM^ tissue, respectively) contribute different amounts of reintegration function. **C)** Popped-out cells (white arrowheads) are observed in Fas2^ΔTM^::GFP397/Fas2^G0336^ mutant tissue. Another popped out cell (yellow box) is shown as an enlarged inset. Scale bar = 20μm. **D)** FRAP analysis at follicle cell–cell borders in proliferative-stage egg chambers. Averaged recovery curves for Fas2::GFP397 and Fas2::YFP483 are shown. *p* < 0.001 using Welch’s unpaired two-tailed t-test on area under the curve using trapezoidal integration

We next addressed the question of which isoforms drive epithelial cell reintegration. Reintegration in the follicle epithelium is a quantifiable behavior; failure produces countable “popped-out” cells that remain apically displaced from the underlying monolayer. These cells are readily identified in fixed egg chambers because of their rounded morphology, apical position, and the presence of a distinct border that separates them from the underlying monolayer.

We previously studied the role played by Fas2 in reintegration using the UAS-GAL4 system to express rescue transgenes – either full length Fas2; Fas2-INTRA (the intracellular domain); or Fas2-EXTRA (the extracellular domain) – in *Fas2^G0336^* null mutant clones (Cammarota et al. 2020). (The latter two are attached to the membrane with a transmembrane domain). Whereas the full length Fas2 rescued cell reintegration, neither Fas2-INTRA nor Fas2-EXTRA were sufficient (Cammarota et al. 2020), suggesting that Fas2’s role in cell reintegration relies on both its extracellular function, namely adhesion, and on its intracellular domain (ICD). We therefore expected that reintegration requires a transmembrane isoform, and that a GPI-linked isoform could not mediate reintegration alone. Contrary to this expectation, we did not detect popped-out cells in egg chambers from flies heterozygous for *Fas2^G0336^* and *Fas2Δ^TM^::GFP397*, in which only the GPI-linked Fas2PC isoform should be expressed (Figure 2B, black circles).

We therefore propose a revised model: Fas2 can function in two distinct modes - a GPI-anchored mode and a transmembrane mode – either of which can support reintegration. Two observations support this model: Firstly, we did not find popped out cells in *Fas2^G0336^*/*Fas2^ΔPC^* egg chambers. (Given that Fas2PB is not expressed in ovaries, we did not expect and did not observe popped-out cells in *Fas2^G0336^*/*Fas2^ΔPB^::GFP397* egg chambers (Supplemental Figure 2F). Thus, TM isoform(s) are sufficient to support reintegration in the absence of GPI-linked Fas2. Secondly, whereas our earlier experiments expressed UAS-Fas2 variants in a *Fas2* null background, *Fas2^ΔTM^::GFP397* allows for expression of the GPI-linked isoform Fas2PC. Thus, a GPI-anchored mode remains functional and allows for reintegration.

We tested this model using a sensitized system that increases the demand for reintegration. Ectopic expression of Inscuteable can reorient epithelial cell divisions so that one daughter is born protruding out from the monolayer (Kraut et al. 1996; Egger et al. 2007; Bergstralh et al. 2015). This manipulation potentiates the need for reintegration, increasing the number of popped-out cells in *Fas2^G0336^*/*Fas2^G0336^* mutant follicle epithelium by ∼6-fold (Figure 2B and shown previously (Bergstralh et al. 2015; Cammarota et al. 2020)). (Importantly, the mutant cells in these experiments are generated clonally and comprise > 60%, but not all, of the tissue.) When Inscuteable is expressed, popped-out cells are readily detected in *Fas2^G0336^*/*Fas2^ΔTM^*(Figure 2B,C) and, to a significantly lesser degree, in *Fas2^G0336^*/*Fas2^ΔPC^*egg chambers. These results are consistent with our model; both the TM isoform(s) and the GPI-linked Fas2PC isoform contribute some but not all reintegration function. In the absence of Inscuteable, that amount of function is sufficient to rescue completely.

To determine whether these functionally distinct modes reflect differences in membrane behavior, we performed Fluorescence Recovery After Photobleaching (FRAP) on full-length transmembrane Fas2 and Fas2-EXTRA, which lacks the intracellular domain. After photobleaching, Fas2-EXTRA fluorescence returned rapidly and to a higher plateau, whereas full-length Fas2 recovered more slowly and to a lower level (Figure 2D). These differences indicate that a larger fraction of Fas2-EXTRA is free to exchange within the membrane, while a substantial portion of full-length Fas2 remains stably retained at cell–cell contacts. Thus, the ICD limits Fas2 mobility and promotes formation of a stabilized membrane-associated pool. Taken together, these results indicate that transmembrane Fas2 requires stabilization through intracellular coupling to promote reintegration. In contrast to this, the less-efficient GPI-linked mode likely operates through extracellular adhesion without stable cortical anchoring.

### Yeast-2-Hybrid screening identifies potential intracellular functional partners for Nrg and Fas2

To identify intracellular partners of Fas2 that contribute to reintegration, we undertook yeast two-hybrid (Y2H) screening against an ovary-derived cDNA library. Because reintegration depends on parallel, partially redundant Nrg- and Fas2-dependent modules (Cammarota et al. 2020), we performed parallel screens using the intracellular domains of each protein as bait.

Only two high-confidence interactors, Ankyrin and Moesin, were recovered in the Nrg screen (Figure 3A). Ankyrin is a cortical anchor for Nrg during axon guidance (Suter and Forscher 2000; Enneking et al. 2013) and performs the same role in epithelial cell reintegration (Cammarota et al. 2020). The Nrg FERM-binding (4.1 protein-, Ezrin-, Radixin-, and Moesin-binding) domain is thought to interact directly with Moesin to mediate axon-axon interactions in the mushroom body (Siegenthaler et al. 2015), and we find that expression of a transgenic variant of Nrg lacking the FERM-binding domain provides only partial rescue of the *Nrg^14^* (null) popped-out cell phenotype (Figure 3B). (As in Figure 2B, this experiment was performed in the presence of UAS-Inscuteable to sensitize the assay.) These results validate the screening approach. They also provide further evidence that Nrg’s roles in the developing nervous system and the proliferative follicular epithelium are mechanistically similar, as suggested previously (Cammarota et al. 2020).

**Figure 3.**
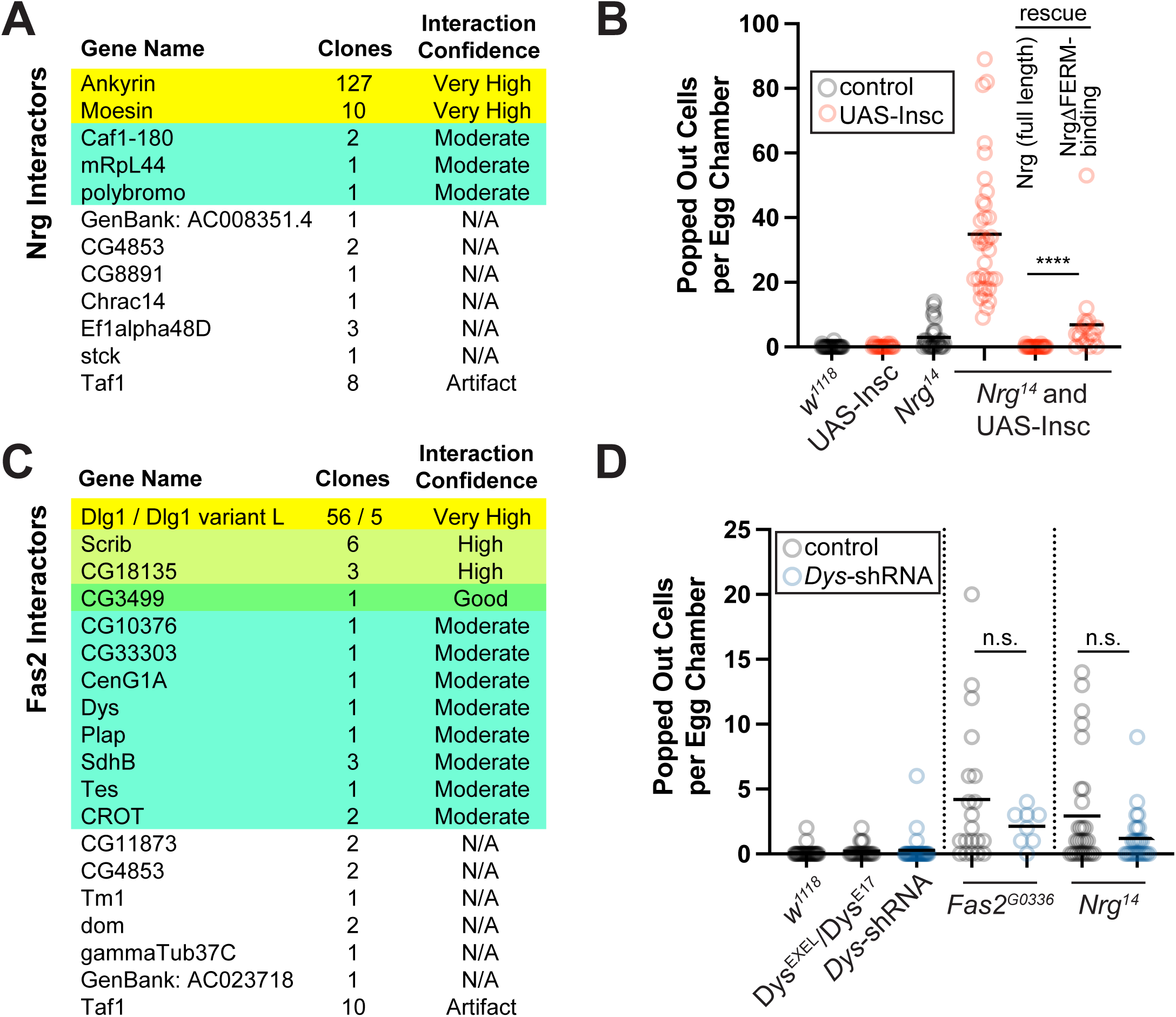
Yeast two-hybrid identifies Dlg1 and other Fas2 interactors. **A)** Summary of Y2H screen using the Nrg intracellular domain as bait. Ankyrin and Moesin are the highest-confidence interactors. **B)** Quantification of popped-out cells shows that an Nrg transgene lacking the FERM-binding domain allows for only partial rescue of reintegration failure in *Nrg^14^* mutant tissue. **C)** Summary of Y2H screen using the Fas2 intracellular domain as bait. Discs large 1 is the most frequently recovered interactor. **D)** Quantification of popped-out cells shows that *Dys* disruption does not cause reintegration failure and that it does not potentiate reintegration failure in *Fas2^G0336^*or *Nrg^14^* mutant tissue.

Significantly more potential interactors were recovered using the intracellular domain of Fas2 as bait (Figure 3C). Discs large (Dlg1) was the highest-confidence interactor in our screen, representing 61 of 103 (>59%) of clones. This result provides confidence, as Fas2 is reported to interact genetically with Dlg1 and can bind the Dlg1 PDZ domains *in vitro* (K. Zito et al. 1997; Thomas et al. 1997). We narrowed the remaining candidates based on expected protein localization, mRNA expression in the follicle epithelium (Supplemental Figure 3A), and interaction confidence (Figure 3C), generating a shortlist of Scribble (Scrib) and Dystrophin (Dys). Interaction with Fas2 was validated using 1X1 Y2H (Supplemental Figure 3B). Although interaction with the uncharacterized protein CG18135 was high, this candidate was not pursued because the Fas2 interaction site maps to the predicted extracellular portion of the protein.

Multiple observations support the possibility of functional interaction between Dys and Fas2: prior affinity capture recovered Dys isoforms using Fas2 as bait (Rees et al. 2011); AlphaFold3 predicts interaction between Fas2 and the Dys ZZ zinc-finger domain (AAs 932–977), corresponding to the region identified in our screen (not shown); and Dys::YFP localizes along proliferative follicle cell borders, although the signal is weak and variable (Supplemental Figure 3C). Despite this evidence, *Dys* disruption did not produce or enhance popped-out cells (Figure 3D), indicating that Fas2-Dys interaction is not required for epithelial reintegration under our assay conditions. We therefore focused subsequent analysis on Dlg1 and Scrib as candidate reintegration factors.

### Fas2 associates with PDZ scaffold proteins through a conserved C-terminal motif

Both Dlg1 and Scrib are multi-PDZ (PSD-95, Dlg, ZO-1) domain scaffolds, and the cytoplasmic domains of most Fas2 transmembrane isoforms (not H) terminate in a Type I PDZ-binding motif (KNSAV in *D. melanogaster*) (Thomas et al. 1997). To address the importance of this domain we compared the C-termini of Fas2 orthologs across arthropods. While the overall intracellular domain is well conserved among insects, it diverges substantially outside this class (Figure 4A). Notably, however, the C-terminal PDZ-binding motif remains clearly identifiable even where surrounding sequence conservation is low, suggesting that PDZ-mediated scaffold interaction is a functionally important feature of transmembrane Fas2 isoforms.

**Figure 4.**
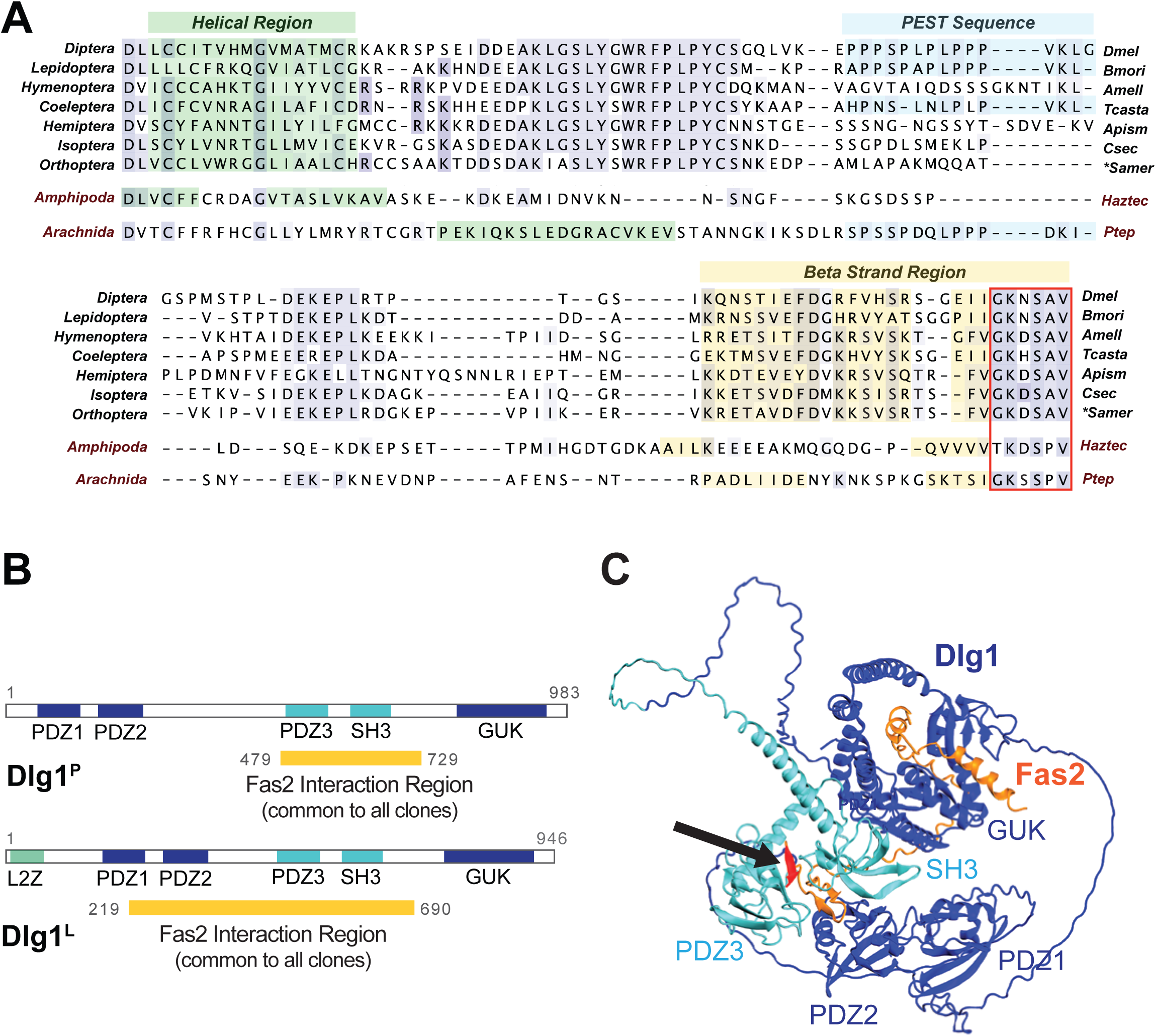
Yeast two-hybrid and structural analyses define the Fas2–Dlg1 interaction interface. **A)** Comparison of C terminal sequences across arthropod species. The C-terminal ∼5 amino acids (KNSAV in *D. melanogaster*) correspond to a Type I PDZ recognition motif (x-x-S/T-x-I/L/V) (Chimura et al. 2011). These motifs are preferentially located at the C-terminus (Chimura et al. 2011). The preceding sequence region is the predicted beta strand, and these strands are associated with “stickiness” that helps to stabilize protein-protein interactions (Stranges et al. 2011). Beta strand structures, though smaller, are also predicted near to the C-termini of Fas2 orthologs in the amphipod crustacean *Hyalella azteca* and the arachnid *Parasteatoda tepidararium*. **B)** Schematics illustrating the domain architecture of Dlg1^P^ and Dlg1^L^, including the interaction regions common to every clone identified through yeast-two hybrid screening for Fas2 interactors. Recovered fragments localize to residues 461–729 (PDZ3-containing region). **C)** AlphaFold structural prediction supporting interaction between Fas2 C-terminus and Dlg1 PDZ3-containing region. Dlg1 is shown in blue except for the Y2H shared interaction region which is shown in cyan. Fas2 cytoplasmic sequence is shown in orange except for the PDZ binding motif which is shown in red (black arrow indicates location).

We also considered the possibility that interaction between Fas2 and PDZ domains is promiscuous, and that our Y2H results may not reflect *in vivo* binding. To explore this, we asked what other PDZ proteins are expressed in the ovary; if Fas2 interacts promiscuously with PDZ domains, these proteins might also be expected to yield positive interaction in the Y2H screen. RNASeq data mined from the FlyCellAtlas (H. Li et al. 2022), shows that many PDZ-domain encoding genes are expressed in the ovary, including but not limited to *Prosap, Spn, polychaetoid, RhoGAP19D,* and *bazooka* (Supplemental Figure 5), increasing our confidence in the *in vivo* relevance of the screen results.

Two Dlg1 isoforms, both of which include a set of three PDZ domains (Kim and Sheng 2004), were identified as Fas2-binding partners in our screen (Figure 3C). Earlier work showed that Fas2 binds to a protein fragment that includes PDZ1 and PDZ2, but not to a fragment that has only PDZ3 (Thomas et al. 1997), leading to the conclusion that Fas2 interacts with one of the first two PDZ domains. Our results do not agree. The number and variability of fragments recovered in our Y2H screen indicate that Fas2 binds Dlg1 residues 461-729, a stretch that includes PDZ3 and SH3 domains but not PDZ1 or PDZ2 (Figure 4B). Structural prediction using AlphaFold3 supports this finding (Figure 4C). The model also predicts close association between Fas2 and the C-terminal GUK domain of Dlg1 (Figure 4C), suggesting the possibility that this domain helps to mediate and/or stabilize the interaction between Fas2 and Dlg1. Consistent with our results, PDZ-SH3-GUK domain tandems have been shown to enhance binding affinity to PDZ-binding motifs when compared to PDZ domain binding alone (Y. Li et al. 2014).

### Scrib disruption causes defective germline cyst separation and misplaced follicle cells

We used RNA interference as a first test to assess whether Scrib disruption produces popped-out cells. UAS-*Scrib*-shRNA was driven by Traffic jam-GAL4, which is active from egg chamber developmental Stage 1 (Figure 5A). Traffic jam-GAL4 is efficient in our system at 29℃ but not 18℃ (Lovegrove et al. 2019). At the higher temperature, we observed gross disorganization and failed egg chamber separation. Though still severe, the phenotype was somewhat weaker when the flies were allowed to develop and eclose at 18℃ then shifted to 29℃ overnight (Figure 5B). These observations suggest that Scrib supports germline cyst encapsulation and/or separation, interlinked processes that occur as a new egg chamber buds off from the germarium, an organ at the anterior end of each ovariole (Duhart et al. 2017). We tested this using GR1-GAL4, which becomes active at developmental Stage ∼3 (Lovegrove et al. 2019), to reduce Scrib protein levels after encapsulation has occurred. We did not observe defective egg chamber separation. We did however identify popped-out cells in these egg chambers. This was not the only disorganization observed, as other misplaced cells appeared to be protruding from, rather than situated apical-to, the monolayer (Figure 5C). Taken together, these results indicate that Scrib promotes organization of the follicle cell epithelium both at the time of cyst encapsulation and later in development and suggest a potential role in reintegration.

**Figure 5.**
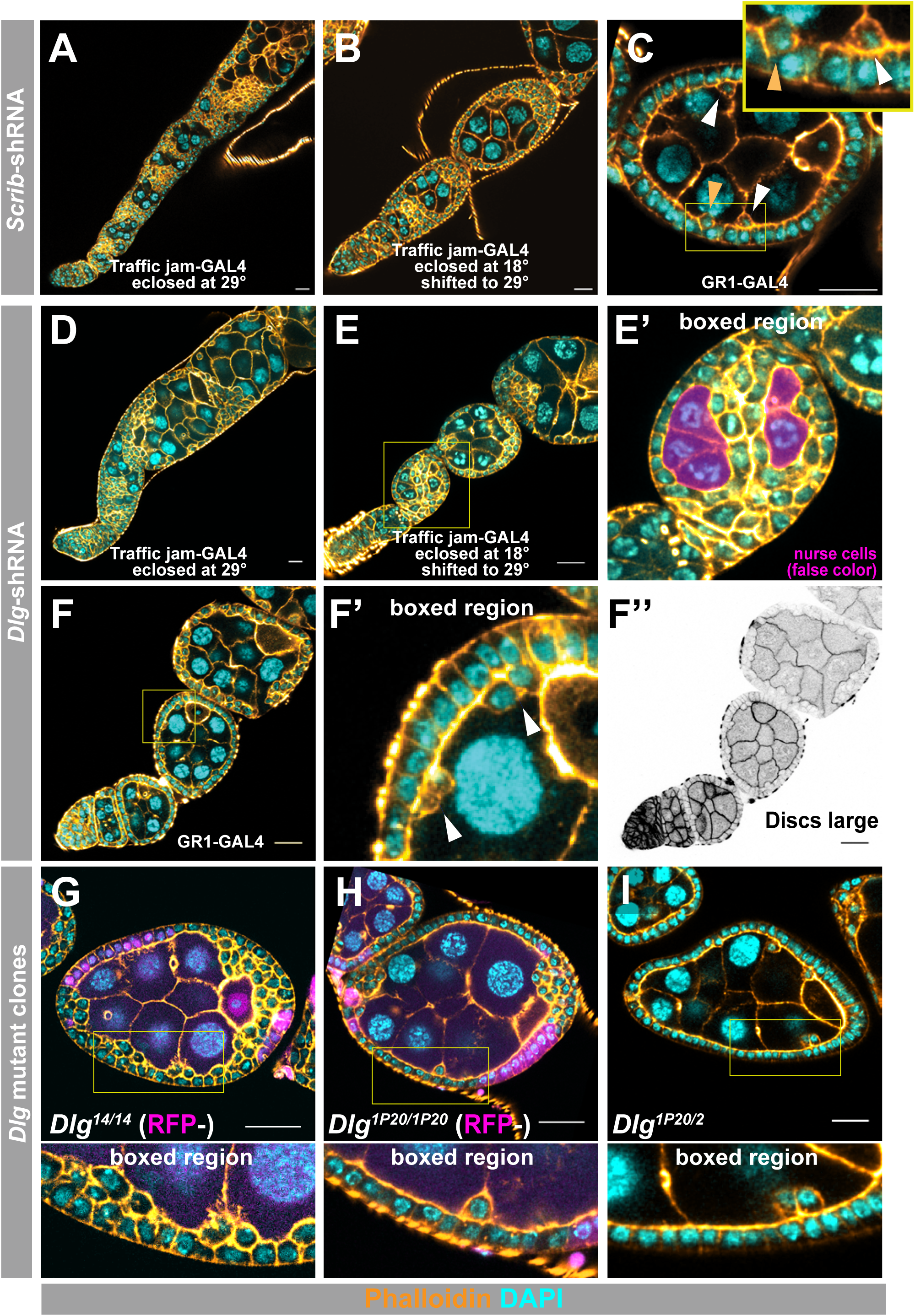
Dlg1 disruption causes defective cyst separation and reintegration phenotypes. A-F) Impact of *Scrib* (A-C) or *Dlg1* (D-F) knockdown on the follicle epithelium. Actin is marked with phalloidin (orange) and DNA with DAPI (cyan). Scale bars set to 20μm. **A, B)** *Scrib*-shRNA driven by Traffic jam-GAL4 results in egg chamber disorganization and failed cyst separation. This phenotype is weaker when the flies develop at 18℃ then switched to 29℃ overnight (B). **C)** *Scrib*-shRNA driven by GR1-GAL4 (post-encapsulation) produces popped-out cells and protruding follicle cells without complete epithelial collapse. **D, E)** *Dlg1*-shRNA driven by Traffic jam-GAL4 results in egg chamber disorganization and failed cyst separation. This phenotype is weaker when the flies develop at 18℃ then switched to 29℃ overnight (E). A magnified image (E’, from the yellow box in E) shows epithelial cells bisecting the interior germline. The nurse (germline) cells are false-colored in magenta. **F)** *Dlg1*-shRNA driven by GR1-GAL4 (post-encapsulation) produces popped-out cells and protruding follicle cells without complete epithelial collapse. A magnified image (F’, from the yellow box in F) shows popped-out cells. Anti-Dlg1 immunostaining (F’’) shows that the appearance of popped-out cells coincides with a decrease in Dlg1 enrichment at cell-cell borders. **G - I)** Popped-out cells are observed in *Dlg1* mutant follicle epithelium (mutant clones in A and B are marked by the absence of RFP). For each of the three mutant genotypes, a magnified image (yellow rectangle) reveals epithelial tissue organization over the main body of the egg chamber.

A potentially confounding issue is that Scrib is well characterized as a polarity regulator and Dlg1-partner protein in the follicular epithelium (reviewed in (Buckley and St Johnston 2022)), raising the possibility that the Scrib phenotype reflects disruption of Dlg1-dependent processes (or vice versa). *In silico* binding competition (AlphaFold3) between the PDZ domains of Dlg1 and Scrib results in preferential binding of the C-terminus of Fas2 to PDZ3 of Dlg1 (example Supplemental Figure 4). For this reason, and in light of the very high confidence interaction detected through Y2H, we chose to focus subsequent analysis on Dlg1.

### Dlg1 disruption causes defective germline cyst separation and misplaced follicle cells

Dlg1 emerged as a particularly strong candidate because it shares functional similarity with the IgCAM reintegration factors identified to date—Nrg, Fas2, and Fas3. All four proteins are A) implicated in axon guidance during development and B) localize to the neuromuscular junction (NMJ), an excitatory synapse (Mendoza et al. 2003; Finegan and Bergstralh 2020). The neural functions of Dlg1 are thought to be isoform-specific, mediated by isoforms termed S97 due to their N-terminal sequence similarity to human SAP97 (Mendoza et al. 2003). Exon-specific RNASeq data (Flybase) indicate that one or more S97 isoforms are expressed in ovaries (not shown). Consistent with this, we identified the S97 isoform Dlg1L as a Fas2-interacting protein in our Y2H screen, confirming its mRNA expression in *Drosophila* ovaries. Taken together, these observations suggest that Dlg1 might function similarly in developing epithelial tissues and the nervous system.

We repeated the knockdown experiments performed previously (Fig 5A,B) and found similar results. At 29℃, *Dlg1*-shRNA driven by Traffic jam-GAL4 caused gross disorganization and failed egg chamber separation (Figure 5D). Again, the phenotype was weaker when the flies were allowed to develop and eclose at 18℃ then shifted to 29℃ overnight (Figure 5E). Using GR1-GAL4, we find popped-out cells (Figure 5F).

While the popped-out cells observed after *Dlg1* knockdown suggest this protein plays a role in reintegration, the accompanying gross disorganization, characterized by a loss of apical-basal cell polarity (Bilder et al. 2000), is a barrier to quantification. We therefore performed a series of genetic disruption experiments using mutant clones, which provide better spatial and temporal control. In agreement with earlier work, *Dlg1^14^*(null; also called *Dlg1^m52^*) mutant tissue is characterized by gross disorganization, which is stronger at the egg chamber poles (Figure 5G). The hypomorphic allele *Dlg1^1P20^* encodes a premature truncation that does not disrupt apical-basal polarity (Bergstralh et al. 2013). Although *Dlg1^1P20^* tissue is disorganized at the egg chamber poles, it forms an orderly monolayer over the main body of the egg chamber, allowing popped-out cells to be identified unambiguously (Figure 5H). Popped-out cells are also observed in tissue transheterozygous for *Dlg1^1P20^*and the temperature-sensitive null allele *Dlg1^2^* (Figure 5I). These results indicate that the contribution of Dlg1 to tissue organization cannot be explained by loss of epithelial polarity alone.

In addition, several observations suggest that Dlg1 supports germline cyst encapsulation and/or egg chamber separation, paralleling our findings for Scrib. Dlg1-knockdown tissue is characterized by failed egg chamber separation, with adjacent chambers sharing a mass of disorganized tissue rather than forming discrete epithelia (Figure 5D,E), and a similar defect is evident in *Dlg1^1P20^*/*Dlg1^2^* egg chambers (Supplemental Figure 6A). The enrichment of disorganization at the poles suggests an early defect in follicle epithelium morphogenesis, consistent with phenotypes observed in tissue mutant for *myospheroid* (Lovegrove et al. 2019), which encodes a β-integrin that participates in encapsulation. We also identify cases in which the follicle epithelium appears to bisect the interior germline (Figure 5E’). Previous studies described invasive follicle cell “tumors” that appear to be detached from the external follicle epithelium in *Dlg1* mutant egg chambers (Goode and Perrimon 1997; Szafranski and Goode 2007), but these interpretations were based on single optical sections. Given this limitation, we tested whether such structures instead represent cross sections through bisected egg chambers arising from defective encapsulation, as we observed in *Dlg1-*knockdown tissue (Figure 5E’). Three-dimensional reconstruction supports this interpretation, revealing that the apparent “tumors” are in fact continuous with the follicular epithelium, as expected if a single epithelial sheet incompletely or aberrantly wraps the germline cyst (Supplemental Figure 6B).

### Dlg1 potentiates reintegration failure in *Nrg*-reduced tissue

Because Dlg1 disruption causes popped-out cells and Dlg1 interacts with Fas2, we hypothesized that Dlg1 participates in the Fas2-dependent reintegration module. If it does, then we would expect partial disruption of Dlg1 to enhance reintegration failure when the parallel, Nrg-dependent reintegration module is compromised. To test this, we made use of another allele combination: *Dlg1^2^* and the weak hypomorph *Dlg1^SW^*, a premature truncation that removes only the last fourteen amino acids (Woods et al. 1996). This allele combination does not lead to popped-out cells (Figure 6A, Supplemental Figure 7A). However, reducing Dlg1 dosage enhances reintegration failure in Nrg-shRNA tissue in a dose-dependent manner: *Dlg1^2^*/+ caused a modest increase in popped-out cells, whereas the *Dlg1^2^*/*Dlg1^SW^*combination enhanced popping-out >3-fold relative to Nrg-shRNA alone (Figure 6A). By contrast, *Dlg1^2^*/*Dlg1^SW^* caused a small, non-significant increase in popped-out cells in Fas2-shRNA tissue (p = 0.11) (Figure 6A). These results support a model in which Dlg1 contributes primarily to the Fas2-dependent arm of reintegration.

**Figure 6.**
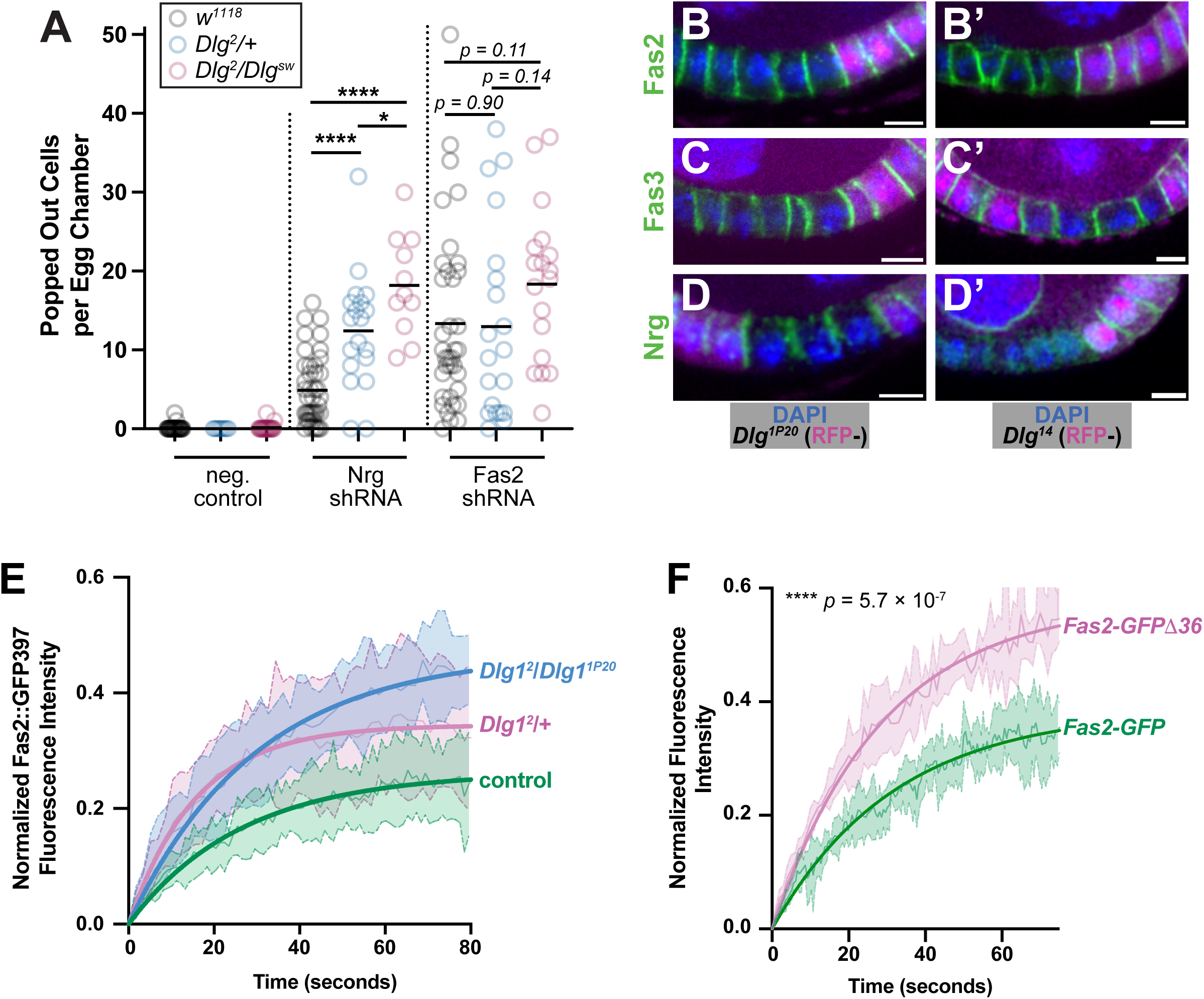
Loss of Dlg1 function potentiates popping out after Nrg disruption. **A)** Mild disruption of Dlg1 increases popped-out cell frequency in *Nrg*-shRNA, but not *Fas2*-shRNA, egg chambers. **B,C**) Localization of Fas2 and Fas3 is preserved in *Dlg1^1P20^*and *Dlg1^14^* clones. **D)** Nrg localizes normally in *Dlg1^1P20^*clones but is lost from cell-cell borders in *Dlg1^14^* clones. Scale bars = 20μm **E)** FRAP analysis demonstrates increased mobility of Fas2::GFP397 in Dlg-disrupted tissue relative to control. Partial disruption of Dlg function (*Dlg1^2^*/+) significantly increases Fas2::GFP397 mobility at cell junctions in Stage 5-6 egg chambers (*p* < 0.0491, Fas2 ::GFP397 Mobile fraction = 26% vs *Dlg1^2^*/+ Mobile fraction = 34%), whereas stronger disruption (*Dlg1^2^*/*Dlg1^1P20^*) results in a further increase in mobility *Dlg1^2^*/*Dlg1^1P20^*(*p* = 0.0121, Mobile fraction = 47%). **F)** A truncated variant of Fas2-TM-GFP lacking the PDZ-binding motif and the adjacent conserved β-strand structure demonstrates statistically significant (*p* < 0.00001) increased mobility at cell junctions in Stage 5-6 egg chambers in comparison to full-length Fas2-TM-GFP (mobile fraction of Fas2 protein full-length Fas2-TM-GFP = 39% vs Fas2-TM-GFPΔ36 = 58%). FRAP statistical significance was calculated using Welch’s unpaired two-tailed t-test on area under the curve using trapezoidal integration.

We next addressed the question of how Dlg1 participates in Fas2-dependent reintegration. A straightforward explanation would be that it participates in Fas2 localization. This is anticipated by previous work showing that Dlg1 promotes efficient clustering of Fas2 at the *Drosophila* neuromuscular junction, though it is not a strict requirement for Fas2 membrane localization in that system (Thomas et al. 1997; Karen Zito et al. 1997). However, we find that Fas2TM isoform(s) are enriched at follicle cell-cell borders in *Dlg1^14^*and *Dlg1^1P20^* mutant tissue (Figure 6B), with no evidence of diminished signal relative to control. Dlg1 therefore is not working at the level of Fas2 membrane localization.

We also examined the other reintegration IgCAMs in *Dlg1^14^*and *Dlg1^1P20^* mutant cells. Fas3 localization is unaffected (Figure 6C). However, while Nrg localizes normally in *Dlg1^1P20^* mutant tissue (Figure 6D), we find that it is reduced or absent from cell-cell borders in *Dlg1^14^*(null) mutant cells (Figure 6D’, Supplemental Figure 7B). This observation places Dlg1 upstream of the Nrg-dependent reintegration mechanism. It does not explain the finding that *Dlg1^2^*/*Dlg1^SW^* enhances reintegration failure in Nrg-shRNA but not Fas2-shRNA tissue. Taken together, these findings indicate that Dlg1 not only contributes to Nrg localization, but separately to the Fas2-dependent reintegration mechanism.

### Dlg1 stabilizes Fas2 at the cortex

Notably, reintegration failure scales non-linearly (exponentially) with disruption of the reintegration machinery (Cammarota et al. 2020). For this reason, we interpret the observed enhancement to Nrg-dependent failure caused by the *Dlg1^2^*/*Dlg1^SW^*combination to be a modest impairment, consistent with the findings that A) *Dlg1^2^*/*Dlg1^SW^* alone does not cause reintegration failure and B) loss of Fas2 enhances Nrg-dependent failure by ∼10-fold (Cammarota et al. 2020).

This modest impairment is consistent with a model in which Dlg1 impacts Fas2TM function while Fas2GPI function remains. A possibility therefore is that while Dlg1 is not required for Fas2TM localization, it does contribute to Fas2 stability. To test this, we performed FRAP on Fas2::GFP397 in control, *Dlg1^2^*/+, and *Dlg1^2^*/*Dlg1^1P20^* tissue. If Dlg1 stabilizes a subset of Fas2 molecules at the cortex, loss of the Dlg1-Fas2 interaction would be expected to reduce the immobile fraction without necessarily altering the exchange rate of the remaining mobile population. Consistent with this, Fas2::GFP397 recovered with broadly similar kinetics across all three genotypes, approaching a steady-state plateau within ∼60–80 seconds (Figure 6E). However, the recovery plateau increased progressively with Dlg1 disruption, with *Dlg1^2^*/+ tissue exhibiting an intermediate phenotype and *Dlg1^2^*/*Dlg1^1P20^*tissue showing the greatest recovery (Figure 6E). We also examined the dynamics of a truncated variant of Fas2TM that lacks the PDZ-binding motif and the adjacent conserved β-strand structure (AAs 1106-1132). This variant is more mobile than the full length control (Figure 6F). Together, these data indicate that Dlg1 promotes retention of a stabilized Fas2 pool at the cortex. Because the *Dlg1^1P20^* allele disrupts the Dlg1 C-terminus, these findings additionally implicate the C-terminal region in stabilization of the transmembrane Fas2 pool.

### Axon guidance genes are broadly expressed in the proliferative follicle epithelium

Reintegration requires mitotic daughter cells to move directionally through the epithelial monolayer while remaining attached to neighboring cells. This resembles axon guidance, in which extending neurites migrate along cellular substrates through adhesive contact. Consistent with this parallel, the reintegration factors identified to date – Nrg and Fas2 (Bergstralh et al. 2015); Fas3 and Ank (Cammarota et al. 2020); Moesin and Dlg1 (this study) - are associated with axon pathfinding in the developing nervous system (Freymuth and Fitzsimons 2017; Koch et al. 2008; Grenningloh et al. 1991; Snow et al. 1989; Hall and Bieber 1997; Mendoza et al. 2003). Scribble, a potential reintegration factor, has likewise been implicated in axon guidance (Walsh et al. 2011).

Several observations suggest that this relationship extends beyond the currently identified reintegration machinery. The Dlg1L neuronal isoform was recovered in our Y2H screen, confirming that it is expressed in follicle cells. Nrg, Fas2, and Fas3 are downregulated after follicle cell proliferation ceases at Stage 6 (reviewed in (Horne-Badovinac and Bilder 2005)), restricting their expression to proliferative stages when reintegration occurs. This temporal alignment suggests that reintegration may draw upon a broader neural adhesion program active in proliferating follicle cells. To test this, we analyzed transcriptomic profiles from the FlyCellAtlas (H. Li et al. 2022), and compared them to later and post-proliferative tissue (stages 6-8) (Figure 7A).

**Figure 7.**
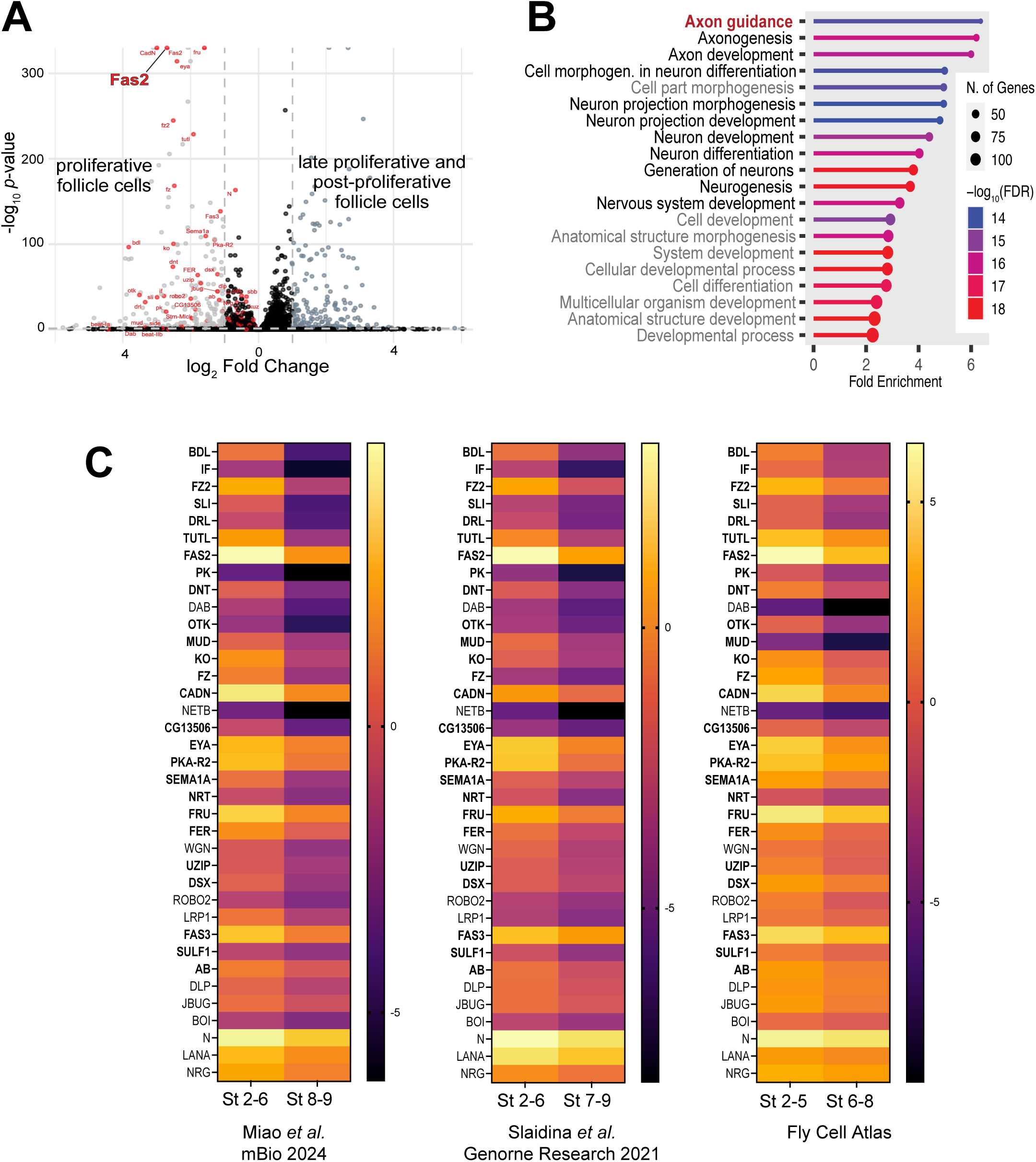
Proliferative follicle epithelium expresses a neural development gene program. **A)** Differential expression analysis comparing proliferative (Stages 2–5 at left) and late/post-proliferative (Stages 6–8 at right) follicle cells using FlyCellAtlas data. Genes in the “Axon Guidance” GO Term are shown in red. **B)** Gene Ontology enrichment showing significant overrepresentation of axon guidance and neuron projection development terms. **C)** Cross-dataset validation of proliferative-stage enrichment of axon guidance genes. Genes marked in bold meet stringent criteria (>2 Fold Change and *p* <0.01 in all three studies). Genes that are not bold meet relaxed criteria (>1.5 Fold change and *p* <0.05 in all three studies). Expression is presented in log_2_. Note that the second category for the Fly Cell Atlas dataset includes some proliferative egg chambers (Stage 6) and for that reason we expect smaller differences.

We identified a set of genes with significantly higher expression in the earlier, more proliferative stages and performed Gene Ontology (GO) analysis on this list using ShinyGo8.0 (Ge et al. 2020). The single most significantly overrepresented GO term is *axon guidance*, with a fold enrichment of ∼6 and a false-discovery rate on the order of 10^-14.^ This is followed by related neuronal developmental terms, including *axonogenesis* and *neuron projection development* (Fig. 7B). As a further test of these expression patterns, we examined two additional datasets that are independent of the FlyCellAtlas and found a large set of axon guidance genes that show consistent enrichment in proliferative stages (Fig. 7C). Notably, the magnitude of differential expression is generally higher in these datasets, consistent with the inclusion of some proliferative egg chambers (Stage 6) within the later-stage category in the FlyCellAtlas. These results are consistent with reintegration drawing on a broader axon guidance gene program active in proliferative follicle cells. This indicates that the axon guidance program – which includes the reintegration machinery - is not restricted to the nervous system but instead constitutes a strategy for cell behavior deployed in both neural and epithelial tissues.

## Discussion

### A Dual-Mode Model for Fas2-Mediated Reintegration

Our molecular dissection of Fas2 reveals that epithelial organization in the *Drosophila* follicle cell monolayer is supported by two mechanistically distinct modes. While both transmembrane (TM) and GPI-linked isoforms contribute to reintegration, they are not functionally equivalent. In the absence of external stress, either mode is sufficient to maintain the monolayer. However, under the sensitized conditions provided by ectopic *Inscuteable* expression, the TM mode proves significantly more effective. This suggests that while GPI-linked Fas2 provides a baseline of homophilic “stickiness,” the TM isoforms provide the mechanical robustness required for active reintegration.

A question raised by these findings is why *Dlg1^1P20^*mutant tissue demonstrates popped-out cells (Figure 5H), given that this mutation should leave at least the Fas2-GPI reintegration mode intact. A potential explanation for these results is that *Dlg1^1P20^* not only disrupts Fas2-TM-dependent reintegration but also spindle orientation (Bergstralh et al. 2013; Neville et al. 2023). This latter activity is expected to cause division misorientation and should thereby increase the need for reintegration.

One additional possibility is that the apparent functional superiority of TM Fas2 reflects isoform abundance rather than intrinsic mechanism. In this model, the weaker activity of GPI-linked Fas2 may reflect its proportional expression rather than – or in addition to – a lack of cortical coupling.

### Dlg1 as a Scaffold for Cortical Stability

The identification of Discs large (Dlg1) as a high-confidence partner for Fas2 provides a mechanistic bridge between extracellular adhesion and intracellular organization. Dlg1’s canonical role in apical-basal polarity relies on a protein region lacking PDZ domains (Khoury and Bilder 2020), and our results suggest at least one of these conserved domains supports epithelial reintegration.

While Dlg1 emerged as the strongest Fas2 interactor in our screen, Scrib was also recovered and Scrib disruption produced popped-out cells (Figure 5C). In many epithelial contexts, Scrib and Dlg1 function as a coordinated unit called the Scribble module (Bonello and Peifer 2018; Ventura et al. 2020; Khoury and Bilder 2020; Bilder et al. 2000). Whether Scrib promotes Fas2-dependent reintegration through a direct interaction with Fas2 or instead through its well-established partnership with Dlg1 remains unclear.

Our FRAP analysis indicates that Dlg1 is not a prerequisite for Fas2 membrane localization but is essential for its cortical retention. In the absence of Dlg1, the mobile fraction of TM Fas2 increases, suggesting that Dlg1-mediated scaffolding transitions Fas2 from a transient adhesive contact to a stable mechanical anchor. This interaction is likely mediated by the C-terminal PDZ-binding motif (KNSAV), which we found to be selectively conserved across arthropod evolution. This motif likely docks into the Dlg1 PDZ3 pocket, a model supported by our Y2H and AlphaFold-Multimer analysis.

### Evolutionary Repurposing of Neural Machinery

The co-option of Fas2, Neuroglian (Nrg), and Dlg1 in the follicular epithelium suggests mechanistic parallels between epithelial reintegration and neural wiring. In the developing *Drosophila* nervous system, Fas2 and Nrg mediate axon fasciculation through adhesive interactions that maintain stable contacts between neighboring axons. Our findings suggest that related IgCAM-based mechanisms operate during epithelial reintegration, where displaced follicle cells must re-establish stable adhesive interactions with neighboring cells as they reincorporate into the monolayer.

The role of Dlg1 in reintegration mirrors its function at the neuromuscular junction (NMJ), where the S97 isoform is required to cluster Fas2 at the postsynaptic membrane (Thomas et al., 1997). In both the synapse and the epithelium, Dlg1 acts as a molecular “clasp,” anchoring the adhesive extracellular domain of Fas2 to the underlying actin cytoskeleton. This anchoring likely allows the follicle cell to generate the mechanical traction necessary to pull itself back into the monolayer.

### Parallel Pathways and Mechanistic Redundancy

The “supra-additive” genetic relationship we observed between Nrg and Fas2 is a hallmark of redundant guidance systems. During midline crossing in the CNS, multiple IgCAMs work in parallel to ensure robust pathfinding. The follicular epithelium employs a similar “belt and braces” strategy, utilizing two independent modules—the Nrg-Ankyrin arm and the Fas2-Dlg1 arm—to ensure that even if one adhesive anchor is compromised, a backup mechanical pathway exists to preserve tissue integrity.

However, the finding that Dlg1 null alleles (but not hypomorphs) also impact Nrg localization suggests Dlg1 may occupy a hierarchical position, maintaining general epithelial polarity while simultaneously acting as a specific lateral anchor for Fas2.

### Broader Implications and Evolutionary Divergence

Notably, recent work in the cnidarian *Nematostella vectensis* identified an NCAM-family IgCAM as an essential component of apical epithelial organization during early embryogenesis. In that system, depletion of NCAM2 disrupts apical junction integrity and blocks gastrulation (Postnikova et al., 2025). The identification of IgCAM-dependent epithelial architecture in an early-branching metazoan reinforces the idea that these adhesion systems represent an evolutionarily conserved module that predates nervous system specialization. This suggests that the neural functions of these proteins may be a specialized derivation of an ancestral epithelial role; before the evolution of complex synapses, primitive metazoan epithelial sheets already required mechanisms to divide and rearrange without losing barrier continuity. Understanding the ancient Fas2-Dlg1 partnership may therefore provide insights into the fundamental rules of tissue integrity, with broad implications for human diseases involving defective cell adhesion, including cancer metastasis and developmental disorders.

## Supplemental Figure Legends

**Supplemental Figure 1**

A) Schematic of annotated Fas2 isoforms, including five transmembrane (TM) isoforms and two GPI-linked isoforms. Positions of the CPTI000483 (Fas2::YFP483) and GFP397 insertions are indicated relative to signal peptide, transmembrane domain, and cytoplasmic regions. B) Alignment of the C-terminal regions of annotated Fas2 isoforms. Predicted transmembrane domains, helical regions, β-strand regions, and the conserved C-terminal PDZ-binding motif are indicated. Transmembrane (TM) isoforms share a conserved intracellular domain containing a predicted juxtamembrane α-helix and C-terminal β-strand region, whereas GPI-linked isoforms terminate prior to the intracellular region.

**Supplemental Figure 2**

A) Anti-GFP western blot using lysates prepared from control and Fas2::YFP483 flies across increasing volumes (2–20 μL). Both standard and heat map color schemes are used to show a single tagged protein band in Fas2::YFP483 lysates. B) Clonal expression of Fas2::YFP483 in proliferative-stage follicle cells. The 1D4 monoclonal antibody, which recognizes the intracellular domain of TM Fas2 isoforms, recognizes signal both in and outside the clone. Scale bars are 20 μm. C) RT-PCR analysis of Fas2 isoforms amplified from ovarian cDNA. The forward primer is within the YFP exon, and reverse primers are isoform-specific as indicated. Products corresponding to predicted GPI-linked and transmembrane isoforms were detected, whereas water controls were negative. D) Alphafold3 model for the translated YFP-Fas2^PA^ product from Fas2::YFP483. The N-terminus contains a 6xHis tag and two StrepTags flanking the YFP. The linker between the tagged region and the Fas2 ectodomain is indicated with an arrow. E) Anti-GFP western blot of whole fly lysates run on a 4-20% gel. Lysates from Fas2::YFP483 flies show some evidence for a YFP-sized band suggesting a limited amount of cleavage has occurred. The GFP control lane is lysate from COS7 cells transfected with an eGFP expression vector. F) Quantification of popped-out cells in control (*w^1118^*) and *Fas2ΔPB*/*Fas2^G0336^*egg chambers.

**Supplemental Figure 3**

A) Heat map showing relative expression of genes encoding potential Fas2-interacting proteins at different stages of follicle epithelium development. B) 1×1 Y2H validation of interactions between Fas2 and Scrib, Dys, and CG18135. C) Dystrophin::YFP localizes to epithelial cell-cell junctions.

**Supplemental Figure 4**

Example AlphaFold model (one of five) shows that the Fas2 PDZ binding motif prefers to bind Dlg1 PDZ3 over Scrib PDZ2. The Fas2 cytoplasmic sequence used in Figure 4C was modeled with isolated PDZ3 of Dlg1 and PDZ2 of Scrib. The highest confidence prediction is shown, and in all five predictions, the Fas2 PDZ binding motif prefers to interact with the Dlg1 PDZ domain.

**Supplemental Figure 5**

Heat map showing relative expression of PDZ genes in the ovary across multiple annotated cell types.

**Supplemental Figure 6**

A) Egg chambers in *Dlg1^1P20^/Dlg1^2^* flies can demonstrate epithelial disorganization, including failure of egg chamber separation. B) Three-dimensional reconstruction shows that a cluster of follicle cells that appears “detached” in one plane is in fact connected to the exterior epithelium. Phalloidin (actin) is revealed in orange and DAPI (DNA) in cyan. A zoom of the highlighted region (yellow square) is shown in the lower right. Scale bars are 20μm.

**Supplemental Figure 7**

A) Popped-out cells are not observed in *Dlg1^SW^*/*Dlg1^2^*egg chambers. B) Nrg is lost from cell-cell borders in *Dlg1^14^*clones (*en face* view). Scale bars = 20μm

## Acknowledgments

We are grateful to the labs of Mark Peifer, Scott Williams, and Holly Lovegrove; the University of Missouri Fly Club; and members of the Finegan-Bergstralh lab for their questions and comments. This work was supported by an NSF CAREER award (PI: Bergstralh) and NIH Grant R01GM125839 (PI: Bergstralh). CPTI fly lines were obtained from KYOTO Drosophila Stock Center at Kyoto Institute of Technology.

## Data availability

The authors affirm that all data necessary for verifying the conclusions are included in the text, figures, and tables.

## Methods

### Drosophila genetics

A list of alleles and transgenes used in this study is found in Supplementary Table 1. With regard to *Dystrophin* stocks: *Df(3R)Exel6184* removes almost all of the Dys gene (Taghli-Lamallem et al. 2008), and the TRiP.JF01118 line was shown effective in prior studies (Sidisky et al. 2024; Clayworth and Auld 2025). Flies were maintained on BDSC Cornmeal Food, generated using the Nutri-Fly Bloomington Formulation.

### Reagents

A list of reagents used in this study is found in Supplementary Table 2. With regard to the 1D4 anti-Fas2 monoclonal antibody; the first reported study to make use of this antibody does not include a description of the epitope (Seeger et al. 1993), but according to the depositor (C.S. Goodman, 2001), the hybridoma maintained at the Developmental Studies Hybridoma Bank produces an antibody raised against “103 a.a. at the intracellular C-terminus” of Fas2. A later report describes the antigen as “103 amino acids of the C-terminal transmembrane domain of FasII, beginning at amino acid 771” (Kumar et al. 2009). Given that the PH transcript was not yet identified at the time the antibody was generated, we conclude that it recognizes one or more of the transmembrane isoforms PA, PD, PG, and PF.

### Immunostaining

Ovaries were fixed for 15 minutes in 10% Formaldehyde and 0.2% Tween in Phosphate Buffered Saline (PBS-T). Following fixation, tissues were incubated in blocking solution (10% Bovine Serum Albumin in PBS) for ∼1hr at room temperature. Primary and secondary immunostainings were performed overnight in PBS-T. Three washes of about 5 minutes each in PBS-T were carried out between stainings and after the secondary staining. Primary and secondary antibodies were used at a concentration of 1:150. Samples were mounted using Vectashield with DAPI (Vector Laboratories).

### Western blots

Whole flies or ovaries were lysed in the following buffer: 1% Triton X-100, 150 mM NaCl, 20 mM Tris-HCl, 1 mM EGTA, 1 mM EDTA, with the addition of protease inhibitors (Roche Applied Science). Samples were heat denatured in Laemmli buffer and resolved on precast 7.5% Mini-PROTEAN TGX gels (BioRad). Precision Plus Protein Kaleidoscope prestained protein standards (Bio-Rad) were loaded to determine target size. Proteins were wet transferred to PVDF for one hour at 100 V in Tris-glycine buffer at 4°C. Transfer membranes were blocked for 30 minutes in TBS-T (10 mM Tris, pH 7.4, 150 mM NaCl, 0.1% Tween-20) with 5% milk. Blots were probed with anti-GFP antibody in blocking buffer overnight at 4°C. Blots were washed six times with TBS-T and probed with HRP conjugated anti-rabbit secondary antibody for one hour in blocking buffer. After six washes with TBS-T, target proteins were detected using SuperSignal West Pico PLUS chemiluminescent substrate (Thermo Scientific) and imaged using a ChemiDoc Imaging System (Bio-Rad). After ECL detection, protein loading was assessed by amido black staining.

### Imaging

Microscopy was performed using either a Leica SP8 point scanning confocal (63x/1.4 HCX PL Apo CS oil lens) or an Andor Dragonfly Spinning Disk confocal microscope (20x/0.75 air objective or 63x/1.4 oil objective). Images were collected with LAS X or the Andor Fusion software respectively. Images were analyzed and prepared using using FiJi (Schindelin et al. 2012). Image preparation included minor processing (Smooth) in FiJi.

### Mitotic clones

*Fas2^EB112^, Fas^G0336^, Nrg^14^, Dlg1^14^,* and *Dlg1^1P20^* clones were generated using hsflp as part of the following background stock: RFP-nls hsflp FRT19A. We used the verified *Fas2^EB112^* line (Finegan et al. 2024). Clones were induced by incubating larvae or pupae at 37° for two out of every twelve hours over a period of at least two days. Adult females were dissected at least two days after the last heat shock. Ectopic protein expression was accomplished using the UAS-GAL4 system (Brand and Perrimon 1993). UAS-GAL4 flies were maintained at 29° for 48 hours before dissection.

### AlphaFold modeling

All structural models were generated using data derived from Alphafold 3 (https://alphafoldserver.com/). The model for figure display was generated using VMD (Humphrey et al. 1996), and analysis of confidence scores was performed using ChimeraX (Goddard et al. 2018). For the Fas2 and Dlg1 L model, the cytoplasmic sequence from Fas2-PA was entered along with the full sequence of Dlg1-PL (FlyBase FBpp0111725). Out of the five .cif files generated, all five showed binding of the Fas2 C-terminal tetrapeptide to PDZ3 of Dlg1 L. Confidence in the Fas2 PBM docking in PDZ3 was determined by examining mean pLDDT values. Across the five predictions, the four residues show a mean pLDDT of approximately 80.1 indicating high confidence in the peptides position in the binding pocket. The -2 residue, S96, exhibited the highest mean confidence at 83.4 consistent with the critical role for this position in binding to PDZ domains. Predicted aligned error (PAE) values were also examined in ChimeraX at the interface between the Fas2 C-terminal tetrapeptide and PDZ3 of Dlg1 L. Low PAE values were observed for the Fas2-PDZ3 interface indicating high confidence in the peptide’s specific position within the binding pocket. For in silico binding competitions between Dlg1 PDZ3 and scrib PDZ2, the isolated PDZ domains were modeled with the cytoplasmic sequence of Fas2-PA. In all five predictions, the C-terminal tetrapeptide of Fas2 bound to Dlg1 PDZ3.

Additional secondary structure prediction for Fas2 (Supplemental Figure 1B) relied on BetaPRO (Cheng and Baldi 2005), PROTEUS2.0 (Montgomerie et al. 2008), and JPRED (Drozdetskiy et al. 2015).

### Expression data

mRNA expression data was mined from FlyBase (Gramates et al. 2022; Öztürk-Çolak et al. 2024) and/or the Fly Cell Atlas (H. Li et al. 2022) as indicated.

### Quantitative PCR

Whole female larvae were processed for RNA isolation with the Zymo Research Quick-RNA MiniPrep Kit according to kit instructions. Isolated RNA was eluted in 50 µL of DNase/RNase-Free Water. Yield and purity was assessed on a Thermo Scientific Nanodrop One UV-Vis Spectrophotometer (Thermo-Fisher Scientific Cat. No 13-400-518). Samples with 260/230 ratios below 1.7 were rejected for purity.

For a 2-step RT-PCR, 1000 ng of total RNA was converted to cDNA with the Invitrogen SuperScript II First-Strand Synthesis System according to kit directions. PCR of the converted cDNA was performed with NEB Taq 5X Master Mix. 2 µL of the cDNA product or H_2_O only was added to PCR reactions of the following final concentrations: 200 nM Forward Primer (Table 3), 200 nM Reverse Primer (Table 3), Taq 5X Master Mix to 1X, NF H_2_O to 25 µL. Samples were mixed and run to the following PCR program:

|  |  |
| --- | --- |
| 1X | 95°C, 0:30 |
| 35X | 95°C, 0:30 ; 58°C, 1:00 ; 68°C, 3:30 |
| 1X | 68°C, 5:00 |

### Yeast Two-Hybrid

Yeast two-hybrid screening was performed by Hybrigenics Services, S.A.S. (Evry, France). Coding sequences used as bait were PCR-amplified and cloned into pB66 as C-terminal fusions to the Gal4 DNA-binding domain. Constructs were verified by sequencing and used to screen a random-primed cDNA prey library cloned into pP6. The pB66 vector derives from pAS2ΔΔ (Fromont-Racine et al. 1997), while pP6 is based on pGADGH (Fields 1993).

Screens and directed 1×1 interaction assays were performed using a mating strategy between CG1945 (mata) and YHGX13 (Y187 ade2-101::loxP-kanMX-loxP, matα) yeast strains as previously described (Fromont-Racine et al. 1997). Diploid yeast were selected on medium lacking tryptophan and leucine (DO-2) to confirm the presence of bait and prey plasmids. Protein interactions were assessed using growth on selective medium lacking tryptophan, leucine, and histidine (DO-3), supplemented with 3-aminotriazole where necessary to suppress bait autoactivation. For directed assays, interactions and controls were tested as streaks from three independent yeast clones. Negative controls included bait constructs tested against empty prey vector (pP6) and prey constructs tested against empty bait vector (pB66). The SMAD–SMURF interaction was used as a positive control (Colland et al. 2004).

For library screens, prey fragments from positive colonies were amplified by PCR and sequenced at both 5′ and 3′ junctions. Corresponding interacting proteins were identified by automated alignment against GenBank (NCBI). Interactions were assigned a Predicted Biological Score (PBS), an indicator of interaction confidence, as previously described (Formstecher et al. 2005). The PBS incorporates both local parameters, including redundancy and independence of prey fragments and reading frame distribution, and global parameters reflecting the frequency of prey recovery across unrelated screens performed with the same library. Scores were categorized from A (highest confidence) to D (lowest confidence), with categories E and F denoting highly connected or previously identified false-positive prey domains, respectively. PBS scores positively correlate with the biological relevance of identified interactions (Wojcik et al. 2002; Rain et al. 2001) (Rain et al. 2001; Wojcik et al. 2002).

A complete list of clones identified in each screen is provided in Table 4 (Nrg) and Table 5 (Fas2).

### RNASeq analysis

The differential expression of genes between developmental stages of the *Drosophila melanogaster* ovarian follicle cell population was analyzed using three publicly available single-cell RNA sequencing datasets. Two datasets were retrieved from the NCBI Gene Expression Omnibus: GSE162192 (Slaidina et al. 2021), obtained as pre-processed Seurat objects, and the Fly Cell Atlas ovary dataset, available at https://flycellatlas.org/, retrieved as an .h5ad file and converted to h5Seurat format with SeuratDisk. The third dataset (Miao et al. 2024) was obtained directly from the authors, as the relevant subset is not part of the public deposit, although the data are also available through NCBI BioProjects (Accession: PRJNA1108780); only the *Wolbachia*-uninfected control sample was used. All datasets were processed and analyzed using the Seurat package (version 5.x) within the RStudio environment. Author-provided cluster annotations were retained as cell identities; the Fly Cell Atlas object was restricted to follicle cell clusters and the Miao object to uninfected cells. Complete lists of genes meeting the >1.5-fold change and *p* < 0.05 criteria are provided in Table 6.

### Misplaced Cell Counting

Quantification of popped-out cells was performed on Stage 6-8 egg chambers using at least 3 dissections of at least 5 flies each. For analyses of clonal mutants, the number of extralayer cells was quantified in egg chambers that were at least 60% mutant. Popped-out cells were quantified manually. The entire depth of each egg chamber was examined using confocal microscopy. Images shown are of representative sagittal planes from each condition, while each data point reflects all misplaced cells in a single egg chamber.

### FRAP

Ovaries were dissected in 10S oil onto 1.5 glass coverslips. FRAP experiments were performed on a Leica SP8 confocal coupled to a Dmi8 inverted microscope. Images were acquired using a Galvano scanner with a 100x/1.44 PLAN APO oil lens using LASX software. Photobleaching was performed on user-defined 2μm^2^ region of interest on cell-cell junctions longer than 2μm using a 488nm argon laser over the ROI at 100% laser power 5 times. Post-bleaching acquisition was performed at 10x zoom over a 18 x 15 µm region at 1s intervals for 150 seconds, or until the sample moved out of the plane of focus. One image of the imaging field of view was acquired pre-bleaching.

### FRAP Analysis

Fluorescence intensity was quantified in FIJI/ImageJ using a 5-pixel-wide linear region of interest (ROI) corresponding to the bleached junction region. Mean fluorescence intensity within the ROI was measured in pre- and post-bleach images. To correct for junction drift, the ROI position was manually adjusted in the x–y plane throughout the time series. FRAP recovery curves were fit using non-linear one-phase association analysis in GraphPad Prism.

## REFERENCES

Bergstralh, Dan T., Holly E. Lovegrove, and Daniel St Johnston. 2013. “Discs Large Links Spindle Orientation to Apical-Basal Polarity in Drosophila Epithelia.” Current Biology : CB 23 (17): 1707–12.

Bergstralh, Dan T., Holly E. Lovegrove, and Daniel St Johnston. 2015. “Lateral Adhesion Drives Reintegration of Misplaced Cells into Epithelial Monolayers.” Nature Cell Biology 17 (11): 1497–503.

Bilder, D., M. Li, and N. Perrimon. 2000. “Cooperative Regulation of Cell Polarity and Growth by Drosophila Tumor Suppressors.” *Science (New York*, NY).

Bonello, Teresa T., and Mark Peifer. 2018. “Scribble: A Master Scaffold in Polarity, Adhesion, Synaptogenesis, and Proliferation.” Journal of Cell Biology 218 (3): 742–56. 10.1083/jcb.201810103.

Brand, A. H., and N. Perrimon. 1993. “Targeted Gene Expression as a Means of Altering Cell Fates and Generating Dominant Phenotypes.” Development 118 (2): 401–15.

Buckley, Clare E., and Daniel St Johnston. 2022. “Apical-Basal Polarity and the Control of Epithelial Form and Function.” Nature Reviews. Molecular Cell Biology 23 (8): 559–77. 10.1038/s41580-022-00465-y.

Cammarota, Christian, Tara M. Finegan, Tyler J. Wilson, Sifan Yang, and Dan T. Bergstralh. 2020. “An Axon-Pathfinding Mechanism Preserves Epithelial Tissue Integrity.” Current Biology 30 (24): 5049–5057.e3. 10.1016/j.cub.2020.09.061.

Cheng, J., and P. Baldi. 2005. “Three-Stage Prediction of Protein -Sheets by Neural Networks, Alignments and Graph Algorithms.” Bioinformatics 21 (Suppl 1): i75–84. 10.1093/bioinformatics/bti1004.

Chimura, Takahiko, Thomas Launey, and Masao Ito. 2011. “Evolutionarily Conserved Bias of Amino-Acid Usage Refines the Definition of PDZ-Binding Motif.” BMC Genomics 12 (1): 300. 10.1186/1471-2164-12-300.

Clayworth, Katherine V., and Vanessa J. Auld. 2025. “Dystroglycan Mediates Polarized Deposition of Laminin and Axon Ensheathment by Wrapping Glia.” Development 152 (10): dev204391. 10.1242/dev.204391.

Colland, Frédéric, Xavier Jacq, Virginie Trouplin, et al. 2004. “Functional Proteomics Mapping of a Human Signaling Pathway.” Genome Research 14 (7): 1324–32. 10.1101/gr.2334104.

Drozdetskiy, Alexey, Christian Cole, James Procter, and Geoffrey J. Barton. 2015. “JPred4: A Protein Secondary Structure Prediction Server.” Nucleic Acids Research 43 (W1): W389–94. 10.1093/nar/gkv332.

Duhart, Juan Carlos, Travis T. Parsons, and Laurel A. Raftery. 2017. “The Repertoire of Epithelial Morphogenesis on Display: Progressive Elaboration of Drosophila Egg Structure.” *Mechanisms of Development*, April.

Egger, Boris, Jason Q. Boone, Naomi R. Stevens, Andrea H. Brand, and Chris Q. Doe. 2007. “Regulation of Spindle Orientation and Neural Stem Cell Fate in the Drosophila Optic Lobe.” Neural Development 2: 1.

Enneking, Eva-Maria, Sirisha R. Kudumala, Eliza Moreno, et al. 2013. Transsynaptic Coordination of Synaptic Growth, Function, and Stability by the L1-Type CAM Neuroglian. 11 (4): e1001537.

Fields, Stanley. 1993. “The Two-Hybrid System to Detect Protein-Protein Interactions.” Methods 5 (2): 116–24. 10.1006/meth.1993.1016.

Finegan, Tara M., and Dan T. Bergstralh. 2020. “Neuronal Immunoglobulin Superfamily Cell Adhesion Molecules in Epithelial Morphogenesis: Insights from Drosophila.” Philosophical Transactions of the Royal Society B: Biological Sciences 375 (1809): 20190553. 10.1098/rstb.2019.0553.

Finegan, Tara M., Christian Cammarota, Oscar Mendoza Andrade, Audrey M. Garoutte, and Dan T. Bergstralh. 2024. “Fas2EB112: A Tale of Two Chromosomes.” G3 Genes|Genomes|Genetics 14 (5): jkae047. 10.1093/g3journal/jkae047.

Formstecher, Etienne, Sandra Aresta, Vincent Collura, et al. 2005. “Protein Interaction Mapping: A Drosophila Case Study.” Genome Research 15 (3): 376–84. 10.1101/gr.2659105.

Freymuth, Patrick S., and Helen L. Fitzsimons. 2017. “The ERM Protein Moesin Is Essential for Neuronal Morphogenesis and Long-Term Memory in Drosophila.” Molecular Brain 10 (1): 41. 10.1186/s13041-017-0322-y.

Fromont-Racine, M., J. C. Rain, and P. Legrain. 1997. “Toward a Functional Analysis of the Yeast Genome through Exhaustive Two-Hybrid Screens.” Nature Genetics 16 (3): 277–82. 10.1038/ng0797-277.

Ge, Steven Xijin, Dongmin Jung, and Runan Yao. 2020. “ShinyGO: A Graphical Gene-Set Enrichment Tool for Animals and Plants.” Bioinformatics 36 (8): 2628–29. 10.1093/bioinformatics/btz931.

Goddard, Thomas D., Conrad C. Huang, Elaine C. Meng, et al. 2018. “UCSF ChimeraX: Meeting Modern Challenges in Visualization and Analysis.” Protein Science: A Publication of the Protein Society 27 (1): 14–25. 10.1002/pro.3235.

Gomez, Juan Manuel, Ying Wang, and Veit Riechmann. 2012. “Tao Controls Epithelial Morphogenesis by Promoting Fasciclin 2 Endocytosis.” The Journal of Cell Biology 199 (7): 1131–43.

Goode, S., and N. Perrimon. 1997. “Inhibition of Patterned Cell Shape Change and Cell Invasion by Discs Large during Drosophila Oogenesis.” Genes & Development 11 (19): 2532–44.

Gramates, L. Sian, Julie Agapite, Helen Attrill, et al. 2022. “FlyBase: A Guided Tour of Highlighted Features.” Genetics 220 (4): iyac035. 10.1093/genetics/iyac035.

Grenningloh, G., E. J. Rehm, and C. S. Goodman. 1991. “Genetic Analysis of Growth Cone Guidance in Drosophila: Fasciclin II Functions as a Neuronal Recognition Molecule.” Cell 67 (1): 45–57.

Halaby, D. M., A. Poupon, and J. Mornon. 1999. “The Immunoglobulin Fold Family: Sequence Analysis and 3D Structure Comparisons.” Protein Engineering 12 (7): 563–71. 10.1093/protein/12.7.563.

Hall, Stephen G., and Allan J. Bieber. 1997. “Mutations in the Drosophila Neuroglian Cell Adhesion Molecule Affect Motor Neuron Pathfinding and Peripheral Nervous System Patterning.” Journal of Neurobiology 32 (3): 325–40. 10.1002/(SICI)1097-4695(199703)32:3%3C325::AID-NEU6%3E3.0.CO;2-9.

Harrelson, A. L., and C. S. Goodman. 1988. “Growth Cone Guidance in Insects: Fasciclin II Is a Member of the Immunoglobulin Superfamily.” *Science (New York*, NY) 242 (4879): 700–708.

Horne-Badovinac, Sally, and David Bilder. 2005. “Mass Transit: Epithelial Morphogenesis in the Drosophila Egg Chamber.” Developmental Dynamics : An Official Publication of the American Association of Anatomists 232 (3): 559–74.

Humphrey, William, Andrew Dalke, and Klaus Schulten. 1996. “VMD: Visual Molecular Dynamics.” Journal of Molecular Graphics 14 (1): 33–38. 10.1016/0263-7855(96)00018-5.

Khoury, Mark J., and David Bilder. 2020. “Distinct Activities of Scrib Module Proteins Organize Epithelial Polarity.” Proceedings of the National Academy of Sciences 117 (21): 11531–40. 10.1073/pnas.1918462117.

Kim, Eunjoon, and Morgan Sheng. 2004. “PDZ Domain Proteins of Synapses.” Nature Reviews Neuroscience 5 (10): 771–81. 10.1038/nrn1517.

Koch, Iris, Heinz Schwarz, Dirk Beuchle, Bernd Goellner, Maria Langegger, and Hermann Aberle. 2008. “*Drosophila* Ankyrin 2 Is Required for Synaptic Stability.” Neuron 58 (2): 210–22. 10.1016/j.neuron.2008.03.019.

Kraut, Rachel, William Chia, Lily Yeh Jan, Yuh Nung Jan, and Jürgen A. Knoblich. 1996. “Role of Inscuteable in Orienting Asymmetric Cell Divisions in Drosophila.” Nature 383 (6595): 50–55. 10.1038/383050a0.

Kumar, Abhilasha, S. Fung, Robert Lichtneckert, Heinrich Reichert, and Volker Hartenstein. 2009. “Arborization Pattern of Engrailed-Positive Neural Lineages Reveal Neuromere Boundaries in the Drosophila Brain Neuropil.” Journal of Comparative Neurology 517 (1): 87–104. 10.1002/cne.22112.

Li, Hongjie, Jasper Janssens, Maxime De Waegeneer, et al. 2022. “Fly Cell Atlas: A Single-Nucleus Transcriptomic Atlas of the Adult Fruit Fly.” *Science*, ahead of print, March 4. World. 10.1126/science.abk2432.

Li, Youjun, Zhiyi Wei, Yan Yan, Qingwen Wan, Quansheng Du, and Mingjie Zhang. 2014. “Structure of Crumbs Tail in Complex with the PALS1 PDZ–SH3–GK Tandem Reveals a Highly Specific Assembly Mechanism for the Apical Crumbs Complex.” Proceedings of the National Academy of Sciences 111 (49): 17444–49. 10.1073/pnas.1416515111.

Lovegrove, Holly E., Dan T. Bergstralh, and Daniel St Johnston. 2019. “The Role of Integrins in *Drosophila* Egg Chamber Morphogenesis.” *Development*, January 1, dev.182774. 10.1242/dev.182774.

Lowe, Nick, Johanna S. Rees, John Roote, et al. 2014. “Analysis of the Expression Patterns, Subcellular Localisations and Interaction Partners of Drosophila Proteins Using a pigP Protein Trap Library.” Development 141 (20): 3994–4005.

Maness, Patricia F., and Melitta Schachner. 2007. “Neural Recognition Molecules of the Immunoglobulin Superfamily: Signaling Transducers of Axon Guidance and Neuronal Migration.” Nature Neuroscience 10 (1): 1. 10.1038/nn1827.

Mendoza, Carolina, Patricio Olguín, Gabriela Lafferte, et al. 2003. “Novel Isoforms of Dlg Are Fundamental for Neuronal Development inDrosophila.” The Journal of Neuroscience 23 (6): 2093–101. 10.1523/JNEUROSCI.23-06-02093.2003.

Miao, Yun-heng, Wei-hao Dou, Jing Liu, Da-wei Huang, and Jin-hua Xiao. 2024. “Single-Cell Transcriptome Sequencing Reveals That Wolbachia Induces Gene Expression Changes in Drosophila Ovary Cells to Favor Its Own Maternal Transmission.” mBio 15 (10): e01473–24. 10.1128/mbio.01473-24.

Montgomerie, Scott, Joseph A. Cruz, Savita Shrivastava, David Arndt, Mark Berjanskii, and David S. Wishart. 2008. “PROTEUS2: A Web Server for Comprehensive Protein Structure Prediction and Structure-Based Annotation.” Nucleic Acids Research 36 (suppl_2): W202–9. 10.1093/nar/gkn255.

Morin, X., R. Daneman, M. Zavortink, and W. Chia. 2001. A Protein Trap Strategy to Detect GFP-Tagged Proteins Expressed from Their Endogenous Loci in Drosophila. 98 (26): 15050–55.

Neuert, Helen, Petra Deing, Karin Krukkert, et al. 2020. “The Drosophila NCAM Homolog Fas2 Signals Independently of Adhesion.” Development 147 (2): dev181479.

Neville, Kathryn E., Tara M. Finegan, Nicholas Lowe, Philip M. Bellomio, Daxiang Na, and Dan T. Bergstralh. 2023. “The Drosophila Mitotic Spindle Orientation Machinery Requires Activation, Not Just Localization.” EMBO Reports 24 (3): e56074. 10.15252/embr.202256074.

Öztürk-Çolak, Arzu, Steven J. Marygold, Giulia Antonazzo, et al. 2024. “FlyBase: Updates to the Drosophila Genes and Genomes Database.” *Genetics*, February 1, iyad211. 10.1093/genetics/iyad211.

Rain, J. C., L. Selig, H. De Reuse, et al. 2001. “The Protein-Protein Interaction Map of Helicobacter Pylori.” Nature 409 (6817): 211–15. 10.1038/35051615.

Rechsteiner, Martin, and Scott W. Rogers. 1996. “PEST Sequences and Regulation by Proteolysis.” Trends in Biochemical Sciences 21 (7): 267–71. 10.1016/S0968-0004(96)10031-1.

Rees, Johanna S., Nick Lowe, Irina M. Armean, et al. 2011. “In Vivo Analysis of Proteomes and Interactomes Using Parallel Affinity Capture (iPAC) Coupled to Mass Spectrometry.” Molecular & Cellular Proteomics : MCP 10 (6): M110.002386.

Rubin, G. M., L. Hong, P. Brokstein, et al. 2000. “A Drosophila Complementary DNA Resource.” *Science (New York*, N.Y.) 287 (5461): 2222–24. 10.1126/science.287.5461.2222.

Sanes, Joshua R., and S. Lawrence Zipursky. 2020. “Synaptic Specificity, Recognition Molecules, and Assembly of Neural Circuits.” Cell 181 (6): 1434–35. 10.1016/j.cell.2020.05.046.

Schindelin, Johannes, Ignacio Arganda-Carreras, Erwin Frise, et al. 2012. “Fiji: An Open-Source Platform for Biological-Image Analysis.” Nature Methods 9 (7): 676–82.

Seeger, Mark, Guy Tear, Dolors Ferres-Marco, and Corey S. Goodman. 1993. “Mutations Affecting Growth Cone Guidance in Drosophila: Genes Necessary for Guidance toward or Away from the Midline.” Neuron 10 (3): 409–26. 10.1016/0896-6273(93)90330-T.

Sidisky, Jessica M., Alex Winters, Russell Caratenuto, and Daniel T. Babcock. 2024. “Synaptic Defects in a Drosophila Model of Muscular Dystrophy.” Frontiers in Cellular Neuroscience 18 (May): 1381112. 10.3389/fncel.2024.1381112.

Siegenthaler, Dominique, Eva-Maria Enneking, Eliza Moreno, and Jan Pielage. 2015. “L1CAM/Neuroglian Controls the Axon–Axon Interactions Establishing Layered and Lobular Mushroom Body Architecture.” Journal of Cell Biology 208 (7): 1003–18. 10.1083/jcb.201407131.

Slaidina, Maija, Selena Gupta, Torsten U. Banisch, and Ruth Lehmann. 2021. A Single-Cell Atlas Reveals Unanticipated Cell Type Complexity in Drosophila Ovaries. October 1. 10.1101/gr.274340.120.

Snow, Peter M., Allan J. Bieber, and Corey S. Goodman. 1989. “Fasciclin III: A Novel Homophilic Adhesion Molecule in Drosophila.” Cell 59 (2): 313–23.

Stranges, P. Benjamin, Mischa Machius, Michael J. Miley, Ashutosh Tripathy, and Brian Kuhlman. 2011. “Computational Design of a Symmetric Homodimer Using β-Strand Assembly.” Proceedings of the National Academy of Sciences 108 (51): 20562–67. 10.1073/pnas.1115124108.

Suter, Daniel M., and Paul Forscher. 2000. “Substrate–Cytoskeletal Coupling as a Mechanism for the Regulation of Growth Cone Motility and Guidance.” Journal of Neurobiology 44 (2): 97–113.

Szafranski, Przemyslaw, and Scott Goode. 2007. “Basolateral Junctions Are Sufficient to Suppress Epithelial Invasion during Drosophila Oogenesis.” Developmental Dynamics : An Official Publication of the American Association of Anatomists 236 (2): 364–73.

Taghli-Lamallem, Ouarda, Takeshi Akasaka, Grant Hogg, et al. 2008. “Dystrophin Deficiency in Drosophila Reduces Lifespan and Causes a Dilated Cardiomyopathy Phenotype.” Aging Cell 7 (2): 237–49. 10.1111/j.1474-9726.2008.00367.x.

Thomas, Ulrich, Eunjoon Kim, Sven Kuhlendahl, et al. 1997. “Synaptic Clustering of the Cell Adhesion Molecule Fasciclin II by Discs-Large and Its Role in the Regulation of Presynaptic Structure.” Neuron 19 (4): 787–99. 10.1016/S0896-6273(00)80961-7.

Ventura, Guilherme, Sofia Moreira, André Barros-Carvalho, Mariana Osswald, and Eurico Morais-de-Sá. 2020. “Lgl Cortical Dynamics Are Independent of Binding to the Scrib-Dlg Complex but Require Dlg-Dependent Restriction of aPKC.” Development 147 (15): dev186593. 10.1242/dev.186593.

Walsh, Gregory S., Paul K. Grant, John A. Morgan, and Cecilia B. Moens. 2011. “Planar Polarity Pathway and Nance-Horan Syndrome-like 1b Have Essential Cell-Autonomous Functions in Neuronal Migration.” Development 138 (14): 3033–42. 10.1242/dev.063842.

Wojcik, Jérôme, Ivo G. Boneca, and Pierre Legrain. 2002. “Prediction, Assessment and Validation of Protein Interaction Maps in Bacteria.” Journal of Molecular Biology 323 (4): 763–70. 10.1016/s0022-2836(02)01009-4.

Woods, D. F., C. Hough, D. Peel, G. Callaini, and P. J. Bryant. 1996. “Dlg Protein Is Required for Junction Structure, Cell Polarity, and Proliferation Control in Drosophila Epithelia.” Journal of Cell Biology 134 (6): 1469–82.

Zito, K., R. D. Fetter, C. S. Goodman, and E. Y. Isacoff. 1997. “Synaptic Clustering of Fascilin II and Shaker: Essential Targeting Sequences and Role of Dlg.” Neuron 19 (5): 1007–16. 10.1016/s0896-6273(00)80393-1.

Zito, Karen, Richard D. Fetter, Corey S. Goodman, and Ehud Y. Isacoff. 1997. “Synaptic Clustering of Fasciclin II and Shaker: Essential Targeting Sequences and Role of Dlg.” Neuron 19 (5): 1007–16. 10.1016/S0896-6273(00)80393-1.

