## Supplemental Files for "Fasciclin 2 Cooperates with Discs Large to Maintain Epithelial Architecture"

### Supplemental Figure 1 - Finegan & Linhoff *et al.*

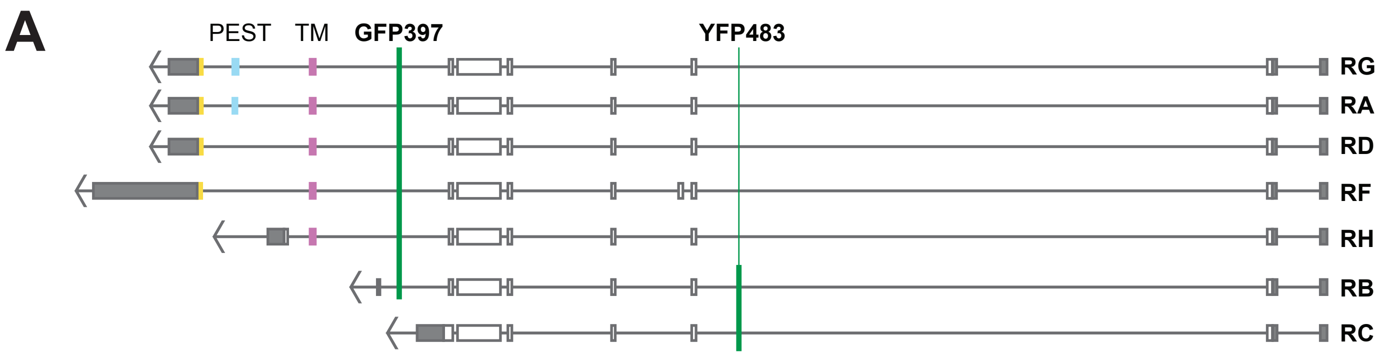

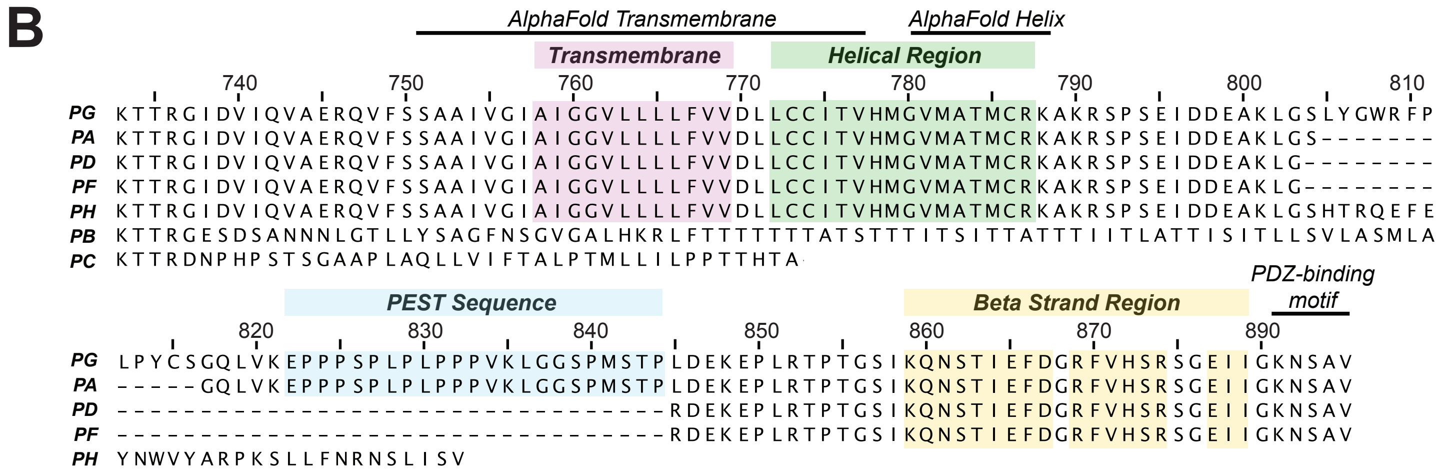

### Supplemental Figure 2 - Finegan & Linhoff *et al.*

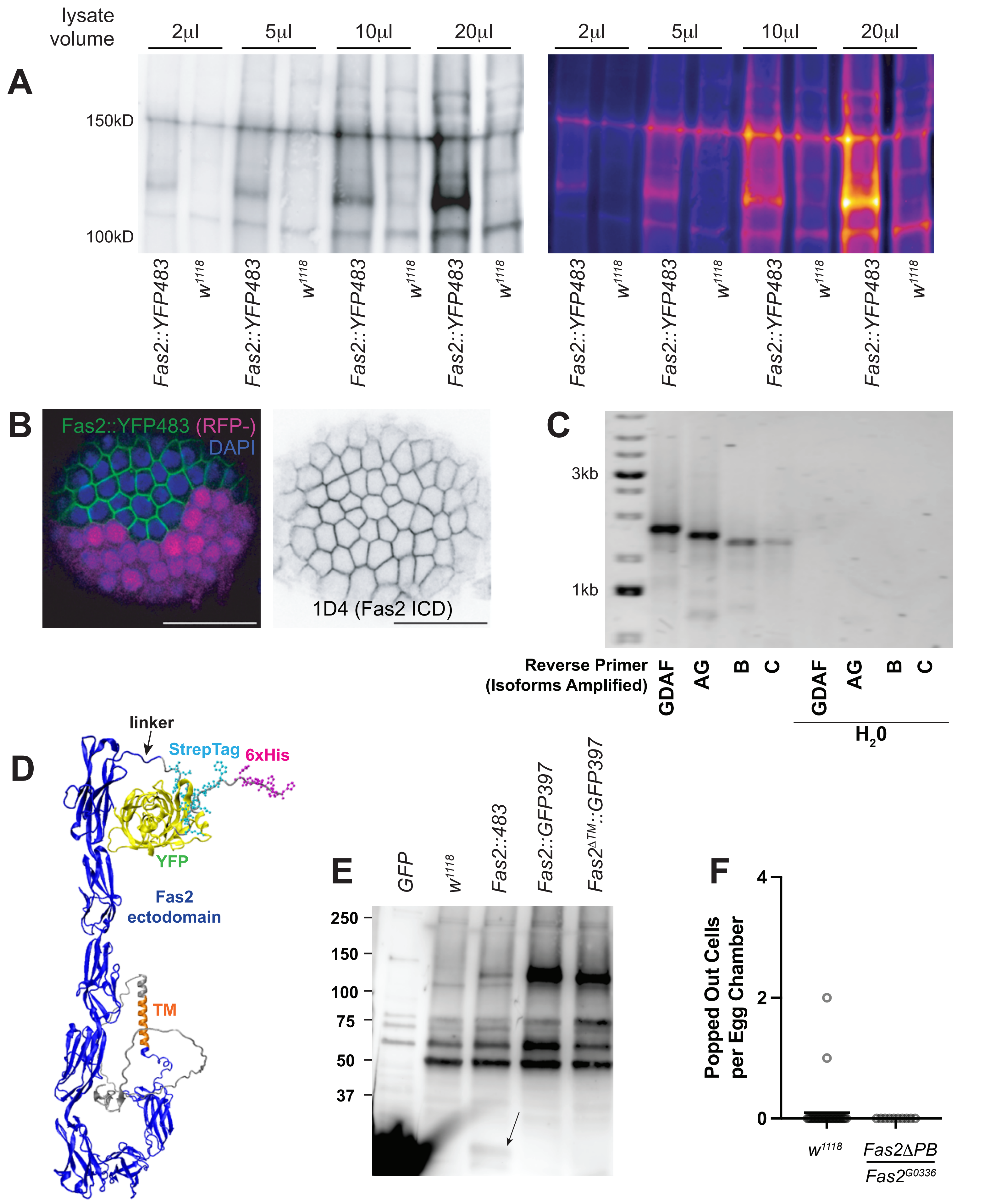

### Supplemental Figure 3 - Finegan & Linhoff *et al.*

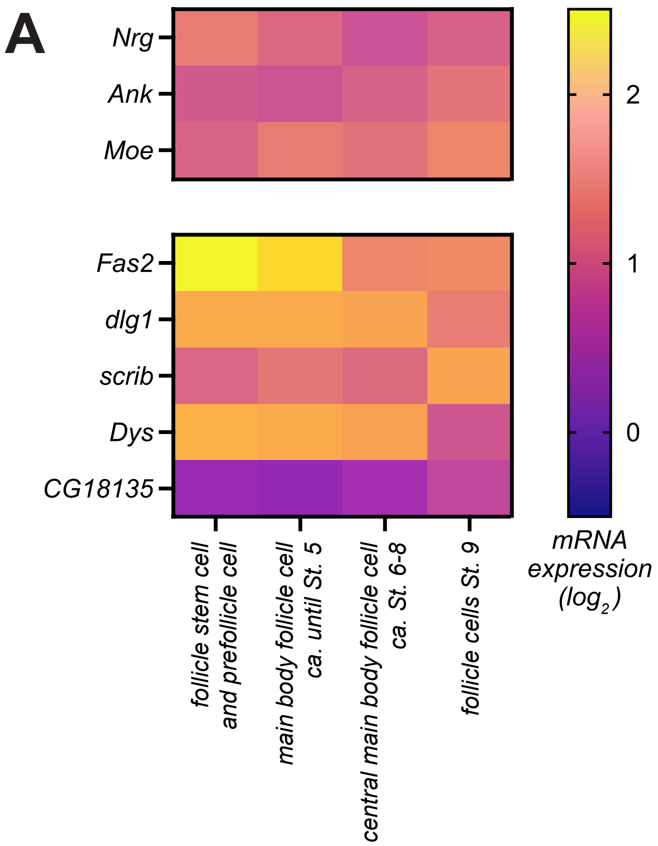

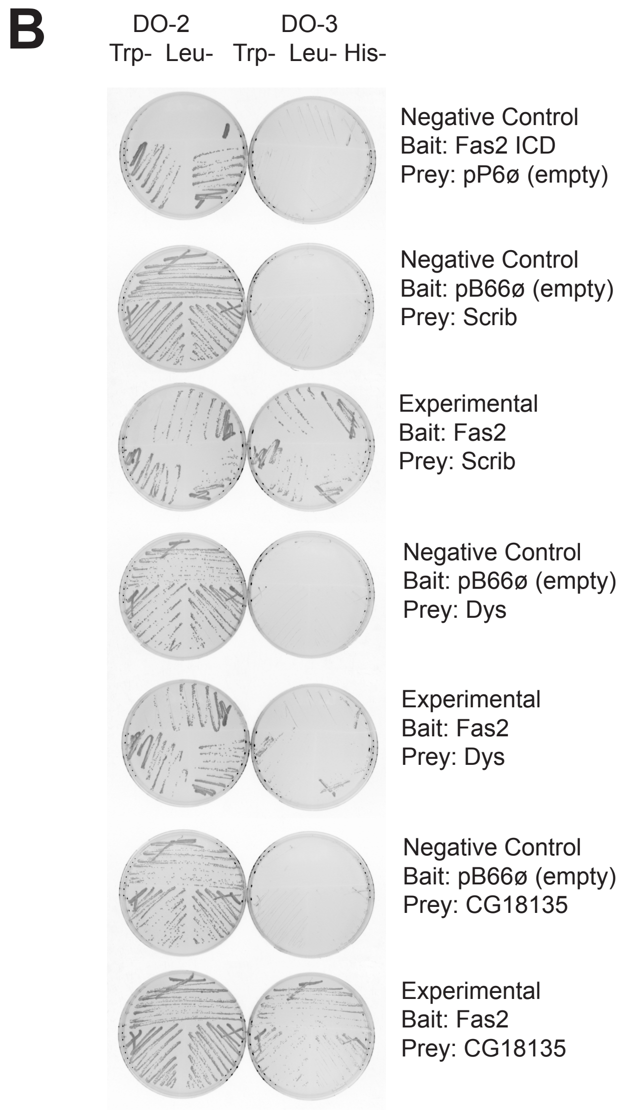

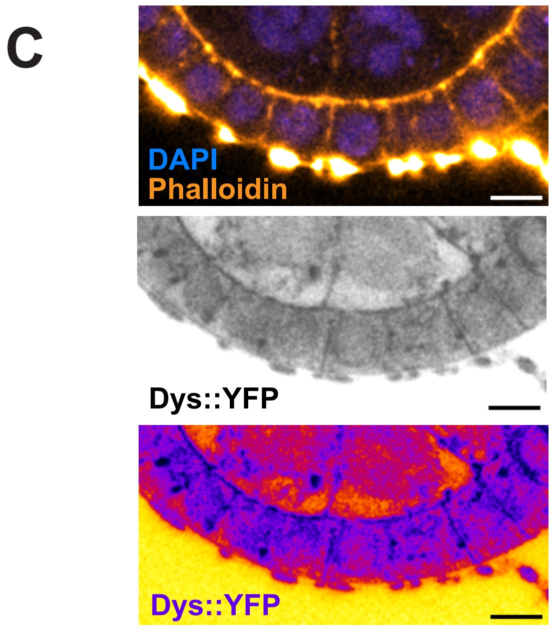

pB66: Gal4 DNA-Binding Domain (DBD) vector, i.e. bait vector (DBD-bait)

pB66ø: empty pB66 vector

pP6: Gal4 Activation Domain (AD) vector, i.e. prey vector (AD-prey).

pP6ø: empty pP6 vector

DO-2: selective medium without tryptophan and leucine

DO-3: selective medium without tryptophan, leucine and histidine

**Supplemental Figure 4 - Finegan & Linhoff *et al.***

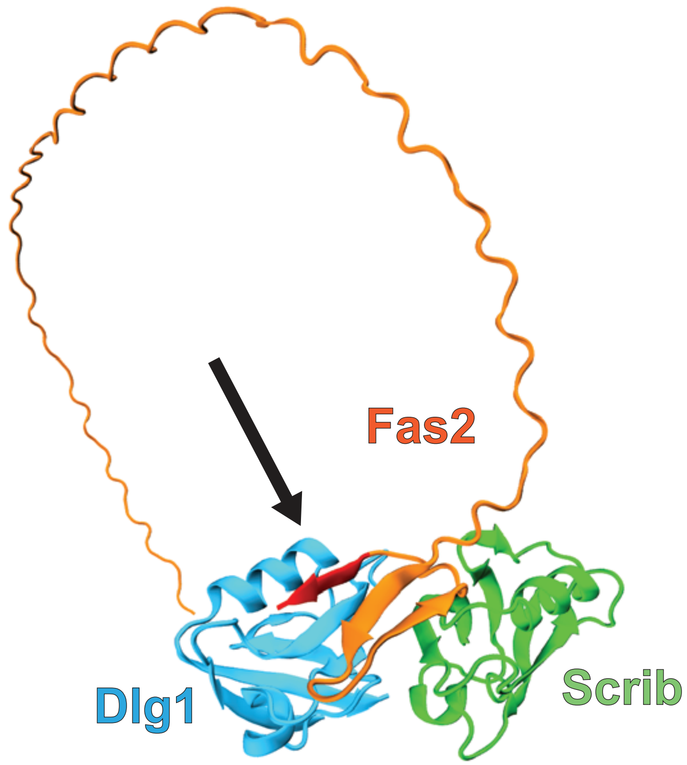

### Supplemental Figure 5 - Finegan & Linhoff *et al.*

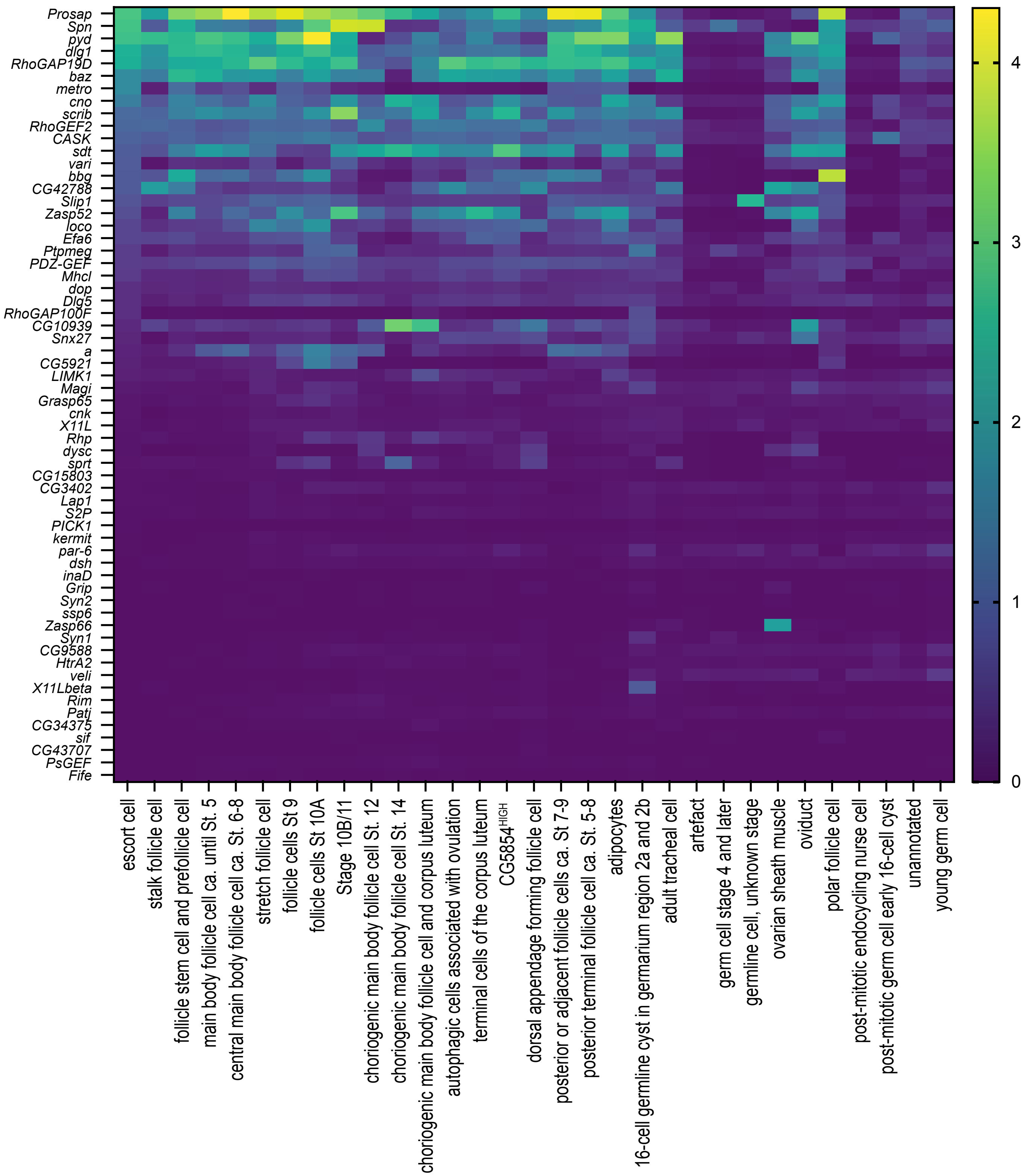

### Supplemental Figure 6 - Finegan & Linhoff *et al.*

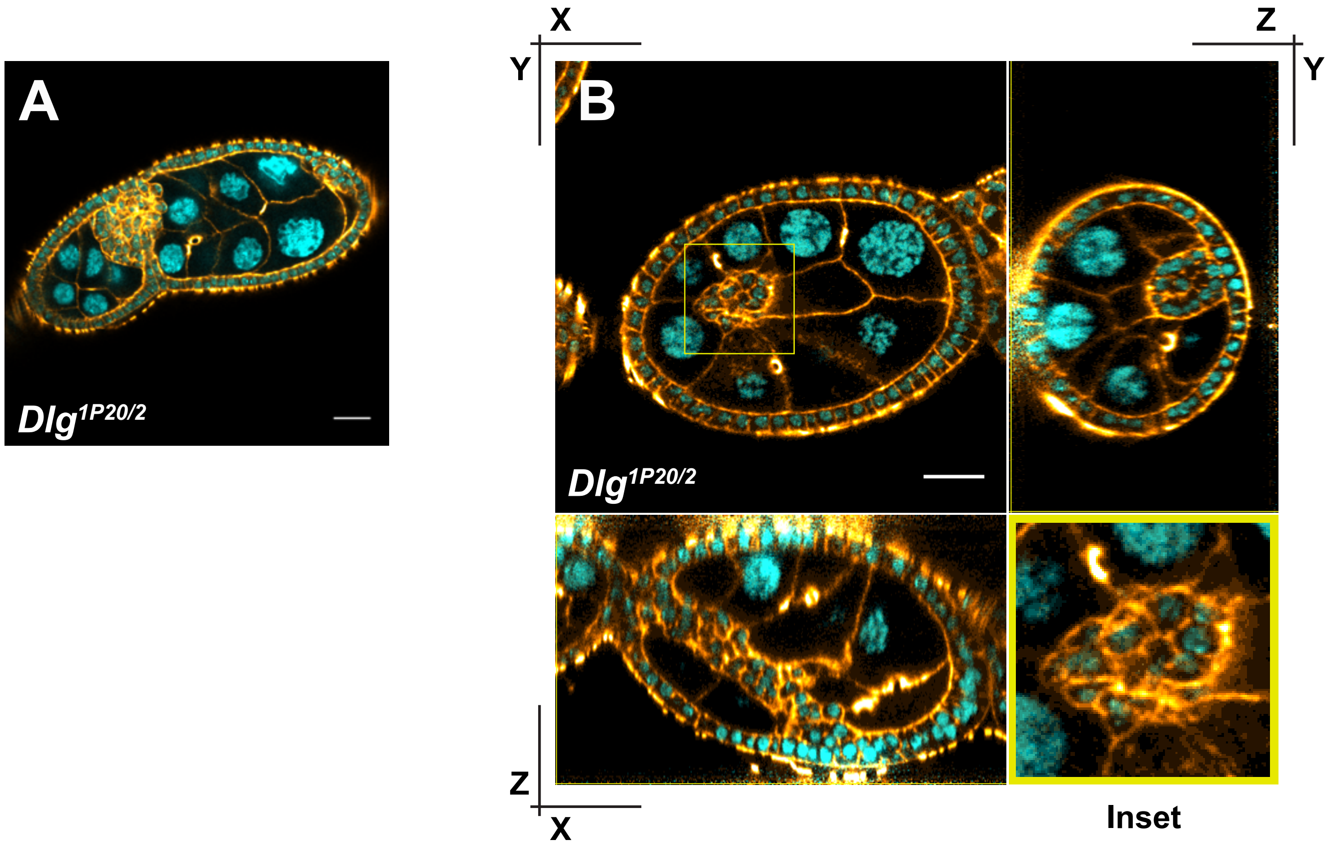

### Supplemental Figure 7 - Finegan & Linhoff *et al.*

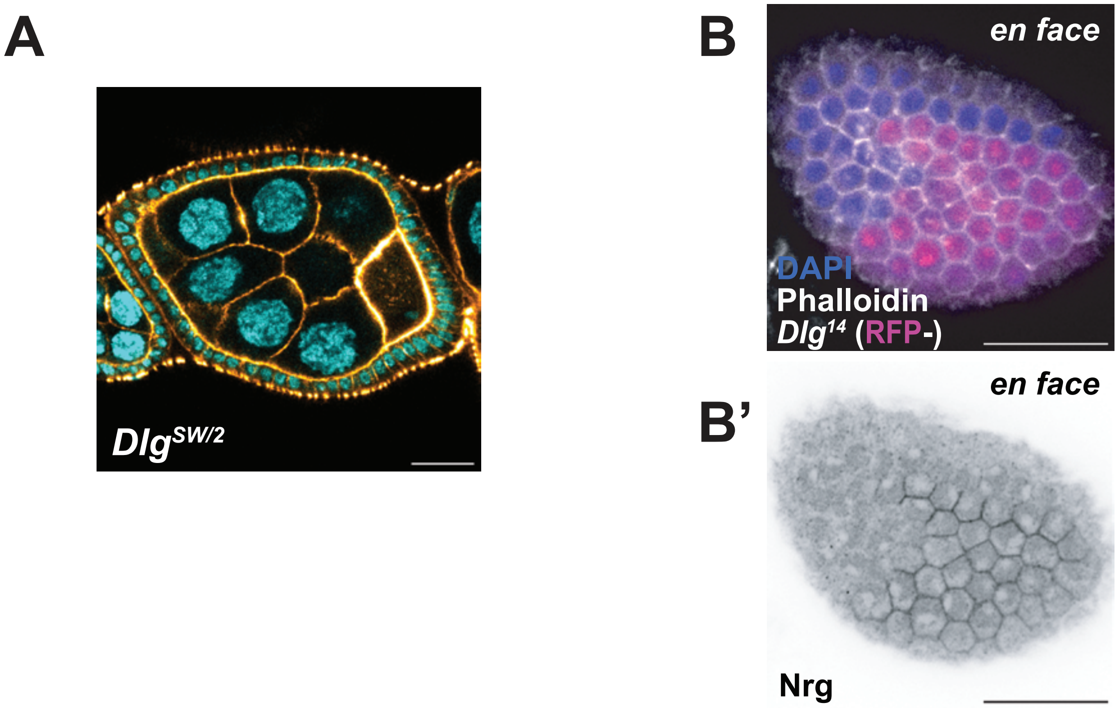

**Table 1 – Finegan & Linhoff et al.**

| Mutant Allele or Transgene | Citation | Stock Number or Lab of Origin |
|---|---|---|
| $Fas2^{EB112}$ | (Grenningloh et al. 1991) | Riechmann lab |
| $Fas2^{G0336}$ | (Peter et al. 2002) | St Johnston lab |
| $Fas2^{\Delta TM::GFP397}$ | (Neuert et al. 2020) | Klämbt lab |
| $Fas2^{\Delta PB::GFP397}$ | (Neuert et al. 2020) | Klämbt lab |
| $Fas2^{\Delta PC}$ | (Neuert et al. 2020) | Klämbt lab |
| *Fas2::YFP483* $^{(CPTI000483)}$ | (Lowe et al. 2014) | KYOTO 115-501 |
| *Fas2::GFP397* | (Morin et al. 2001) | BDSC 602883 |
| UAS-Fas2-RNAi TRiP.HMS01098 | (Perkins et al. 2015) | BDSC 34084 |
| $Nrg^{14}$ (aka $Nrg^{l4}$) | (Hortsch et al. 1998) | BDSC 5708 |
| $Nrg^{DFERM\text{-}binding}$ | (Siegenthaler et al. 2015) | Pielage lab |
| UAS-Nrg-RNAi TRiP.HMS01638 | (Perkins et al. 2015) | BDSC 37496 |
| $Dlg^{14}$ (aka $Dlg1^{m52}$) | (Woods and Bryant 1991) | BDSC 36283 |
| $Dlg^{1P20}$ (aka $Dlg1^{18}$) | (Woods and Bryant 1991) | BDSC 36279 |
| $Dlg^{SW}$ | (Woods and Bryant 1991) | Bellaïche lab |
| $Dlg^{2}$ | (Perrimon 1988) | BDSC 36278 |
| UAS-Dlg-RNAi TRiP.HMS00014 | (Perkins et al. 2015) | BDSC 33620 |
| Dp(1;3)DC238 | (Venken et al. 2010) | BDSC 30357 |
| UAS-Scrib-RNAi TRiP.HMS01490 | (Ni et al. 2011) | BDSC 35748 |
| *Df(3R)Exel6184* | (Taghli-Lamallem et al. 2008) | BDSC 7663 |
| $Dys^{E17}$ | | BDSC 63047 |

| | | |
|---|---|---|
| UAS-Dys-RNAi TRiP.JF01118 | (Perkins et al. 2015) | BDSC 31553 |
| *Dys*$^{CPTI002256}$ | (Lowe et al. 2014) | KYOTO 115-283 |
| Traffic jam-GAL4 | (Olivieri et al. 2010) | Lehmann lab |
| GR1-GAL4 | (Tran and Berg 2003) | BDSC 36287 |
| UAS-Inscuteable | (Kraut et al. 1996) | Knoblich lab |
| Ubi-mRFP.nls, hsFLP, FRT19A | (Chou and Perrimon 1996) | BDSC 31418 |

***Drosophila* References**

Chou, T. B., and N. Perrimon. 1996. “The Autosomal FLP-DFS Technique for Generating Germline Mosaics in Drosophila Melanogaster.” *Genetics* 144 (4): 1673–79. https://doi.org/10.1093/genetics/144.4.1673.

Grenningloh, G., E. J. Rehm, and C. S. Goodman. 1991. “Genetic Analysis of Growth Cone Guidance in Drosophila: Fasciclin II Functions as a Neuronal Recognition Molecule.” *Cell* 67 (1): 45–57.

Hortsch, M., D. Homer, J. D. Malhotra, et al. 1998. “Structural Requirements for Outside-in and inside-out Signaling by Drosophila Neuroglian, a Member of the L1 Family of Cell Adhesion Molecules.” *Journal of Cell Biology* 142 (1): 251–61.

Kraut, R., W. Chia, L. Y. Jan, Y. N. Jan, and J. A. Knoblich. 1996. “Role of Inscuteable in Orienting Asymmetric Cell Divisions in Drosophila.” *Nature* 383 (6595): 50–55.

Lowe, Nick, Johanna S. Rees, John Roote, et al. 2014. “Analysis of the Expression Patterns, Subcellular Localisations and Interaction Partners of Drosophila Proteins Using a pigP Protein Trap Library.” *Development* 141 (20): 3994–4005.

Morin, X., R. Daneman, M. Zavortink, and W. Chia. 2001. *A Protein Trap Strategy to Detect GFP-Tagged Proteins Expressed from Their Endogenous Loci in Drosophila.* 98 (26): 15050–55.

Neuert, Helen, Petra Deing, Karin Krukkert, et al. 2020. “The Drosophila NCAM Homolog Fas2 Signals Independently of Adhesion.” *Development* 147 (2): dev181479.

Ni, Jian-Quan, Rui Zhou, Benjamin Czech, et al. 2011. “A Genome-Scale shRNA Resource for Transgenic RNAi in Drosophila.” *Nature Methods* 8 (5): 405–7. https://doi.org/10.1038/nmeth.1592.

Olivieri, Daniel, Martina M. Sykora, Ravi Sachidanandam, Karl Mechtler, and Julius Brennecke. 2010. “An in Vivo RNAi Assay Identifies Major Genetic and Cellular Requirements for Primary piRNA Biogenesis in Drosophila.” *The EMBO Journal* 29 (19): 3301–17.

Perkins, Lizabeth A., Laura Holderbaum, Rong Tao, et al. 2015. “The Transgenic RNAi Project at Harvard Medical School: Resources and Validation.” *Genetics* 201 (3): 843–52.

Perrimon, Norbert. 1988. “The Maternal Effect of *Lethal(1)Discs-Large-1:* A Recessive Oncogene of *Drosophila Melanogaster*.” *Developmental Biology* 127 (2): 392–407. https://doi.org/10.1016/0012-1606(88)90326-0.

Peter, Annette, Petra Schöttler, Meike Werner, et al. 2002. “Mapping and Identification of Essential Gene Functions on the X Chromosome of Drosophila.” *The EMBO Reports* 3 (1): 34–38. https://doi.org/10.1093/embo-reports/kvf012.

Siegenthaler, Dominique, Eva-Maria Enneking, Eliza Moreno, and Jan Pielage. 2015. “L1CAM/Neuroglian Controls the Axon–Axon Interactions Establishing Layered and Lobular Mushroom Body Architecture.” *Journal of Cell Biology* 208 (7): 1003–18.

Taghli-Lamallem, Ouarda, Takeshi Akasaka, Grant Hogg, et al. 2008. “Dystrophin Deficiency in Drosophila Reduces Lifespan and Causes a Dilated Cardiomyopathy Phenotype.” *Aging Cell* 7 (2): 237–49. https://doi.org/10.1111/j.1474-9726.2008.00367.x.

Tran, David H., and Celeste A. Berg. 2003. “Bullwinkle and Shark Regulate Dorsal-Appendage Morphogenesis in Drosophila Oogenesis.” *Development* 130 (25): 6273–82. https://doi.org/10.1242/dev.00854.

Venken, Koen J. T., Ellen Popodi, Stacy L. Holtzman, et al. 2010. “A Molecularly Defined Duplication Set for the X Chromosome of Drosophila Melanogaster.” *Genetics* 186 (4): 1111–25. https://doi.org/10.1534/genetics.110.121285.

Woods, D. F., and P. J. Bryant. 1991. “The Discs-Large Tumor Suppressor Gene of Drosophila Encodes a Guanylate Kinase Homolog Localized at Septate Junctions.” *Cell* 66 (3): 451–64.

**Table 2 – Finegan & Linhoff *et al.***

| Reagent | Source |
|---|---|
| **Imaging Reagents** | |
| mouse anti-Fas2 (1D4) | DSHB |
| mouse anti-Fas3 | DSHB |
| rabbit anti-Nrg | Our lab (Finegan et al. (2024) |
| rabbit anti-GFP | Proteintech: 50430-2-AP Lot# 00179327 |
| mouse anti-Dlg (4F3) | DSHB 2/18/16 |
| mouse anti-Fas2 (1D4) | DSHB 6/9/16 |
| Conjugated secondary antibodies | ThermoFisher Scientific |
| Fluorescein Phalloidin | Invitrogen: Lot# 2138394 |
| Alexa fluor 633 Phalloidin | Invitrogen: Lot# 2274768 |
| Vectashield with DAPI | Vector Labs: Lot# ZK03217 |
| **Western Blotting** | |
| SuperSignal West Pico PLUS chemiluminescent substrate | Thermo Scientific |
| Pierce protease inhibitor | Thermo Fisher: Lot# SB2334961 |
| Amido Black | Sigma: Lot# SLBR9882V |
| Precision Plus Protein Kaleidoscope prestained protein standards | Bio-Rad |
| Protease Inhibitors | Roche Applied Science |
| 7.5% Mini-PROTEAN TGX gels | Bio-Rad |
| **RT-PCR** | |
| Zymo Research Quick-RNA MiniPrep Kit | Zymo Research Cat.: Lot# 256847 |
| Invitrogen SuperScript II First-Strand Synthesis System | ThermoFisher Scientific: Lot#1824456 |
| NEB Taq 5X Master Mix | New England Biolabs: Lot# 10215729 |
| Invitrogen SYBR Safe DNA Gel Stain | ThermoFisher Scientific: Lot# 3253469 |
| Thermo Scientific GeneRuler 1KB DNA ladder | ThermoFisher Scientific |
| NEB 6X Gel Loading Dye | New England Biolabs |
| **Fly Food** | |
| Apex Tegosept | Genesee Scientific |
| Apex Propionic Acid | Genesee Scientific |
| Nutrifly Bloomington Formulation | Genesee Scientific |

**Table X**
RT-PCR primers

| | | |
|---|---|---|
| Forward Primer | | |
| CPTI Tag | | AAGCGCGATCACATGGTCC |
| Reverse Primers | Isoforms Amplified | |
| Fas2 A_rev | GDAF | GCCGAATTCTTCCCGATTAT |
| Fas2 A_PEST_rev | AG | CGGTGGCTCCTTTACCAG |
| Fas2 B_rev | B | CAAAATCAGCAGCATTGTCG |
| Fas2 C_rev | C | TGTGGCTGTTGTTGTTGTTG |

**Table 4** - Nrg (aa 1155-1239) vs Drosophila Ovaries

Clone Name;Type Seq;Contig(s) Name;Gene Name (Best Match);GenBank ID (NCBI);Gene ID (NCBI);Other ID when available;Global PBS;Additional Gene Notes;Start;Stop;Frame;Sense;UTR Inclusi
DMOV_RP1_hgx6470v1_pB66_A-99;3p ;162046772;Drosophila melanogaster - Ank;https://www.ncbi.nlm.nih.gov/nuccore/NM_001169348.2;https://www.ncbi.nlm.nih.gov/gene/?term=43770;FBgn00
DMOV_RP1_hgx6470v1_pB66_A-107;3p ;162046772;Drosophila melanogaster - Ank;https://www.ncbi.nlm.nih.gov/nuccore/NM_001169348.2;https://www.ncbi.nlm.nih.gov/gene/?term=43770;FBgn0
DMOV_RP1_hgx6470v1_pB66_A-137;3p ;162046772;Drosophila melanogaster - Ank;https://www.ncbi.nlm.nih.gov/nuccore/NM_001169348.2;https://www.ncbi.nlm.nih.gov/gene/?term=43770;FBgn0
DMOV_RP1_hgx6470v1_pB66_A-138;3p ;162046772;Drosophila melanogaster - Ank;https://www.ncbi.nlm.nih.gov/nuccore/NM_001169348.2;https://www.ncbi.nlm.nih.gov/gene/?term=43770;FBgn0
DMOV_RP1_hgx6470v1_pB66_A-143;3p ;162046772;Drosophila melanogaster - Ank;https://www.ncbi.nlm.nih.gov/nuccore/NM_001169348.2;https://www.ncbi.nlm.nih.gov/gene/?term=43770;FBgn0
DMOV_RP1_hgx6470v1_pB66_A-148;3p ;162046772;Drosophila melanogaster - Ank;https://www.ncbi.nlm.nih.gov/nuccore/NM_001169348.2;https://www.ncbi.nlm.nih.gov/gene/?term=43770;FBgn0
DMOV_RP1_hgx6470v1_pB66_A-150;3p ;162046772;Drosophila melanogaster - Ank;https://www.ncbi.nlm.nih.gov/nuccore/NM_001169348.2;https://www.ncbi.nlm.nih.gov/gene/?term=43770;FBgn0
DMOV_RP1_hgx6470v1_pB66_A-178;3p ;162046772;Drosophila melanogaster - Ank;https://www.ncbi.nlm.nih.gov/nuccore/NM_001169348.2;https://www.ncbi.nlm.nih.gov/gene/?term=43770;FBgn0
DMOV_RP1_hgx6470v1_pB66_A-22;3p ;162046772;Drosophila melanogaster - Ank;https://www.ncbi.nlm.nih.gov/nuccore/NM_001169348.2;https://www.ncbi.nlm.nih.gov/gene/?term=43770;FBgn00
DMOV_RP1_hgx6470v1_pB66_A-24;3p ;162046772;Drosophila melanogaster - Ank;https://www.ncbi.nlm.nih.gov/nuccore/NM_001169348.2;https://www.ncbi.nlm.nih.gov/gene/?term=43770;FBgn00
DMOV_RP1_hgx6470v1_pB66_A-41;3p ;162046772;Drosophila melanogaster - Ank;https://www.ncbi.nlm.nih.gov/nuccore/NM_001169348.2;https://www.ncbi.nlm.nih.gov/gene/?term=43770;FBgn00
DMOV_RP1_hgx6470v1_pB66_A-50;3p ;162046772;Drosophila melanogaster - Ank;https://www.ncbi.nlm.nih.gov/nuccore/NM_001169348.2;https://www.ncbi.nlm.nih.gov/gene/?term=43770;FBgn00
DMOV_RP1_hgx6470v1_pB66_A-63;3p ;162046772;Drosophila melanogaster - Ank;https://www.ncbi.nlm.nih.gov/nuccore/NM_001169348.2;https://www.ncbi.nlm.nih.gov/gene/?term=43770;FBgn00
DMOV_RP1_hgx6470v1_pB66_A-91;3p ;162046772;Drosophila melanogaster - Ank;https://www.ncbi.nlm.nih.gov/nuccore/NM_001169348.2;https://www.ncbi.nlm.nih.gov/gene/?term=43770;FBgn00
DMOV_RP1_hgx6470v1_pB66_A-31;5p ;162046772;Drosophila melanogaster - Ank;https://www.ncbi.nlm.nih.gov/nuccore/NM_001169348.2;https://www.ncbi.nlm.nih.gov/gene/?term=43770;FBgn00
DMOV_RP1_hgx6470v1_pB66_A-55;5p 3p ;162046772;Drosophila melanogaster - Ank;https://www.ncbi.nlm.nih.gov/nuccore/NM_001169348.2;https://www.ncbi.nlm.nih.gov/gene/?term=43770;FBg
DMOV_RP1_hgx6470v1_pB66_A-100;5p ;162046772;Drosophila melanogaster - Ank;https://www.ncbi.nlm.nih.gov/nuccore/NM_001169348.2;https://www.ncbi.nlm.nih.gov/gene/?term=43770;FBgn0
DMOV_RP1_hgx6470v1_pB66_A-140;5p ;162046772;Drosophila melanogaster - Ank;https://www.ncbi.nlm.nih.gov/nuccore/NM_001169348.2;https://www.ncbi.nlm.nih.gov/gene/?term=43770;FBgn0
DMOV_RP1_hgx6470v1_pB66_A-139;5p 3p ;162046772;Drosophila melanogaster - Ank;https://www.ncbi.nlm.nih.gov/nuccore/NM_001169348.2;https://www.ncbi.nlm.nih.gov/gene/?term=43770;FB
DMOV_RP1_hgx6470v1_pB66_A-20;5p 3p ;162046772;Drosophila melanogaster - Ank;https://www.ncbi.nlm.nih.gov/nuccore/NM_001169348.2;https://www.ncbi.nlm.nih.gov/gene/?term=43770;FBg
DMOV_RP1_hgx6470v1_pB66_A-23;5p 3p ;162046772;Drosophila melanogaster - Ank;https://www.ncbi.nlm.nih.gov/nuccore/NM_001169348.2;https://www.ncbi.nlm.nih.gov/gene/?term=43770;FBg
DMOV_RP1_hgx6470v1_pB66_A-7;5p 3p ;162046772;Drosophila melanogaster - Ank;https://www.ncbi.nlm.nih.gov/nuccore/NM_001169348.2;https://www.ncbi.nlm.nih.gov/gene/?term=43770;FBgn
DMOV_RP1_hgx6470v1_pB66_A-124;5p 3p ;162046772;Drosophila melanogaster - Ank;https://www.ncbi.nlm.nih.gov/nuccore/NM_001169348.2;https://www.ncbi.nlm.nih.gov/gene/?term=43770;FB
DMOV_RP1_hgx6470v1_pB66_A-169;5p 3p ;162046772;Drosophila melanogaster - Ank;https://www.ncbi.nlm.nih.gov/nuccore/NM_001169348.2;https://www.ncbi.nlm.nih.gov/gene/?term=43770;FB
DMOV_RP1_hgx6470v1_pB66_A-177;5p 3p ;162046772;Drosophila melanogaster - Ank;https://www.ncbi.nlm.nih.gov/nuccore/NM_001169348.2;https://www.ncbi.nlm.nih.gov/gene/?term=43770;FB
DMOV_RP1_hgx6470v1_pB66_A-125;5p 3p ;162046772;Drosophila melanogaster - Ank;https://www.ncbi.nlm.nih.gov/nuccore/NM_001169348.2;https://www.ncbi.nlm.nih.gov/gene/?term=43770;FB
DMOV_RP1_hgx6470v1_pB66_A-80;5p 3p ;162046772;Drosophila melanogaster - Ank;https://www.ncbi.nlm.nih.gov/nuccore/NM_001169348.2;https://www.ncbi.nlm.nih.gov/gene/?term=43770;FBg
DMOV_RP1_hgx6470v1_pB66_A-53;5p 3p ;162046772;Drosophila melanogaster - Ank;https://www.ncbi.nlm.nih.gov/nuccore/NM_001169348.2;https://www.ncbi.nlm.nih.gov/gene/?term=43770;FBg
DMOV_RP1_hgx6470v1_pB66_A-164;5p 3p ;162046772;Drosophila melanogaster - Ank;https://www.ncbi.nlm.nih.gov/nuccore/NM_001169348.2;https://www.ncbi.nlm.nih.gov/gene/?term=43770;FB
DMOV_RP1_hgx6470v1_pB66_A-101;5p 3p ;162046772;Drosophila melanogaster - Ank;https://www.ncbi.nlm.nih.gov/nuccore/NM_001169348.2;https://www.ncbi.nlm.nih.gov/gene/?term=43770;FB
DMOV_RP1_hgx6470v1_pB66_A-149;5p 3p ;162046772;Drosophila melanogaster - Ank;https://www.ncbi.nlm.nih.gov/nuccore/NM_001169348.2;https://www.ncbi.nlm.nih.gov/gene/?term=43770;FB
DMOV_RP1_hgx6470v1_pB66_A-89;5p 3p ;162046772;Drosophila melanogaster - Ank;https://www.ncbi.nlm.nih.gov/nuccore/NM_001169348.2;https://www.ncbi.nlm.nih.gov/gene/?term=43770;FBg
DMOV_RP1_hgx6470v1_pB66_A-90;5p 3p ;162046772;Drosophila melanogaster - Ank;https://www.ncbi.nlm.nih.gov/nuccore/NM_001169348.2;https://www.ncbi.nlm.nih.gov/gene/?term=43770;FBg
DMOV_RP1_hgx6470v1_pB66_A-84;5p 3p ;162046772;Drosophila melanogaster - Ank;https://www.ncbi.nlm.nih.gov/nuccore/NM_001169348.2;https://www.ncbi.nlm.nih.gov/gene/?term=43770;FBg
DMOV_RP1_hgx6470v1_pB66_A-33;5p 3p ;162046772;Drosophila melanogaster - Ank;https://www.ncbi.nlm.nih.gov/nuccore/NM_001169348.2;https://www.ncbi.nlm.nih.gov/gene/?term=43770;FBg
DMOV_RP1_hgx6470v1_pB66_A-70;5p 3p ;162046772;Drosophila melanogaster - Ank;https://www.ncbi.nlm.nih.gov/nuccore/NM_001169348.2;https://www.ncbi.nlm.nih.gov/gene/?term=43770;FBg
DMOV_RP1_hgx6470v1_pB66_A-71;5p 3p ;162046772;Drosophila melanogaster - Ank;https://www.ncbi.nlm.nih.gov/nuccore/NM_001169348.2;https://www.ncbi.nlm.nih.gov/gene/?term=43770;FBg
DMOV_RP1_hgx6470v1_pB66_A-181;5p 3p ;162046772;Drosophila melanogaster - Ank;https://www.ncbi.nlm.nih.gov/nuccore/NM_001169348.2;https://www.ncbi.nlm.nih.gov/gene/?term=43770;FB
DMOV_RP1_hgx6470v1_pB66_A-119;5p ;162046772;Drosophila melanogaster - Ank;https://www.ncbi.nlm.nih.gov/nuccore/NM_001169348.2;https://www.ncbi.nlm.nih.gov/gene/?term=43770;FBgn0
DMOV_RP1_hgx6470v1_pB66_A-120;5p 3p ;162046772;Drosophila melanogaster - Ank;https://www.ncbi.nlm.nih.gov/nuccore/NM_001169348.2;https://www.ncbi.nlm.nih.gov/gene/?term=43770;FB

DMOV_RP1_hgx6470v1_pB66_A-162;5p 3p ;162046772;Drosophila melanogaster - Ank;https://www.ncbi.nlm.nih.gov/nuccore/NM_001169348.2;https://www.ncbi.nlm.nih.gov/gene/?term=43770;FBg
DMOV_RP1_hgx6470v1_pB66_A-92;5p 3p ;162046772;Drosophila melanogaster - Ank;https://www.ncbi.nlm.nih.gov/nuccore/NM_001169348.2;https://www.ncbi.nlm.nih.gov/gene/?term=43770;FBgn
DMOV_RP1_hgx6470v1_pB66_A-95;5p 3p ;162046772;Drosophila melanogaster - Ank;https://www.ncbi.nlm.nih.gov/nuccore/NM_001169348.2;https://www.ncbi.nlm.nih.gov/gene/?term=43770;FBgn
DMOV_RP1_hgx6470v1_pB66_A-122;5p ;162046772;Drosophila melanogaster - Ank;https://www.ncbi.nlm.nih.gov/nuccore/NM_001169348.2;https://www.ncbi.nlm.nih.gov/gene/?term=43770;FBgn0
DMOV_RP1_hgx6470v1_pB66_A-2;5p 3p ;162046772;Drosophila melanogaster - Ank;https://www.ncbi.nlm.nih.gov/nuccore/NM_001169348.2;https://www.ncbi.nlm.nih.gov/gene/?term=43770;FBgn0
DMOV_RP1_hgx6470v1_pB66_A-113;5p 3p ;162046772;Drosophila melanogaster - Ank;https://www.ncbi.nlm.nih.gov/nuccore/NM_001169348.2;https://www.ncbi.nlm.nih.gov/gene/?term=43770;FBg
DMOV_RP1_hgx6470v1_pB66_A-88;5p 3p ;162046772;Drosophila melanogaster - Ank;https://www.ncbi.nlm.nih.gov/nuccore/NM_001169348.2;https://www.ncbi.nlm.nih.gov/gene/?term=43770;FBgn
DMOV_RP1_hgx6470v1_pB66_A-39;5p 3p ;162046772;Drosophila melanogaster - Ank;https://www.ncbi.nlm.nih.gov/nuccore/NM_001169348.2;https://www.ncbi.nlm.nih.gov/gene/?term=43770;FBgn
DMOV_RP1_hgx6470v1_pB66_A-21;5p 3p ;162046772;Drosophila melanogaster - Ank;https://www.ncbi.nlm.nih.gov/nuccore/NM_001169348.2;https://www.ncbi.nlm.nih.gov/gene/?term=43770;FBgn
DMOV_RP1_hgx6470v1_pB66_A-52;5p 3p ;162046772;Drosophila melanogaster - Ank;https://www.ncbi.nlm.nih.gov/nuccore/NM_001169348.2;https://www.ncbi.nlm.nih.gov/gene/?term=43770;FBgn
DMOV_RP1_hgx6470v1_pB66_A-58;5p 3p ;162046772;Drosophila melanogaster - Ank;https://www.ncbi.nlm.nih.gov/nuccore/NM_001169348.2;https://www.ncbi.nlm.nih.gov/gene/?term=43770;FBgn
DMOV_RP1_hgx6470v1_pB66_A-98;5p 3p ;162046772;Drosophila melanogaster - Ank;https://www.ncbi.nlm.nih.gov/nuccore/NM_001169348.2;https://www.ncbi.nlm.nih.gov/gene/?term=43770;FBgn
DMOV_RP1_hgx6470v1_pB66_A-10;5p 3p ;162046772;Drosophila melanogaster - Ank;https://www.ncbi.nlm.nih.gov/nuccore/NM_001169348.2;https://www.ncbi.nlm.nih.gov/gene/?term=43770;FBgn
DMOV_RP1_hgx6470v1_pB66_A-15;5p 3p ;162046772;Drosophila melanogaster - Ank;https://www.ncbi.nlm.nih.gov/nuccore/NM_001169348.2;https://www.ncbi.nlm.nih.gov/gene/?term=43770;FBgn
DMOV_RP1_hgx6470v1_pB66_A-123;5p 3p ;162046772;Drosophila melanogaster - Ank;https://www.ncbi.nlm.nih.gov/nuccore/NM_001169348.2;https://www.ncbi.nlm.nih.gov/gene/?term=43770;FBg
DMOV_RP1_hgx6470v1_pB66_A-127;5p 3p ;162046772;Drosophila melanogaster - Ank;https://www.ncbi.nlm.nih.gov/nuccore/NM_001169348.2;https://www.ncbi.nlm.nih.gov/gene/?term=43770;FBg
DMOV_RP1_hgx6470v1_pB66_A-112;5p 3p ;162046772;Drosophila melanogaster - Ank;https://www.ncbi.nlm.nih.gov/nuccore/NM_001169348.2;https://www.ncbi.nlm.nih.gov/gene/?term=43770;FBg
DMOV_RP1_hgx6470v1_pB66_A-25;5p ;162046772;Drosophila melanogaster - Ank;https://www.ncbi.nlm.nih.gov/nuccore/NM_001169348.2;https://www.ncbi.nlm.nih.gov/gene/?term=43770;FBgn00
DMOV_RP1_hgx6470v1_pB66_A-109;5p 3p ;162046772;Drosophila melanogaster - Ank;https://www.ncbi.nlm.nih.gov/nuccore/NM_001169348.2;https://www.ncbi.nlm.nih.gov/gene/?term=43770;FBg
DMOV_RP1_hgx6470v1_pB66_A-97;5p 3p ;162046772;Drosophila melanogaster - Ank;https://www.ncbi.nlm.nih.gov/nuccore/NM_001169348.2;https://www.ncbi.nlm.nih.gov/gene/?term=43770;FBgn
DMOV_RP1_hgx6470v1_pB66_A-186;5p 3p ;162046772;Drosophila melanogaster - Ank;https://www.ncbi.nlm.nih.gov/nuccore/NM_001169348.2;https://www.ncbi.nlm.nih.gov/gene/?term=43770;FBg
DMOV_RP1_hgx6470v1_pB66_A-18;5p 3p ;162046772;Drosophila melanogaster - Ank;https://www.ncbi.nlm.nih.gov/nuccore/NM_001169348.2;https://www.ncbi.nlm.nih.gov/gene/?term=43770;FBgn
DMOV_RP1_hgx6470v1_pB66_A-117;5p 3p ;162046772;Drosophila melanogaster - Ank;https://www.ncbi.nlm.nih.gov/nuccore/NM_001169348.2;https://www.ncbi.nlm.nih.gov/gene/?term=43770;FBg
DMOV_RP1_hgx6470v1_pB66_A-76;5p 3p ;162046772;Drosophila melanogaster - Ank;https://www.ncbi.nlm.nih.gov/nuccore/NM_001169348.2;https://www.ncbi.nlm.nih.gov/gene/?term=43770;FBgn
DMOV_RP1_hgx6470v1_pB66_A-81;5p 3p ;162046772;Drosophila melanogaster - Ank;https://www.ncbi.nlm.nih.gov/nuccore/NM_001169348.2;https://www.ncbi.nlm.nih.gov/gene/?term=43770;FBgn
DMOV_RP1_hgx6470v1_pB66_A-34;5p 3p ;162046772;Drosophila melanogaster - Ank;https://www.ncbi.nlm.nih.gov/nuccore/NM_001169348.2;https://www.ncbi.nlm.nih.gov/gene/?term=43770;FBgn
DMOV_RP1_hgx6470v1_pB66_A-43;5p 3p ;162046772;Drosophila melanogaster - Ank;https://www.ncbi.nlm.nih.gov/nuccore/NM_001169348.2;https://www.ncbi.nlm.nih.gov/gene/?term=43770;FBgn
DMOV_RP1_hgx6470v1_pB66_A-26;5p 3p ;162046772;Drosophila melanogaster - Ank;https://www.ncbi.nlm.nih.gov/nuccore/NM_001169348.2;https://www.ncbi.nlm.nih.gov/gene/?term=43770;FBgn
DMOV_RP1_hgx6470v1_pB66_A-32;5p 3p ;162046772;Drosophila melanogaster - Ank;https://www.ncbi.nlm.nih.gov/nuccore/NM_001169348.2;https://www.ncbi.nlm.nih.gov/gene/?term=43770;FBgn
DMOV_RP1_hgx6470v1_pB66_A-65;5p 3p ;162046772;Drosophila melanogaster - Ank;https://www.ncbi.nlm.nih.gov/nuccore/NM_001169348.2;https://www.ncbi.nlm.nih.gov/gene/?term=43770;FBgn
DMOV_RP1_hgx6470v1_pB66_A-72;5p 3p ;162046772;Drosophila melanogaster - Ank;https://www.ncbi.nlm.nih.gov/nuccore/NM_001169348.2;https://www.ncbi.nlm.nih.gov/gene/?term=43770;FBgn
DMOV_RP1_hgx6470v1_pB66_A-62;5p 3p ;162046772;Drosophila melanogaster - Ank;https://www.ncbi.nlm.nih.gov/nuccore/NM_001169348.2;https://www.ncbi.nlm.nih.gov/gene/?term=43770;FBgn
DMOV_RP1_hgx6470v1_pB66_A-110;5p 3p ;162046772;Drosophila melanogaster - Ank;https://www.ncbi.nlm.nih.gov/nuccore/NM_001169348.2;https://www.ncbi.nlm.nih.gov/gene/?term=43770;FBg
DMOV_RP1_hgx6470v1_pB66_A-96;5p 3p ;162046772;Drosophila melanogaster - Ank;https://www.ncbi.nlm.nih.gov/nuccore/NM_001169348.2;https://www.ncbi.nlm.nih.gov/gene/?term=43770;FBgn
DMOV_RP1_hgx6470v1_pB66_A-168;5p 3p ;162046772;Drosophila melanogaster - Ank;https://www.ncbi.nlm.nih.gov/nuccore/NM_001169348.2;https://www.ncbi.nlm.nih.gov/gene/?term=43770;FBg
DMOV_RP1_hgx6470v1_pB66_A-179;5p 3p ;162046772;Drosophila melanogaster - Ank;https://www.ncbi.nlm.nih.gov/nuccore/NM_001169348.2;https://www.ncbi.nlm.nih.gov/gene/?term=43770;FBg
DMOV_RP1_hgx6470v1_pB66_A-130;5p 3p ;162046772;Drosophila melanogaster - Ank;https://www.ncbi.nlm.nih.gov/nuccore/NM_001169348.2;https://www.ncbi.nlm.nih.gov/gene/?term=43770;FBg
DMOV_RP1_hgx6470v1_pB66_A-115;5p 3p ;162046772;Drosophila melanogaster - Ank;https://www.ncbi.nlm.nih.gov/nuccore/NM_001169348.2;https://www.ncbi.nlm.nih.gov/gene/?term=43770;FBg
DMOV_RP1_hgx6470v1_pB66_A-154;5p 3p ;162046772;Drosophila melanogaster - Ank;https://www.ncbi.nlm.nih.gov/nuccore/NM_001169348.2;https://www.ncbi.nlm.nih.gov/gene/?term=43770;FBg
DMOV_RP1_hgx6470v1_pB66_A-165;5p 3p ;162046772;Drosophila melanogaster - Ank;https://www.ncbi.nlm.nih.gov/nuccore/NM_001169348.2;https://www.ncbi.nlm.nih.gov/gene/?term=43770;FBg
DMOV_RP1_hgx6470v1_pB66_A-93;5p 3p ;162046772;Drosophila melanogaster - Ank;https://www.ncbi.nlm.nih.gov/nuccore/NM_001169348.2;https://www.ncbi.nlm.nih.gov/gene/?term=43770;FBgn
DMOV_RP1_hgx6470v1_pB66_A-77;5p 3p ;162046772;Drosophila melanogaster - Ank;https://www.ncbi.nlm.nih.gov/nuccore/NM_001169348.2;https://www.ncbi.nlm.nih.gov/gene/?term=43770;FBgn
DMOV_RP1_hgx6470v1_pB66_A-35;5p 3p ;162046772;Drosophila melanogaster - Ank;https://www.ncbi.nlm.nih.gov/nuccore/NM_001169348.2;https://www.ncbi.nlm.nih.gov/gene/?term=43770;FBgn

DMOV_RP1_hgx6470v1_pB66_A-49;5p ;162046772;Drosophila melanogaster - Ank;https://www.ncbi.nlm.nih.gov/nuccore/NM_001169348.2;https://www.ncbi.nlm.nih.gov/gene/?term=43770;FBgn00
DMOV_RP1_hgx6470v1_pB66_A-51;5p 3p ;162046772;Drosophila melanogaster - Ank;https://www.ncbi.nlm.nih.gov/nuccore/NM_001169348.2;https://www.ncbi.nlm.nih.gov/gene/?term=43770;FBg
DMOV_RP1_hgx6470v1_pB66_A-68;5p 3p ;162046772;Drosophila melanogaster - Ank;https://www.ncbi.nlm.nih.gov/nuccore/NM_001169348.2;https://www.ncbi.nlm.nih.gov/gene/?term=43770;FBg
DMOV_RP1_hgx6470v1_pB66_A-54;5p 3p ;162046772;Drosophila melanogaster - Ank;https://www.ncbi.nlm.nih.gov/nuccore/NM_001169348.2;https://www.ncbi.nlm.nih.gov/gene/?term=43770;FBg
DMOV_RP1_hgx6470v1_pB66_A-121;5p 3p ;162046772;Drosophila melanogaster - Ank;https://www.ncbi.nlm.nih.gov/nuccore/NM_001169348.2;https://www.ncbi.nlm.nih.gov/gene/?term=43770;FB
DMOV_RP1_hgx6470v1_pB66_A-167;5p 3p ;162046772;Drosophila melanogaster - Ank;https://www.ncbi.nlm.nih.gov/nuccore/NM_001169348.2;https://www.ncbi.nlm.nih.gov/gene/?term=43770;FB
DMOV_RP1_hgx6470v1_pB66_A-61;5p 3p ;162046772;Drosophila melanogaster - Ank;https://www.ncbi.nlm.nih.gov/nuccore/NM_001169348.2;https://www.ncbi.nlm.nih.gov/gene/?term=43770;FBg
DMOV_RP1_hgx6470v1_pB66_A-87;5p 3p ;162046772;Drosophila melanogaster - Ank;https://www.ncbi.nlm.nih.gov/nuccore/NM_001169348.2;https://www.ncbi.nlm.nih.gov/gene/?term=43770;FBg
DMOV_RP1_hgx6470v1_pB66_A-82;5p 3p ;162046772;Drosophila melanogaster - Ank;https://www.ncbi.nlm.nih.gov/nuccore/NM_001169348.2;https://www.ncbi.nlm.nih.gov/gene/?term=43770;FBg
DMOV_RP1_hgx6470v1_pB66_A-64;5p 3p ;162046772;Drosophila melanogaster - Ank;https://www.ncbi.nlm.nih.gov/nuccore/NM_001169348.2;https://www.ncbi.nlm.nih.gov/gene/?term=43770;FBg
DMOV_RP1_hgx6470v1_pB66_A-57;5p 3p ;162046772;Drosophila melanogaster - Ank;https://www.ncbi.nlm.nih.gov/nuccore/NM_001169348.2;https://www.ncbi.nlm.nih.gov/gene/?term=43770;FBg
DMOV_RP1_hgx6470v1_pB66_A-103;5p 3p ;162046772;Drosophila melanogaster - Ank;https://www.ncbi.nlm.nih.gov/nuccore/NM_001169348.2;https://www.ncbi.nlm.nih.gov/gene/?term=43770;FB
DMOV_RP1_hgx6470v1_pB66_A-183;5p 3p ;162046772;Drosophila melanogaster - Ank;https://www.ncbi.nlm.nih.gov/nuccore/NM_001169348.2;https://www.ncbi.nlm.nih.gov/gene/?term=43770;FB
DMOV_RP1_hgx6470v1_pB66_A-185;5p 3p ;162046772;Drosophila melanogaster - Ank;https://www.ncbi.nlm.nih.gov/nuccore/NM_001169348.2;https://www.ncbi.nlm.nih.gov/gene/?term=43770;FB
DMOV_RP1_hgx6470v1_pB66_A-174;5p 3p ;162046772;Drosophila melanogaster - Ank;https://www.ncbi.nlm.nih.gov/nuccore/NM_001169348.2;https://www.ncbi.nlm.nih.gov/gene/?term=43770;FB
DMOV_RP1_hgx6470v1_pB66_A-134;5p 3p ;162046772;Drosophila melanogaster - Ank;https://www.ncbi.nlm.nih.gov/nuccore/NM_001169348.2;https://www.ncbi.nlm.nih.gov/gene/?term=43770;FB
DMOV_RP1_hgx6470v1_pB66_A-83;5p 3p ;162046772;Drosophila melanogaster - Ank;https://www.ncbi.nlm.nih.gov/nuccore/NM_001169348.2;https://www.ncbi.nlm.nih.gov/gene/?term=43770;FBg
DMOV_RP1_hgx6470v1_pB66_A-66;5p 3p ;162046772;Drosophila melanogaster - Ank;https://www.ncbi.nlm.nih.gov/nuccore/NM_001169348.2;https://www.ncbi.nlm.nih.gov/gene/?term=43770;FBg
DMOV_RP1_hgx6470v1_pB66_A-9;5p 3p ;162046772;Drosophila melanogaster - Ank;https://www.ncbi.nlm.nih.gov/nuccore/NM_001169348.2;https://www.ncbi.nlm.nih.gov/gene/?term=43770;FBgn
DMOV_RP1_hgx6470v1_pB66_A-73;5p ;162046772;Drosophila melanogaster - Ank;https://www.ncbi.nlm.nih.gov/nuccore/NM_001169348.2;https://www.ncbi.nlm.nih.gov/gene/?term=43770;FBgn00
DMOV_RP1_hgx6470v1_pB66_A-104;5p 3p ;162046772;Drosophila melanogaster - Ank;https://www.ncbi.nlm.nih.gov/nuccore/NM_001169348.2;https://www.ncbi.nlm.nih.gov/gene/?term=43770;FB
DMOV_RP1_hgx6470v1_pB66_A-106;5p 3p ;162046772;Drosophila melanogaster - Ank;https://www.ncbi.nlm.nih.gov/nuccore/NM_001169348.2;https://www.ncbi.nlm.nih.gov/gene/?term=43770;FB
DMOV_RP1_hgx6470v1_pB66_A-5;5p 3p ;162046772;Drosophila melanogaster - Ank;https://www.ncbi.nlm.nih.gov/nuccore/NM_001169348.2;https://www.ncbi.nlm.nih.gov/gene/?term=43770;FBgn
DMOV_RP1_hgx6470v1_pB66_A-6;5p 3p ;162046772;Drosophila melanogaster - Ank;https://www.ncbi.nlm.nih.gov/nuccore/NM_001169348.2;https://www.ncbi.nlm.nih.gov/gene/?term=43770;FBgn
DMOV_RP1_hgx6470v1_pB66_A-1;5p ;162046772;Drosophila melanogaster - Ank;https://www.ncbi.nlm.nih.gov/nuccore/NM_001169348.2;https://www.ncbi.nlm.nih.gov/gene/?term=43770;FBgn001
DMOV_RP1_hgx6470v1_pB66_A-36;5p 3p ;162046772;Drosophila melanogaster - Ank;https://www.ncbi.nlm.nih.gov/nuccore/NM_001169348.2;https://www.ncbi.nlm.nih.gov/gene/?term=43770;FBg
DMOV_RP1_hgx6470v1_pB66_A-30;5p 3p ;162046772;Drosophila melanogaster - Ank;https://www.ncbi.nlm.nih.gov/nuccore/NM_001169348.2;https://www.ncbi.nlm.nih.gov/gene/?term=43770;FBg
DMOV_RP1_hgx6470v1_pB66_A-147;5p 3p ;162046772;Drosophila melanogaster - Ank;https://www.ncbi.nlm.nih.gov/nuccore/NM_001169348.2;https://www.ncbi.nlm.nih.gov/gene/?term=43770;FB
DMOV_RP1_hgx6470v1_pB66_A-182;5p 3p ;162046772;Drosophila melanogaster - Ank;https://www.ncbi.nlm.nih.gov/nuccore/NM_001169348.2;https://www.ncbi.nlm.nih.gov/gene/?term=43770;FB
DMOV_RP1_hgx6470v1_pB66_A-102;5p 3p ;162046772;Drosophila melanogaster - Ank;https://www.ncbi.nlm.nih.gov/nuccore/NM_001169348.2;https://www.ncbi.nlm.nih.gov/gene/?term=43770;FB
DMOV_RP1_hgx6470v1_pB66_A-172;5p 3p ;162046772;Drosophila melanogaster - Ank;https://www.ncbi.nlm.nih.gov/nuccore/NM_001169348.2;https://www.ncbi.nlm.nih.gov/gene/?term=43770;FB
DMOV_RP1_hgx6470v1_pB66_A-17;5p 3p ;162046772;Drosophila melanogaster - Ank;https://www.ncbi.nlm.nih.gov/nuccore/NM_001169348.2;https://www.ncbi.nlm.nih.gov/gene/?term=43770;FBg
DMOV_RP1_hgx6470v1_pB66_A-159;5p 3p ;162046772;Drosophila melanogaster - Ank;https://www.ncbi.nlm.nih.gov/nuccore/NM_001169348.2;https://www.ncbi.nlm.nih.gov/gene/?term=43770;FB
DMOV_RP1_hgx6470v1_pB66_A-14;5p 3p ;162046772;Drosophila melanogaster - Ank;https://www.ncbi.nlm.nih.gov/nuccore/NM_001169348.2;https://www.ncbi.nlm.nih.gov/gene/?term=43770;FBg
DMOV_RP1_hgx6470v1_pB66_A-114;5p 3p ;162046772;Drosophila melanogaster - Ank;https://www.ncbi.nlm.nih.gov/nuccore/NM_001169348.2;https://www.ncbi.nlm.nih.gov/gene/?term=43770;FB
DMOV_RP1_hgx6470v1_pB66_A-166;5p 3p ;162046772;Drosophila melanogaster - Ank;https://www.ncbi.nlm.nih.gov/nuccore/NM_001169348.2;https://www.ncbi.nlm.nih.gov/gene/?term=43770;FB
DMOV_RP1_hgx6470v1_pB66_A-126;5p 3p ;162046772;Drosophila melanogaster - Ank;https://www.ncbi.nlm.nih.gov/nuccore/NM_001169348.2;https://www.ncbi.nlm.nih.gov/gene/?term=43770;FB
DMOV_RP1_hgx6470v1_pB66_A-131;5p ;162046772;Drosophila melanogaster - Ank;https://www.ncbi.nlm.nih.gov/nuccore/NM_001169348.2;https://www.ncbi.nlm.nih.gov/gene/?term=43770;FBgn0
DMOV_RP1_hgx6470v1_pB66_A-156;5p 3p ;162046772;Drosophila melanogaster - Ank;https://www.ncbi.nlm.nih.gov/nuccore/NM_001169348.2;https://www.ncbi.nlm.nih.gov/gene/?term=43770;FB
DMOV_RP1_hgx6470v1_pB66_A-157;5p 3p ;162046772;Drosophila melanogaster - Ank;https://www.ncbi.nlm.nih.gov/nuccore/NM_001169348.2;https://www.ncbi.nlm.nih.gov/gene/?term=43770;FB
DMOV_RP1_hgx6470v1_pB66_A-161;5p 3p ;162046772;Drosophila melanogaster - Ank;https://www.ncbi.nlm.nih.gov/nuccore/NM_001169348.2;https://www.ncbi.nlm.nih.gov/gene/?term=43770;FB
DMOV_RP1_hgx6470v1_pB66_A-144;5p ;162046772;Drosophila melanogaster - Ank;https://www.ncbi.nlm.nih.gov/nuccore/NM_001169348.2;https://www.ncbi.nlm.nih.gov/gene/?term=43770;FBgn0
DMOV_RP1_hgx6470v1_pB66_A-152;5p 3p ;162046772;Drosophila melanogaster - Ank;https://www.ncbi.nlm.nih.gov/nuccore/NM_001169348.2;https://www.ncbi.nlm.nih.gov/gene/?term=43770;FB

DMOV_RP1_hgx6470v1_pB66_A-75;5p 3p ;162046772;Drosophila melanogaster - Ank;https://www.ncbi.nlm.nih.gov/nuccore/NM_001169348.2;https://www.ncbi.nlm.nih.gov/gene/?term=43770;FBg
DMOV_RP1_hgx6470v1_pB66_A-132;5p 3p ;162046772;Drosophila melanogaster - Ank;https://www.ncbi.nlm.nih.gov/nuccore/NM_001169348.2;https://www.ncbi.nlm.nih.gov/gene/?term=43770;FB
DMOV_RP1_hgx6470v1_pB66_A-94;5p 3p ;162046755;Drosophila melanogaster - CG4853;https://www.ncbi.nlm.nih.gov/nuccore/NM_001299606.1;https://www.ncbi.nlm.nih.gov/gene/?term=36974
DMOV_RP1_hgx6470v1_pB66_A-160;3p ;162046755;Drosophila melanogaster - CG4853;https://www.ncbi.nlm.nih.gov/nuccore/NM_001299606.1;https://www.ncbi.nlm.nih.gov/gene/?term=36974;F
DMOV_RP1_hgx6470v1_pB66_A-42;5p 3p ;162046901;Drosophila melanogaster - CG8891;https://www.ncbi.nlm.nih.gov/nuccore/NM_135046.5;https://www.ncbi.nlm.nih.gov/gene/?term=33718;FB
DMOV_RP1_hgx6470v1_pB66_A-176;5p 3p ;162046918;Drosophila melanogaster - Chrac-14;https://www.ncbi.nlm.nih.gov/nuccore/NM_057298.4;https://www.ncbi.nlm.nih.gov/gene/?term=37723
DMOV_RP1_hgx6470v1_pB66_A-116;5p 3p ;162046906;Drosophila melanogaster - Ef1alpha48D;https://www.ncbi.nlm.nih.gov/nuccore/NM_058027.5;https://www.ncbi.nlm.nih.gov/gene/?term=362
DMOV_RP1_hgx6470v1_pB66_A-8;5p 3p ;162046906;Drosophila melanogaster - Ef1alpha48D;https://www.ncbi.nlm.nih.gov/nuccore/NM_058027.5;https://www.ncbi.nlm.nih.gov/gene/?term=36271;
DMOV_RP1_hgx6470v1_pB66_A-46;5p 3p ;162046906;Drosophila melanogaster - Ef1alpha48D;https://www.ncbi.nlm.nih.gov/nuccore/NM_058027.5;https://www.ncbi.nlm.nih.gov/gene/?term=3627
DMOV_RP1_hgx6470v1_pB66_A-153;5p 3p ;162046760 / 162046769;Drosophila melanogaster - Moe;https://www.ncbi.nlm.nih.gov/nuccore/NM_080343.4;https://www.ncbi.nlm.nih.gov/gene/?term=
DMOV_RP1_hgx6470v1_pB66_A-175;5p 3p ;162046760 / 162046769;Drosophila melanogaster - Moe;https://www.ncbi.nlm.nih.gov/nuccore/NM_080343.4;https://www.ncbi.nlm.nih.gov/gene/?term=
DMOV_RP1_hgx6470v1_pB66_A-111;5p 3p ;162046760 / 162046769;Drosophila melanogaster - Moe;https://www.ncbi.nlm.nih.gov/nuccore/NM_080343.4;https://www.ncbi.nlm.nih.gov/gene/?term=
DMOV_RP1_hgx6470v1_pB66_A-16;5p 3p ;162046760 / 162046769;Drosophila melanogaster - Moe;https://www.ncbi.nlm.nih.gov/nuccore/NM_080343.4;https://www.ncbi.nlm.nih.gov/gene/?term=3
DMOV_RP1_hgx6470v1_pB66_A-47;5p 3p ;162046760 / 162046769;Drosophila melanogaster - Moe;https://www.ncbi.nlm.nih.gov/nuccore/NM_080343.4;https://www.ncbi.nlm.nih.gov/gene/?term=3
DMOV_RP1_hgx6470v1_pB66_A-59;5p 3p ;162046760 / 162046769;Drosophila melanogaster - Moe;https://www.ncbi.nlm.nih.gov/nuccore/NM_080343.4;https://www.ncbi.nlm.nih.gov/gene/?term=3
DMOV_RP1_hgx6470v1_pB66_A-135;5p 3p ;162046760 / 162046769;Drosophila melanogaster - Moe;https://www.ncbi.nlm.nih.gov/nuccore/NM_080343.4;https://www.ncbi.nlm.nih.gov/gene/?term=
DMOV_RP1_hgx6470v1_pB66_A-145;3p ;162046769;Drosophila melanogaster - Moe;https://www.ncbi.nlm.nih.gov/nuccore/NM_080343.4;https://www.ncbi.nlm.nih.gov/gene/?term=31816;FBgn001
DMOV_RP1_hgx6470v1_pB66_A-40;3p ;162046769;Drosophila melanogaster - Moe;https://www.ncbi.nlm.nih.gov/nuccore/NM_080343.4;https://www.ncbi.nlm.nih.gov/gene/?term=31816;FBgn0011
DMOV_RP1_hgx6470v1_pB66_A-173;5p 3p ;162046760 / 162046769;Drosophila melanogaster - Moe;https://www.ncbi.nlm.nih.gov/nuccore/NM_080343.4;https://www.ncbi.nlm.nih.gov/gene/?term=
DMOV_RP1_hgx6470v1_pB66_A-108;3p ;162046746;Drosophila melanogaster - Taf1;https://www.ncbi.nlm.nih.gov/nuccore/NM_001300272.1;https://www.ncbi.nlm.nih.gov/gene/?term=40813;FBgn
DMOV_RP1_hgx6470v1_pB66_A-3;5p 3p ;162046910 / 162046746;Drosophila melanogaster - Taf1;https://www.ncbi.nlm.nih.gov/nuccore/NM_001300272.1;https://www.ncbi.nlm.nih.gov/gene/?term
DMOV_RP1_hgx6470v1_pB66_A-86;5p ;162046910;Drosophila melanogaster - Taf1;https://www.ncbi.nlm.nih.gov/nuccore/NM_001300272.1;https://www.ncbi.nlm.nih.gov/gene/?term=40813;FBgn0
DMOV_RP1_hgx6470v1_pB66_A-141;5p 3p ;162046754 / 162046746;Drosophila melanogaster - Taf1;https://www.ncbi.nlm.nih.gov/nuccore/NM_001300272.1;https://www.ncbi.nlm.nih.gov/gene/?te
DMOV_RP1_hgx6470v1_pB66_A-78;5p 3p ;162046754 / 162046746;Drosophila melanogaster - Taf1;https://www.ncbi.nlm.nih.gov/nuccore/NM_001300272.1;https://www.ncbi.nlm.nih.gov/gene/?ter
DMOV_RP1_hgx6470v1_pB66_A-28;5p 3p ;162046746;Drosophila melanogaster - Taf1;https://www.ncbi.nlm.nih.gov/nuccore/NM_001300272.1;https://www.ncbi.nlm.nih.gov/gene/?term=40813;FBg
DMOV_RP1_hgx6470v1_pB66_A-187;5p 3p ;162046746;Drosophila melanogaster - Taf1;https://www.ncbi.nlm.nih.gov/nuccore/NM_001300272.1;https://www.ncbi.nlm.nih.gov/gene/?term=40813;FB
DMOV_RP1_hgx6470v1_pB66_A-11;5p 3p ;162046746;Drosophila melanogaster - Taf1;https://www.ncbi.nlm.nih.gov/nuccore/NM_001300272.1;https://www.ncbi.nlm.nih.gov/gene/?term=40813;FBg
DMOV_RP1_hgx6470v1_pB66_A-142;5p 3p ;162046920;Drosophila melanogaster - mRpL44;https://www.ncbi.nlm.nih.gov/nuccore/NM_141284.4;https://www.ncbi.nlm.nih.gov/gene/?term=40656;F
DMOV_RP1_hgx6470v1_pB66_A-190;5p 3p ;162046914 / 162046912;Drosophila melanogaster - polybromo;https://www.ncbi.nlm.nih.gov/nuccore/NM_143031.2;https://www.ncbi.nlm.nih.gov/gene/?
DMOV_RP1_hgx6470v1_pB66_A-74;5p 3p ;162046758;Drosophila melanogaster - stck;https://www.ncbi.nlm.nih.gov/nuccore/NM_001260058.2;https://www.ncbi.nlm.nih.gov/gene/?term=40999;FBg
DMOV_RP1_hgx6470v1_pB66_A-171;5p ;162046903;Drosophila melanogaster - GenMatch;https://www.ncbi.nlm.nih.gov/nuccore/CP122090.1;;;D;unknown| [prey3678174 - Droso - GenMatch];-1;N
DMOV_RP1_hgx6470v1_pB66_A-29;5p 3p ;162046903 / 162046915;Drosophila melanogaster - GenMatch;https://www.ncbi.nlm.nih.gov/nuccore/CP122090.1;;;D;unknown| [prey3678174 - Droso - G
DMOV_RP1_hgx6470v1_pB66_A-13;3p ;162046916;Drosophila melanogaster - GenMatch;https://www.ncbi.nlm.nih.gov/nuccore/CP122092.1;;;N/A;unknown| [prey3678176 - Droso - GenMatch];No

**Table 5** - Fas2 (aa 778-885) vs Drosophila Ovaries

Clone Name;Type Seq;Contig(s) Name;Gene Name (Best Match);GenBank ID (NCBI);Gene ID (NCBI);Other ID when available;Global PBS;Additional Gene Notes;Start;Stop;Frame;Sense;UTR Inclusion;% Id 5p/3p;Raw experimental 5p SEQ;Post linker extraction 5p SEQ;Raw experimental 3p SEQ;Post linker extraction 3p SEQ;Frag theoretical Sequence

DMOV_RP1_hgx5775v2_pB66_A-75;5p ;161682852;Drosophila melanogaster - CG10376;https://www.ncbi.nlm.nih.gov/nuccore/NM_136055.3;https://www.ncbi.nlm.nih.gov/gene/?term=35126;FBgn0032702;D;CG10376| [NM_136055.3 Drosophila melanogaster uncharacterized protein (CG10376)

DMOV_RP1_hgx5775v2_pB66_A-34;5p 3p ;161682877;Drosophila melanogaster - CG11873;https://www.ncbi.nlm.nih.gov/nuccore/NM_001276124.1;https://www.ncbi.nlm.nih.gov/gene/?term=43435;FBgn0039633;N/A;CG11873| [NM_001276124.1 Drosophila melanogaster uncharacterized protein

DMOV_RP1_hgx5775v2_pB66_A-33;5p 3p ;161682877;Drosophila melanogaster - CG11873;https://www.ncbi.nlm.nih.gov/nuccore/NM_001276124.1;https://www.ncbi.nlm.nih.gov/gene/?term=43435;FBgn0039633;N/A;CG11873| [NM_001276124.1 Drosophila melanogaster uncharacterized protein

DMOV_RP1_hgx5775v2_pB66_A-32;5p 3p ;161682880 / 161682951;Drosophila melanogaster - CG18135;https://www.ncbi.nlm.nih.gov/nuccore/NM_001275099.1;https://www.ncbi.nlm.nih.gov/gene/?term=40073;FBgn0036837;B;CG18135| [NM_001275099.1 Drosophila melanogaster uncharacterized protein

DMOV_RP1_hgx5775v2_pB66_A-26;5p 3p ;161682880;Drosophila melanogaster - CG18135;https://www.ncbi.nlm.nih.gov/nuccore/NM_001275099.1;https://www.ncbi.nlm.nih.gov/gene/?term=40073;FBgn0036837;B;CG18135| [NM_001275099.1 Drosophila melanogaster uncharacterized protein

DMOV_RP1_hgx5775v2_pB66_A-114;5p 3p ;161682880;Drosophila melanogaster - CG18135;https://www.ncbi.nlm.nih.gov/nuccore/NM_001275099.1;https://www.ncbi.nlm.nih.gov/gene/?term=40073;FBgn0036837;B;CG18135| [NM_001275099.1 Drosophila melanogaster uncharacterized protein

DMOV_RP1_hgx5775v2_pB66_A-117;5p 3p ;161682955;Drosophila melanogaster - CG33303;https://www.ncbi.nlm.nih.gov/nuccore/NM_001273406.1;https://www.ncbi.nlm.nih.gov/gene/?term=2768916;FBgn0053303;D;CG33303| [gi|442627247|ref|NM_001273406.1| Drosophila melanogaster CG33303 (CG33303)

DMOV_RP1_hgx5775v2_pB66_A-66;5p ;161682857;Drosophila melanogaster - CG3499;https://www.ncbi.nlm.nih.gov/nuccore/NM_001274213.2;https://www.ncbi.nlm.nih.gov/gene/?term=37636;FBgn0034792;C;CG3499| [prey1840196 - Droso CG3499];1269;No Data;IF;Sense;;99.6;

DMOV_RP1_hgx5775v2_pB66_A-105;5p 3p ;161682870;Drosophila melanogaster - CG4853;https://www.ncbi.nlm.nih.gov/nuccore/NM_001299606.1;https://www.ncbi.nlm.nih.gov/gene/?term=36974;FBgn0034230;N/A;CG4853| [NM_001299606.1 Drosophila melanogaster uncharacterized protein

DMOV_RP1_hgx5775v2_pB66_A-50;5p 3p ;161682870;Drosophila melanogaster - CG4853;https://www.ncbi.nlm.nih.gov/nuccore/NM_001299606.1;https://www.ncbi.nlm.nih.gov/gene/?term=36974;FBgn0034230;N/A;CG4853| [NM_001299606.1 Drosophila melanogaster uncharacterized protein

DMOV_RP1_hgx5775v2_pB66_A-52;5p 3p ;161682861 / 161682859;Drosophila melanogaster - CenG1A;https://www.ncbi.nlm.nih.gov/nuccore/NM_165067.2;https://www.ncbi.nlm.nih.gov/gene/?term=34803;FBgn0028509;D;CenG1A| [NM_165067.2 Drosophila melanogaster centaurin gamma 1A

DMOV_RP1_hgx5775v2_pB66_A-28;5p 3p ;161682884;Drosophila melanogaster - Dys;https://www.ncbi.nlm.nih.gov/nuccore/NM_001275804.1;https://www.ncbi.nlm.nih.gov/gene/?term=42327;FBgn0260003;D;Dys| [NM_001275804.1 Drosophila melanogaster dystrophin

DMOV_RP1_hgx5775v2_pB66_A-15;5p 3p ;161682954 / 161682952;Drosophila melanogaster - Plap;https://www.ncbi.nlm.nih.gov/nuccore/NM_001298628.1;https://www.ncbi.nlm.nih.gov/gene/?term=43978;FBgn0024314;D;Plap| [gi|665404077|ref|NM_001298628.1| Drosophila melanogaster phospholipase A2 activator protein (Plap)

DMOV_RP1_hgx5775v2_pB66_A-47;5p ;161682873;Drosophila melanogaster - SdhB;https://www.ncbi.nlm.nih.gov/nuccore/NM_057753.5;https://www.ncbi.nlm.nih.gov/gene/?term=35590;FBgn0014028;D;SdhB| [NM_057753.5 Drosophila melanogaster succinate dehydrogenase

DMOV_RP1_hgx5775v2_pB66_A-35;5p ;161682873;Drosophila melanogaster - SdhB;https://www.ncbi.nlm.nih.gov/nuccore/NM_057753.5;https://www.ncbi.nlm.nih.gov/gene/?term=35590;FBgn0014028;D;SdhB| [NM_057753.5 Drosophila melanogaster succinate dehydrogenase

DMOV_RP1_hgx5775v2_pB66_A-90;5p ;161682873;Drosophila melanogaster - SdhB;https://www.ncbi.nlm.nih.gov/nuccore/NM_057753.5;https://www.ncbi.nlm.nih.gov/gene/?term=35590;FBgn0014028;D;SdhB| [NM_057753.5 Drosophila melanogaster succinate dehydrogenase

DMOV_RP1_hgx5775v2_pB66_A-94;5p 3p ;161682862 / 161682970;Drosophila melanogaster - Taf1;https://www.ncbi.nlm.nih.gov/nuccore/NM_001300272.1;https://www.ncbi.nlm.nih.gov/gene/?term=40813;FBgn0010355;F;Taf1| [gi|665393367|ref|NM_001300272.1| Drosophila melanogaster TBP-associated factor 1 (Taf1)

DMOV_RP1_hgx5775v2_pB66_A-78;5p 3p ;161682862 / 161682970;Drosophila melanogaster - Taf1;https://www.ncbi.nlm.nih.gov/nuccore/NM_001300272.1;https://www.ncbi.nlm.nih.gov/gene/?term=40813;FBgn0010355;F;Taf1| [gi|665393367|ref|NM_001300272.1| Drosophila melanogaster TBP-associated factor 1 (Taf1)

DMOV_RP1_hgx5775v2_pB66_A-36;5p 3p ;161682862 / 161682977;Drosophila melanogaster - Taf1;https://www.ncbi.nlm.nih.gov/nuccore/NM_001300272.1;https://www.ncbi.nlm.nih.gov/gene/?term=40813;FBgn0010355;F;Taf1| [gi|665393367|ref|NM_001300272.1| Drosophila melanogaster TBP-associated factor 1 (Taf1)

DMOV_RP1_hgx5775v2_pB66_A-96;5p 3p ;161682862 / 161682977;Drosophila melanogaster - Taf1;https://www.ncbi.nlm.nih.gov/nuccore/NM_001300272.1;https://www.ncbi.nlm.nih.gov/gene/?term=40813;FBgn0010355;F;Taf1| [gi|665393367|ref|NM_001300272.1| Drosophila melanogaster TBP-associated factor 1 (Taf1)

DMOV_RP1_hgx5775v2_pB66_A-97;5p 3p ;161682862 / 161682977;Drosophila melanogaster - Taf1;https://www.ncbi.nlm.nih.gov/nuccore/NM_001300272.1;https://www.ncbi.nlm.nih.gov/gene/?term=40813;FBgn0010355;F;Taf1| [gi|665393367|ref|NM_001300272.1| Drosophila melanogaster TBP-associated factor 1 (Taf1)

DMOV_RP1_hgx5775v2_pB66_A-109;5p 3p ;161682970 / 161682977;Drosophila melanogaster - Taf1;https://www.ncbi.nlm.nih.gov/nuccore/NM_001300272.1;https://www.ncbi.nlm.nih.gov/gene/?term=40813;FBgn0010355;F;Taf1| [gi|665393367|ref|NM_001300272.1| Drosophila melanogaster TBP-associated factor 1 (Taf1)

DMOV_RP1_hgx5775v2_pB66_A-88;5p 3p ;161682970;Drosophila melanogaster - Taf1;https://www.ncbi.nlm.nih.gov/nuccore/NM_001300272.1;https://www.ncbi.nlm.nih.gov/gene/?term=40813;FBgn0010355;F;Taf1| [gi|665393367|ref|NM_001300272.1| Drosophila melanogaster TBP-associated factor 1 (Taf1)

DMOV_RP1_hgx5775v2_pB66_A-122;5p 3p ;161682970;Drosophila melanogaster - Taf1;https://www.ncbi.nlm.nih.gov/nuccore/NM_001300272.1;https://www.ncbi.nlm.nih.gov/gene/?term=40813;FBgn0010355;F;Taf1| [gi|665393367|ref|NM_001300272.1| Drosophila melanogaster TBP-associated factor 1 (Taf1)

DMOV_RP1_hgx5775v2_pB66_A-102;5p 3p ;161682970;Drosophila melanogaster - Taf1;https://www.ncbi.nlm.nih.gov/nuccore/NM_001300272.1;https://www.ncbi.nlm.nih.gov/gene/?term=40813;FBgn0010355;F;Taf1| [gi|665393367|ref|NM_001300272.1| Drosophila melanogaster TBP-associated factor 1 (Taf1)

DMOV_RP1_hgx5775v2_pB66_A-80;5p 3p ;161682970;Drosophila melanogaster - Taf1;https://www.ncbi.nlm.nih.gov/nuccore/NM_001300272.1;https://www.ncbi.nlm.nih.gov/gene/?term=40813;FBgn0010355;F;Taf1| [gi|665393367|ref|NM_001300272.1| Drosophila melanogaster TBP-associated factor 1 (Taf1)

DMOV_RP1_hgx5775v2_pB66_A-24;5p 3p ;161682970;Drosophila melanogaster - Taf1;https://www.ncbi.nlm.nih.gov/nuccore/NM_001300272.1;https://www.ncbi.nlm.nih.gov/gene/?term=40813;FBgn0010355;F;Taf1| [gi|665393367|ref|NM_001300272.1| Drosophila melanogaster TBP-associated factor 1 (Taf1)

DMOV_RP1_hgx5775v2_pB66_A-54;5p 3p ;161682868;Drosophila melanogaster - Tes;https://www.ncbi.nlm.nih.gov/nuccore/NM_137371.4;https://www.ncbi.nlm.nih.gov/gene/?term=36965;FBgn0034223;D;Tes| [NM_137371.4 Drosophila melanogaster testin (Tes)

DMOV_RP1_hgx5775v2_pB66_A-108;5p 3p ;161682980 / 161682978;Drosophila melanogaster - Tm1;https://www.ncbi.nlm.nih.gov/nuccore/NM_079636.4;https://www.ncbi.nlm.nih.gov/gene/?term=41852;FBgn0003721;N/A;Tm1| [NM_079636.4 Drosophila melanogaster tropomyosin 1

DMOV_RP1_hgx5775v2_pB66_A-99;5p 3p ;161682886;Drosophila melanogaster - dlg1;https://www.ncbi.nlm.nih.gov/nuccore/NM_001258693.1;https://www.ncbi.nlm.nih.gov/gene/?term=32083;FBgn0001624;A;dlg1| [gi|386764219|ref|NM_001258693.1| Drosophila melanogaster discs large 1 (dlg1)

DMOV_RP1_hgx5775v2_pB66_A-124;3p ;161682886;Drosophila melanogaster - dlg1;https://www.ncbi.nlm.nih.gov/nuccore/NM_001258693.1;https://www.ncbi.nlm.nih.gov/gene/?term=32083;FBgn0001624;A;dlg1| [gi|386764219|ref|NM_001258693.1| Drosophila melanogaster discs large 1 (dlg1)

DMOV_RP1_hgx5775v2_pB66_A-123;3p ;161682886;Drosophila melanogaster - dlg1;https://www.ncbi.nlm.nih.gov/nuccore/NM_001258693.1;https://www.ncbi.nlm.nih.gov/gene/?term=32083;FBgn0001624;A;dlg1| [gi|386764219|ref|NM_001258693.1| Drosophila melanogaster discs large 1 (dlg1)

DMOV_RP1_hgx5775v2_pB66_A-121;3p ;161682886;Drosophila melanogaster - dlg1;https://www.ncbi.nlm.nih.gov/nuccore/NM_001258693.1;https://www.ncbi.nlm.nih.gov/gene/?term=32083;FBgn0001624;A;dlg1| [gi|386764219|ref|NM_001258693.1| Drosophila melanogaster discs large 1 (dlg1)

DMOV_RP1_hgx5775v2_pB66_A-91;3p ;161682886;Drosophila melanogaster - dlg1;https://www.ncbi.nlm.nih.gov/nuccore/NM_001258693.1;https://www.ncbi.nlm.nih.gov/gene/?term=32083;FBgn0001624;A;dlg1| [gi|386764219|ref|NM_001258693.1| Drosophila melanogaster discs large 1 (dlg1)

DMOV_RP1_hgx5775v2_pB66_A-1;3p ;161682886;Drosophila melanogaster - dlg1;https://www.ncbi.nlm.nih.gov/nuccore/NM_001258693.1;https://www.ncbi.nlm.nih.gov/gene/?term=32083;FBgn0001624;A;dlg1| [gi|386764219|ref|NM_001258693.1| Drosophila melanogaster discs large 1 (dlg1)

DMOV_RP1_hgx5775v2_pB66_A-3;3p ;161682886;Drosophila melanogaster - dlg1;https://www.ncbi.nlm.nih.gov/nuccore/NM_001258693.1;https://www.ncbi.nlm.nih.gov/gene/?term=32083;FBgn0001624;A;dlg1| [gi|386764219|ref|NM_001258693.1| Drosophila melanogaster discs large 1 (dlg1)

DMOV_RP1_hgx5775v2_pB66_A-69;3p ;161682886;Drosophila melanogaster - dlg1;https://www.ncbi.nlm.nih.gov/nuccore/NM_001258693.1;https://www.ncbi.nlm.nih.gov/gene/?term=32083;FBgn0001624;A;dlg1| [gi|386764219|ref|NM_001258693.1| Drosophila melanogaster discs large 1 (dlg1)

DMOV_RP1_hgx5775v2_pB66_A-25;3p ;161682886;Drosophila melanogaster - dlg1;https://www.ncbi.nlm.nih.gov/nuccore/NM_001258693.1;https://www.ncbi.nlm.nih.gov/gene/?term=32083;FBgn0001624;A;dlg1| [gi|386764219|ref|NM_001258693.1| Drosophila melanogaster discs large 1 (dlg1)

DMOV_RP1_hgx5775v2_pB66_A-40;5p 3p ;161682886;Drosophila melanogaster - dlg1;https://www.ncbi.nlm.nih.gov/nuccore/NM_001258693.1;https://www.ncbi.nlm.nih.gov/gene/?term=32083;FBgn0001624;A;dlg1| [gi|386764219|ref|NM_001258693.1| Drosophila melanogaster discs large 1 (dlg1)

DMOV_RP1_hgx5775v2_pB66_A-67;5p 3p ;161682886;Drosophila melanogaster - dlg1;https://www.ncbi.nlm.nih.gov/nuccore/NM_001258693.1;https://www.ncbi.nlm.nih.gov/gene/?term=32083;FBgn0001624;A;dlg1| [gi|386764219|ref|NM_001258693.1| Drosophila melanogaster discs large 1 (dlg1)

DMOV_RP1_hgx5775v2_pB66_A-8;5p 3p ;161682886;Drosophila melanogaster - dlg1;https://www.ncbi.nlm.nih.gov/nuccore/NM_001258693.1;https://www.ncbi.nlm.nih.gov/gene/?term=32083;FBgn0001624;A;dlg1| [gi|386764219|ref|NM_001258693.1| Drosophila melanogaster discs large 1 (dlg1)

DMOV_RP1_hgx5775v2_pB66_A-5;5p 3p ;161682886;Drosophila melanogaster - dlg1;https://www.ncbi.nlm.nih.gov/nuccore/NM_001258693.1;https://www.ncbi.nlm.nih.gov/gene/?term=32083;FBgn0001624;A;dlg1| [gi|386764219|ref|NM_001258693.1| Drosophila melanogaster discs large 1 (dlg1)

DMOV_RP1_hgx5775v2_pB66_A-12;5p 3p ;161682886;Drosophila melanogaster - dlg1;https://www.ncbi.nlm.nih.gov/nuccore/NM_001258693.1;https://www.ncbi.nlm.nih.gov/gene/?term=32083;FBgn0001624;A;dlg1| [gi|386764219|ref|NM_001258693.1| Drosophila melanogaster discs large 1 (dlg1)

DMOV_RP1_hgx5775v2_pB66_A-29;5p 3p ;161682886;Drosophila melanogaster - dlg1;https://www.ncbi.nlm.nih.gov/nuccore/NM_001258693.1;https://www.ncbi.nlm.nih.gov/gene/?term=32083;FBgn0001624;A;dlg1| [gi|386764219|ref|NM_001258693.1| Drosophila melanogaster discs large 1 (dlg1)

DMOV_RP1_hgx5775v2_pB66_A-21;5p 3p ;161682886;Drosophila melanogaster - dlg1;https://www.ncbi.nlm.nih.gov/nuccore/NM_001258693.1;https://www.ncbi.nlm.nih.gov/gene/?term=32083;FBgn0001624;A;dlg1| [gi|386764219|ref|NM_001258693.1| Drosophila melanogaster discs large 1 (dlg1)

DMOV_RP1_hgx5775v2_pB66_A-48;5p 3p ;161682886;Drosophila melanogaster - dlg1;https://www.ncbi.nlm.nih.gov/nuccore/NM_001258693.1;https://www.ncbi.nlm.nih.gov/gene/?term=32083;FBgn0001624;A;dlg1| [gi|386764219|ref|NM_001258693.1| Drosophila melanogaster discs large 1 (dlg1)

DMOV_RP1_hgx5775v2_pB66_A-57;5p 3p ;161682886;Drosophila melanogaster - dlg1;https://www.ncbi.nlm.nih.gov/nuccore/NM_001258693.1;https://www.ncbi.nlm.nih.gov/gene/?term=32083;FBgn0001624;A;dlg1| [gi|386764219|ref|NM_001258693.1| Drosophila melanogaster discs large 1 (dlg1)

DMOV_RP1_hgx5775v2_pB66_A-51;5p 3p ;161682886;Drosophila melanogaster - dlg1;https://www.ncbi.nlm.nih.gov/nuccore/NM_001258693.1;https://www.ncbi.nlm.nih.gov/gene/?term=32083;FBgn0001624;A;dlg1| [gi|386764219|ref|NM_001258693.1| Drosophila melanogaster discs large 1 (dlg1)

DMOV_RP1_hgx5775v2_pB66_A-60;5p 3p ;161682886;Drosophila melanogaster - dlg1;https://www.ncbi.nlm.nih.gov/nuccore/NM_001258693.1;https://www.ncbi.nlm.nih.gov/gene/?term=32083;FBgn0001624;A;dlg1| [gi|386764219|ref|NM_001258693.1| Drosophila melanogaster discs large 1 (dlg1)

DMOV_RP1_hgx5775v2_pB66_A-68;5p 3p ;161682886;Drosophila melanogaster - dlg1;https://www.ncbi.nlm.nih.gov/nuccore/NM_001258693.1;https://www.ncbi.nlm.nih.gov/gene/?term=32083;FBgn0001624;A;dlg1| [gi|386764219|ref|NM_001258693.1| Drosophila melanogaster discs large 1 (dlg1)

DMOV_RP1_hgx5775v2_pB66_A-72;5p 3p ;161682886;Drosophila melanogaster - dlg1;https://www.ncbi.nlm.nih.gov/nuccore/NM_001258693.1;https://www.ncbi.nlm.nih.gov/gene/?term=32083;FBgn0001624;A;dlg1| [gi|386764219|ref|NM_001258693.1| Drosophila melanogaster discs large 1 (dlg1)

DMOV_RP1_hgx5775v2_pB66_A-45;5p 3p ;161682886;Drosophila melanogaster - dlg1;https://www.ncbi.nlm.nih.gov/nuccore/NM_001258693.1;https://www.ncbi.nlm.nih.gov/gene/?term=32083;FBgn0001624;A;dlg1| [gi|386764219|ref|NM_001258693.1| Drosophila melanogaster discs large 1 (dlg1)

DMOV_RP1_hgx5775v2_pB66_A-81;5p 3p ;161682886;Drosophila melanogaster - dlg1;https://www.ncbi.nlm.nih.gov/nuccore/NM_001258693.1;https://www.ncbi.nlm.nih.gov/gene/?term=32083;FBgn0001624;A;dlg1| [gi|386764219|ref|NM_001258693.1| Drosophila melanogaster discs large 1 (dlg1)

DMOV_RP1_hgx5775v2_pB66_A-6;5p 3p ;161682886;Drosophila melanogaster - dlg1;https://www.ncbi.nlm.nih.gov/nuccore/NM_001258693.1;https://www.ncbi.nlm.nih.gov/gene/?term=32083;FBgn0001624;A;dlg1| [gi|386764219|ref|NM_001258693.1| Drosophila melanogaster discs large 1 (dlg1)

DMOV_RP1_hgx5775v2_pB66_A-44;5p 3p ;161682886;Drosophila melanogaster - dlg1;https://www.ncbi.nlm.nih.gov/nuccore/NM_001258693.1;https://www.ncbi.nlm.nih.gov/gene/?term=32083;FBgn0001624;A;dlg1| [gi|386764219|ref|NM_001258693.1| Drosophila melanogaster discs large 1 (dlg1)

DMOV_RP1_hgx5775v2_pB66_A-71;5p 3p ;161682886;Drosophila melanogaster - dlg1;https://www.ncbi.nlm.nih.gov/nuccore/NM_001258693.1;https://www.ncbi.nlm.nih.gov/gene/?term=32083;FBgn0001624;A;dlg1| [gi|386764219|ref|NM_001258693.1| Drosophila melanogaster discs large 1 (dlg1)

DMOV_RP1_hgx5775v2_pB66_A-59;5p 3p ;161682886;Drosophila melanogaster - dlg1;https://www.ncbi.nlm.nih.gov/nuccore/NM_001258693.1;https://www.ncbi.nlm.nih.gov/gene/?term=32083;FBgn0001624;A;dlg1| [gi|386764219|ref|NM_001258693.1| Drosophila melanogaster discs large 1 (dlg1)

DMOV_RP1_hgx5775v2_pB66_A-82;5p 3p ;161682886 / 161682850;Drosophila melanogaster - dlg1;https://www.ncbi.nlm.nih.gov/nuccore/NM_001258693.1;https://www.ncbi.nlm.nih.gov/gene/?term=32083;FBgn0001624;A;dlg1| [gi|386764219|ref|NM_001258693.1| Drosophila melanogaster discs large 1 (dlg1)

DMOV_RP1_hgx5775v2_pB66_A-17;5p 3p ;161682886;Drosophila melanogaster - dlg1;https://www.ncbi.nlm.nih.gov/nuccore/NM_001258693.1;https://www.ncbi.nlm.nih.gov/gene/?term=32083;FBgn0001624;A;dlg1| [gi|386764219|ref|NM_001258693.1| Drosophila melanogaster discs large 1 (dlg1)

DMOV_RP1_hgx5775v2_pB66_A-74;5p 3p ;161682886;Drosophila melanogaster - dlg1;https://www.ncbi.nlm.nih.gov/nuccore/NM_001258693.1;https://www.ncbi.nlm.nih.gov/gene/?term=32083;FBgn0001624;A;dlg1| [gi|386764219|ref|NM_001258693.1| Drosophila melanogaster discs large 1 (dlg1)

DMOV_RP1_hgx5775v2_pB66_A-98;5p 3p ;161682886;Drosophila melanogaster - dlg1;https://www.ncbi.nlm.nih.gov/nuccore/NM_001258693.1;https://www.ncbi.nlm.nih.gov/gene/?term=32083;FBgn0001624;A;dlg1| [gi|386764219|ref|NM_001258693.1| Drosophila melanogaster discs large 1 (dlg1)

DMOV_RP1_hgx5775v2_pB66_A-100;5p 3p ;161682886;Drosophila melanogaster - dlg1;https://www.ncbi.nlm.nih.gov/nuccore/NM_001258693.1;https://www.ncbi.nlm.nih.gov/gene/?term=32083;FBgn0001624;A;dlg1| [gi|386764219|ref|NM_001258693.1| Drosophila melanogaster discs large 1 (dlg1)

DMOV_RP1_hgx5775v2_pB66_A-128;5p 3p ;161682886;Drosophila melanogaster - dlg1;https://www.ncbi.nlm.nih.gov/nuccore/NM_001258693.1;https://www.ncbi.nlm.nih.gov/gene/?term=32083;FBgn0001624;A;dlg1| [gi|386764219|ref|NM_001258693.1| Drosophila melanogaster discs large 1 (dlg1)

DMOV_RP1_hgx5775v2_pB66_A-9;5p 3p ;161682886;Drosophila melanogaster - dlg1;https://www.ncbi.nlm.nih.gov/nuccore/NM_001258693.1;https://www.ncbi.nlm.nih.gov/gene/?term=32083;FBgn0001624;A;dlg1| [gi|386764219|ref|NM_001258693.1| Drosophila melanogaster discs large 1 (dlg1)

DMOV_RP1_hgx5775v2_pB66_A-70;5p 3p ;161682886;Drosophila melanogaster - dlg1;https://www.ncbi.nlm.nih.gov/nuccore/NM_001258693.1;https://www.ncbi.nlm.nih.gov/gene/?term=32083;FBgn0001624;A;dlg1| [gi|386764219|ref|NM_001258693.1| Drosophila melanogaster discs large 1 (dlg1)

DMOV_RP1_hgx5775v2_pB66_A-116;5p 3p ;161682886;Drosophila melanogaster - dlg1;https://www.ncbi.nlm.nih.gov/nuccore/NM_001258693.1;https://www.ncbi.nlm.nih.gov/gene/?term=32083;FBgn0001624;A;dlg1| [gi|386764219|ref|NM_001258693.1| Drosophila melanogaster discs large 1 (dlg1)

DMOV_RP1_hgx5775v2_pB66_A-111;5p 3p ;161682886;Drosophila melanogaster - dlg1;https://www.ncbi.nlm.nih.gov/nuccore/NM_001258693.1;https://www.ncbi.nlm.nih.gov/gene/?term=32083;FBgn0001624;A;dlg1| [gi|386764219|ref|NM_001258693.1| Drosophila melanogaster discs large 1 (dlg1)

DMOV_RP1_hgx5775v2_pB66_A-110;5p ;161682886;Drosophila melanogaster - dlg1;https://www.ncbi.nlm.nih.gov/nuccore/NM_001258693.1;https://www.ncbi.nlm.nih.gov/gene/?term=32083;FBgn0001624;A;dlg1| [gi|386764219|ref|NM_001258693.1| Drosophila melanogaster discs large 1 (dlg1)

DMOV_RP1_hgx5775v2_pB66_A-119;5p 3p ;161682886;Drosophila melanogaster - dlg1;https://www.ncbi.nlm.nih.gov/nuccore/NM_001258693.1;https://www.ncbi.nlm.nih.gov/gene/?term=32083;FBgn0001624;A;dlg1| [gi|386764219|ref|NM_001258693.1| Drosophila melanogaster discs large 1 (dlg1)

DMOV_RP1_hgx5775v2_pB66_A-27;5p 3p ;161682886;Drosophila melanogaster - dlg1;https://www.ncbi.nlm.nih.gov/nuccore/NM_001258693.1;https://www.ncbi.nlm.nih.gov/gene/?term=32083;FBgn0001624;A;dlg1| [gi|386764219|ref|NM_001258693.1| Drosophila melanogaster discs large 1 (dlg1)

DMOV_RP1_hgx5775v2_pB66_A-65;5p 3p ;161682886;Drosophila melanogaster - dlg1;https://www.ncbi.nlm.nih.gov/nuccore/NM_001258693.1;https://www.ncbi.nlm.nih.gov/gene/?term=32083;FBgn0001624;A;dlg1| [gi|386764219|ref|NM_001258693.1| Drosophila melanogaster discs large 1 (dlg1)

DMOV_RP1_hgx5775v2_pB66_A-79;5p 3p ;161682886;Drosophila melanogaster - dlg1;https://www.ncbi.nlm.nih.gov/nuccore/NM_001258693.1;https://www.ncbi.nlm.nih.gov/gene/?term=32083;FBgn0001624;A;dlg1| [gi|386764219|ref|NM_001258693.1| Drosophila melanogaster discs large 1 (dlg1)

DMOV_RP1_hgx5775v2_pB66_A-106;5p 3p ;161682886;Drosophila melanogaster - dlg1;https://www.ncbi.nlm.nih.gov/nuccore/NM_001258693.1;https://www.ncbi.nlm.nih.gov/gene/?term=32083;FBgn0001624;A;dlg1| [gi|386764219|ref|NM_001258693.1| Drosophila melanogaster discs large 1 (dlg1)

DMOV_RP1_hgx5775v2_pB66_A-129;5p 3p ;161682886;Drosophila melanogaster - dlg1;https://www.ncbi.nlm.nih.gov/nuccore/NM_001258693.1;https://www.ncbi.nlm.nih.gov/gene/?term=32083;FBgn0001624;A;dlg1| [gi|386764219|ref|NM_001258693.1| Drosophila melanogaster discs large 1 (dlg1)

DMOV_RP1_hgx5775v2_pB66_A-7;5p 3p ;161682886;Drosophila melanogaster - dlg1;https://www.ncbi.nlm.nih.gov/nuccore/NM_001258693.1;https://www.ncbi.nlm.nih.gov/gene/?term=32083;FBgn0001624;A;dlg1| [gi|386764219|ref|NM_001258693.1| Drosophila melanogaster discs large 1 (dlg1)

DMOV_RP1_hgx5775v2_pB66_A-62;5p 3p ;161682886;Drosophila melanogaster - dlg1;https://www.ncbi.nlm.nih.gov/nuccore/NM_001258693.1;https://www.ncbi.nlm.nih.gov/gene/?term=32083;FBgn0001624;A;dlg1| [gi|386764219|ref|NM_001258693.1| Drosophila melanogaster discs large 1 (dlg1)

DMOV_RP1_hgx5775v2_pB66_A-61;5p 3p ;161682886;Drosophila melanogaster - dlg1;https://www.ncbi.nlm.nih.gov/nuccore/NM_001258693.1;https://www.ncbi.nlm.nih.gov/gene/?term=32083;FBgn0001624;A;dlg1| [gi|386764219|ref|NM_001258693.1| Drosophila melanogaster discs large 1 (dlg1)
DMOV_RP1_hgx5775v2_pB66_A-104;5p ;161682886;Drosophila melanogaster - dlg1;https://www.ncbi.nlm.nih.gov/nuccore/NM_001258693.1;https://www.ncbi.nlm.nih.gov/gene/?term=32083;FBgn0001624;A;dlg1| [gi|386764219|ref|NM_001258693.1| Drosophila melanogaster discs large 1 (dlg1)
DMOV_RP1_hgx5775v2_pB66_A-13;5p 3p ;161682886;Drosophila melanogaster - dlg1;https://www.ncbi.nlm.nih.gov/nuccore/NM_001258693.1;https://www.ncbi.nlm.nih.gov/gene/?term=32083;FBgn0001624;A;dlg1| [gi|386764219|ref|NM_001258693.1| Drosophila melanogaster discs large 1 (dlg1)
DMOV_RP1_hgx5775v2_pB66_A-132;5p 3p ;161682886;Drosophila melanogaster - dlg1;https://www.ncbi.nlm.nih.gov/nuccore/NM_001258693.1;https://www.ncbi.nlm.nih.gov/gene/?term=32083;FBgn0001624;A;dlg1| [gi|386764219|ref|NM_001258693.1| Drosophila melanogaster discs large 1 (dlg1)
DMOV_RP1_hgx5775v2_pB66_A-49;5p 3p ;161682886;Drosophila melanogaster - dlg1;https://www.ncbi.nlm.nih.gov/nuccore/NM_001258693.1;https://www.ncbi.nlm.nih.gov/gene/?term=32083;FBgn0001624;A;dlg1| [gi|386764219|ref|NM_001258693.1| Drosophila melanogaster discs large 1 (dlg1)
DMOV_RP1_hgx5775v2_pB66_A-63;5p 3p ;161682886;Drosophila melanogaster - dlg1;https://www.ncbi.nlm.nih.gov/nuccore/NM_001258693.1;https://www.ncbi.nlm.nih.gov/gene/?term=32083;FBgn0001624;A;dlg1| [gi|386764219|ref|NM_001258693.1| Drosophila melanogaster discs large 1 (dlg1)
DMOV_RP1_hgx5775v2_pB66_A-85;5p 3p ;161682886;Drosophila melanogaster - dlg1;https://www.ncbi.nlm.nih.gov/nuccore/NM_001258693.1;https://www.ncbi.nlm.nih.gov/gene/?term=32083;FBgn0001624;A;dlg1| [gi|386764219|ref|NM_001258693.1| Drosophila melanogaster discs large 1 (dlg1)
DMOV_RP1_hgx5775v2_pB66_A-107;5p 3p ;161682886;Drosophila melanogaster - dlg1;https://www.ncbi.nlm.nih.gov/nuccore/NM_001258693.1;https://www.ncbi.nlm.nih.gov/gene/?term=32083;FBgn0001624;A;dlg1| [gi|386764219|ref|NM_001258693.1| Drosophila melanogaster discs large 1 (dlg1)
DMOV_RP1_hgx5775v2_pB66_A-77;5p 3p ;161682886;Drosophila melanogaster - dlg1;https://www.ncbi.nlm.nih.gov/nuccore/NM_001258693.1;https://www.ncbi.nlm.nih.gov/gene/?term=32083;FBgn0001624;A;dlg1| [gi|386764219|ref|NM_001258693.1| Drosophila melanogaster discs large 1 (dlg1)
DMOV_RP1_hgx5775v2_pB66_A-30;5p 3p ;161682886;Drosophila melanogaster - dlg1;https://www.ncbi.nlm.nih.gov/nuccore/NM_001258693.1;https://www.ncbi.nlm.nih.gov/gene/?term=32083;FBgn0001624;A;dlg1| [gi|386764219|ref|NM_001258693.1| Drosophila melanogaster discs large 1 (dlg1)
DMOV_RP1_hgx5775v2_pB66_A-20;5p 3p ;161682948 / 161682886;Drosophila melanogaster - dlg1 varL;https://www.ncbi.nlm.nih.gov/nuccore/NM_001103486.3;https://www.ncbi.nlm.nih.gov/gene/?term=32083;FBgn0001624;B;dlg1 varL| [NM_001103486.3 Drosophila melanogaster discs large 1 (dlg1)
DMOV_RP1_hgx5775v2_pB66_A-38;5p 3p ;161682948 / 161682886;Drosophila melanogaster - dlg1 varL;https://www.ncbi.nlm.nih.gov/nuccore/NM_001103486.3;https://www.ncbi.nlm.nih.gov/gene/?term=32083;FBgn0001624;B;dlg1 varL| [NM_001103486.3 Drosophila melanogaster discs large 1 (dlg1)
DMOV_RP1_hgx5775v2_pB66_A-14;5p 3p ;161682948 / 161682886;Drosophila melanogaster - dlg1 varL;https://www.ncbi.nlm.nih.gov/nuccore/NM_001103486.3;https://www.ncbi.nlm.nih.gov/gene/?term=32083;FBgn0001624;B;dlg1 varL| [NM_001103486.3 Drosophila melanogaster discs large 1 (dlg1)
DMOV_RP1_hgx5775v2_pB66_A-4;5p 3p ;161682948 / 161682886;Drosophila melanogaster - dlg1 varL;https://www.ncbi.nlm.nih.gov/nuccore/NM_001103486.3;https://www.ncbi.nlm.nih.gov/gene/?term=32083;FBgn0001624;B;dlg1 varL| [NM_001103486.3 Drosophila melanogaster discs large 1 (dlg1)
DMOV_RP1_hgx5775v2_pB66_A-86;5p 3p ;161682948 / 161682886;Drosophila melanogaster - dlg1 varL;https://www.ncbi.nlm.nih.gov/nuccore/NM_001103486.3;https://www.ncbi.nlm.nih.gov/gene/?term=32083;FBgn0001624;B;dlg1 varL| [NM_001103486.3 Drosophila melanogaster discs large 1 (dlg1)
DMOV_RP1_hgx5775v2_pB66_A-101;3p ;161682960;Drosophila melanogaster - dom;https://www.ncbi.nlm.nih.gov/nuccore/NM_001299747.1;https://www.ncbi.nlm.nih.gov/gene/?term=45655;FBgn0020306;N/A;dom| [gi|665402523|ref|NM_001299747.1| Drosophila melanogaster domino (dom)
DMOV_RP1_hgx5775v2_pB66_A-92;3p ;161682960;Drosophila melanogaster - dom;https://www.ncbi.nlm.nih.gov/nuccore/NM_001299747.1;https://www.ncbi.nlm.nih.gov/gene/?term=45655;FBgn0020306;N/A;dom| [gi|665402523|ref|NM_001299747.1| Drosophila melanogaster domino (dom)
DMOV_RP1_hgx5775v2_pB66_A-73;5p 3p ;161682856 / 161682854;Drosophila melanogaster - gammaTub37C;https://www.ncbi.nlm.nih.gov/nuccore/NM_001273636.1;https://www.ncbi.nlm.nih.gov/gene/?term=35199;FBgn0010097;N/A;gammaTub37C| [NM_001273636.1 Drosophila melanogaster gamma-Tubulin at 37C (gammaTub37C)
DMOV_RP1_hgx5775v2_pB66_A-83;5p 3p ;161682963;Drosophila melanogaster - scrib;https://www.ncbi.nlm.nih.gov/nuccore/NM_001276060.1;https://www.ncbi.nlm.nih.gov/gene/?term=44448;FBgn0263289;B;scrib| [gi|442621261|ref|NM_001276060.1| Drosophila melanogaster scribbled (scrib)
DMOV_RP1_hgx5775v2_pB66_A-95;5p 3p ;161682963;Drosophila melanogaster - scrib;https://www.ncbi.nlm.nih.gov/nuccore/NM_001276060.1;https://www.ncbi.nlm.nih.gov/gene/?term=44448;FBgn0263289;B;scrib| [gi|442621261|ref|NM_001276060.1| Drosophila melanogaster scribbled (scrib)
DMOV_RP1_hgx5775v2_pB66_A-11;5p 3p ;161682963;Drosophila melanogaster - scrib;https://www.ncbi.nlm.nih.gov/nuccore/NM_001276060.1;https://www.ncbi.nlm.nih.gov/gene/?term=44448;FBgn0263289;B;scrib| [gi|442621261|ref|NM_001276060.1| Drosophila melanogaster scribbled (scrib)
DMOV_RP1_hgx5775v2_pB66_A-126;5p 3p ;161682963;Drosophila melanogaster - scrib;https://www.ncbi.nlm.nih.gov/nuccore/NM_001276060.1;https://www.ncbi.nlm.nih.gov/gene/?term=44448;FBgn0263289;B;scrib| [gi|442621261|ref|NM_001276060.1| Drosophila melanogaster scribbled (scrib)
DMOV_RP1_hgx5775v2_pB66_A-89;5p 3p ;161682963;Drosophila melanogaster - scrib;https://www.ncbi.nlm.nih.gov/nuccore/NM_001276060.1;https://www.ncbi.nlm.nih.gov/gene/?term=44448;FBgn0263289;B;scrib| [gi|442621261|ref|NM_001276060.1| Drosophila melanogaster scribbled (scrib)
DMOV_RP1_hgx5775v2_pB66_A-64;5p 3p ;161682963;Drosophila melanogaster - scrib;https://www.ncbi.nlm.nih.gov/nuccore/NM_001276060.1;https://www.ncbi.nlm.nih.gov/gene/?term=44448;FBgn0263289;B;scrib| [gi|442621261|ref|NM_001276060.1| Drosophila melanogaster scribbled (scrib)
DMOV_RP1_hgx5775v2_pB66_A-46;3p ;161682949;Drosophila melanogaster - GenMatch;https://www.ncbi.nlm.nih.gov/nuccore/AC023718.4;;;N/A;unknown| [prey3672821 - Droso - GenMatch];No Data;501;??;Sense;;100.0;;;
DMOV_RP1_hgx5775v2_pB66_A-93;5p 3p ;161682957;Drosophila melanogaster - GenMatch;https://www.ncbi.nlm.nih.gov/nuccore/AE014297.3;;;D;unknown| [prey3672822 - Droso - GenMatch];-1;790;IF;Sense;;99.5 /
DMOV_RP1_hgx5775v2_pB66_A-120;5p 3p ;161682957;Drosophila melanogaster - GenMatch;https://www.ncbi.nlm.nih.gov/nuccore/AE014297.3;;;D;unknown| [prey3672822 - Droso - GenMatch];-1;790;IF;Sense;;100.0 / 99.7;

**Table 6**
Finegan & Linhoff et al.

| Overlap | Miao *et al.* | | | | | Slaidina *et al.* | | | | | Fly Cell Atlas | | | | |
|---|---|---|---|---|---|---|---|---|---|---|---|---|---|---|---|
| **Gene** | logFC | pval | adj. pval | Stage 2-6 | Stage 8-9 | logFC | pval | adj. pval | Stage 2-6 | Stage 7-9 | logFC | pval | adj. pval | Stage 2-5 | Stage 6-8 |
| CG42788 | 2.163982461 | 1.14798E-22 | 5.6452E-21 | 1.566775244 | 0.349609375 | 1.094762046 | 2.3157E-24 | 1.34994E-23 | 0.12144394 | 0.056861673 | 0.677380241 | 6.36264E-08 | 6.80099E-07 | 2.394096839 | 1.497028064 |
| CG33970 | 3.177016789 | 1.32679E-17 | 4.07502E-16 | 0.918566775 | 0.1015625 | 2.792873179 | 9.2095E-107 | 2.1868E-105 | 0.20965814 | 0.030253326 | 1.283170303 | 3.45979E-23 | 8.93793E-22 | 2.316225495 | 0.951717569 |
| FER | 1.559268678 | 1.63425E-20 | 6.39419E-19 | 3.286644951 | 1.115234375 | 1.686195713 | 1.0028E-158 | 3.95E-157 | 0.508486732 | 0.158009841 | 1.789087053 | 6.10652E-69 | 5.38888E-67 | 3.687794985 | 1.067082264 |
| jigr1 | 2.713461975 | 7.67981E-79 | 7.67895E-76 | 8.006514658 | 1.220703125 | 2.068212329 | 0 | 0 | 1.132679895 | 0.270092947 | 3.156188018 | 3.1599E-268 | 2.0504E-265 | 6.880312299 | 0.77179309 |
| rumi | 1.335467692 | 6.3562E-12 | 1.10211E-10 | 0.749185668 | 0.296875 | 0.89790191 | 7.98074E-45 | 7.60192E-44 | 0.337556777 | 0.181155458 | 1.31021385 | 0.002493148 | 0.014125279 | 0.14713004 | 0.059331822 |
| Hs6st | 1.332648676 | 9.16846E-11 | 1.39135E-09 | 0.964169381 | 0.3828125 | 0.851168844 | 8.06307E-22 | 4.37651E-21 | 0.167344011 | 0.092764717 | 0.878184905 | 1.15693E-26 | 3.45858E-25 | 6.051935104 | 3.292564293 |
| mfas | 2.969230701 | 6.4358E-33 | 7.0629E-31 | 2.462540717 | 0.314453125 | 3.05823617 | 2.60461E-49 | 2.68024E-48 | 0.091082955 | 0.010934937 | 1.056360312 | 6.04281E-15 | 1.05975E-13 | 1.365762155 | 0.656718022 |
| CG7059 | 5.972209384 | 1.20562E-22 | 5.89642E-21 | 1.716612378 | 0.02734375 | 4.128445172 | 8.0119E-182 | 3.7364E-180 | 0.430313172 | 0.024603609 | 4.08345915 | 1.60645E-13 | 2.61754E-12 | 0.227925482 | 0.01344464 |
| wake | 1.751201977 | 2.73324E-20 | 1.04666E-18 | 1.42019544 | 0.421875 | 1.330920746 | 2.25205E-65 | 3.0963E-64 | 0.273248864 | 0.108620375 | 0.821026318 | 6.32838E-17 | 1.22043E-15 | 3.060126265 | 1.732151194 |
| CG7702 | 1.773974409 | 2.23458E-15 | 5.47928E-14 | 1.055374593 | 0.30859375 | 3.443164567 | 0 | 0 | 1.256753526 | 0.115545836 | 2.860978967 | 1.6759E-147 | 3.6248E-145 | 3.926601553 | 0.540475918 |
| SPARC | 4.610483526 | 2.61188E-59 | 1.17522E-56 | 14.07491857 | 0.576171875 | 4.063679532 | 0 | 0 | 20.65646665 | 1.235283397 | 2.626470741 | 6.5739E-228 | 3.2962E-225 | 9.302874883 | 1.50650544 |
| E(spl)m2-BFM | 3.664903324 | 2.94152E-32 | 3.04261E-30 | 1.882736156 | 0.1484375 | 2.640994739 | 0 | 0 | 1.630169735 | 0.261344997 | 3.353770459 | 5.45898E-08 | 5.88641E-07 | 0.14911718 | 0.0145862 |
| Rad60 | 1.165766694 | 4.98795E-10 | 6.88444E-09 | 0.736156352 | 0.328125 | 1.465395887 | 4.43663E-62 | 5.78598E-61 | 0.215395649 | 0.078002551 | #NUM! | 0.002635552 | 0.014878597 | 0.023126777 | 0 |
| CG31176 | 1.596265856 | 1.97105E-14 | 4.50191E-13 | 1.54723127 | 0.51171875 | 0.843800211 | 7.82091E-28 | 5.00736E-27 | 0.284245757 | 0.158374339 | 2.553111079 | 1.52612E-72 | 1.39129E-70 | 2.516335527 | 0.428751381 |
| Cenp-C | 1.808924618 | 5.53066E-13 | 1.07264E-11 | 0.745928339 | 0.212890625 | 2.1850287 | 2.7954E-172 | 1.2026E-170 | 0.385369352 | 0.084745763 | 1.910644826 | 0.005847337 | 0.030282615 | 0.032649154 | 0.008683814 |
| CG17230 | 1.529318533 | 0.000796364 | 0.003223787 | 0.146579805 | 0.05078125 | 1.077819099 | 6.35546E-08 | 2.01683E-07 | 0.033468802 | 0.015855659 | 0.615970019 | 0.002870529 | 0.016065351 | 0.39907368 | 0.260391998 |
| CycB3 | 1.313217485 | 2.68477E-08 | 2.8592E-07 | 0.48534202 | 0.1953125 | 1.565508978 | 4.99066E-63 | 6.5896E-62 | 0.210375329 | 0.071077091 | 2.029123884 | 0.002333284 | 0.013260412 | 0.051231386 | 0.012551885 |
| srp | 3.601903656 | 1.16541E-60 | 5.82642E-58 | 3.13029316 | 0.2578125 | 2.452935781 | 1.2462E-132 | 3.9374E-131 | 0.269423858 | 0.049207217 | 1.664730387 | 1.38837E-46 | 7.5444E-45 | 2.555280369 | 0.805943834 |
| stg | 1.848388103 | 5.30228E-37 | 8.22676E-35 | 12.68078176 | 3.521484375 | 1.098809058 | 0 | 0 | 7.087736075 | 3.309276472 | 2.386292057 | 4.977E-129 | 9.3053E-127 | 2.913869985 | 0.557345735 |
| glec | 1.563975193 | 5.84144E-25 | 3.48126E-23 | 4.296416938 | 1.453125 | 0.875884704 | 4.08948E-97 | 8.7165E-96 | 1.957207746 | 1.066520868 | 1.244207987 | 9.18369E-27 | 2.7679E-25 | 2.036985085 | 0.859892031 |
| Tsp96F | 2.692101465 | 1.27169E-10 | 1.89156E-09 | 0.302931596 | 0.046875 | 2.125235616 | 1.20848E-47 | 1.20941E-46 | 0.10733923 | 0.024603609 | #NUM! | 0.003207039 | 0.017777311 | 0.016046662 | 0 |
| hdc | 2.14717176 | 2.3281E-81 | 2.99294E-78 | 29.5276873 | 6.666015625 | 1.992344453 | 0 | 0 | 7.043987569 | 1.77036632 | 2.073847002 | 0 | 0 | 79.5712742 | 18.90018951 |
| CG7530 | 0.710204559 | 1.19736E-09 | 1.57991E-08 | 2.016286645 | 1.232421875 | 0.69103986 | 1.25982E-55 | 1.45141E-54 | 0.765001195 | 0.473847275 | 0.587353217 | 2.0929E-11 | 2.97893E-10 | 3.055512192 | 2.033635359 |
| E(spl)m7-HLH | 3.111094265 | 2.30557E-37 | 3.70497E-35 | 3.172638436 | 0.3671875 | 1.405187832 | 1.9465E-204 | 1.0964E-202 | 1.971790581 | 0.744486969 | 2.62196539 | 7.99229E-16 | 1.45483E-14 | 0.391793806 | 0.063645446 |
| CG5455 | 2.217374527 | 1.11732E-24 | 6.48693E-23 | 0.944625407 | 0.203125 | 0.800766637 | 4.27229E-28 | 2.7459E-27 | 0.259383218 | 0.148897394 | 1.143060118 | 4.07846E-06 | 3.58482E-05 | 0.47450793 | 0.2148563 |
| RhoL | 0.892879815 | 0.000175371 | 0.000842136 | 0.351791531 | 0.189453125 | 1.331208787 | 1.373E-108 | 3.3558E-107 | 0.610805642 | 0.242755604 | 0.964525029 | 0.000624869 | 0.00403804 | 0.371634175 | 0.190442853 |
| msi | 2.428850585 | 7.50244E-69 | 5.6262E-66 | 8.140065147 | 1.51171875 | 2.650616561 | 0 | 0 | 1.590724361 | 0.253326043 | 2.080536543 | 2.182E-262 | 1.2668E-259 | 17.45068437 | 4.125803662 |
| dsx | 2.310895014 | 1.17696E-23 | 6.38042E-22 | 1.172638436 | 0.236328125 | 1.056322855 | 3.8464E-54 | 4.34968E-53 | 0.306239541 | 0.147257153 | 1.205397918 | 3.07967E-68 | 2.65405E-66 | 5.172453077 | 2.243032827 |
| CRMP | 2.606660621 | 8.44678E-11 | 1.29273E-09 | 0.273615635 | 0.044921875 | 1.606985614 | 6.24801E-32 | 4.42561E-31 | 0.099928281 | 0.032804811 | 1.071430099 | 6.46242E-12 | 9.45451E-11 | 0.823968678 | 0.392083083 |
| btsz | 2.402801671 | 2.6458E-54 | 9.92064E-52 | 8.521172638 | 1.611328125 | 1.039514054 | 1.2718E-232 | 8.6211E-231 | 2.154434616 | 1.048113723 | 2.234958303 | 1.592E-273 | 1.3509E-270 | 12.80124245 | 2.719334513 |
| CG5191 | 1.927729714 | 4.02412E-07 | 3.4921E-06 | 0.237785016 | 0.0625 | 2.699213791 | 5.63882E-95 | 1.15949E-93 | 0.169256514 | 0.0260616 | 1.428674761 | 1.21235E-15 | 2.19958E-14 | 0.81263732 | 0.301871982 |
| CG45263 | 2.668003141 | 1.69162E-09 | 2.18406E-08 | 0.918566775 | 0.14453125 | 4.797349317 | 8.7413E-259 | 7.0286E-257 | 0.562514941 | 0.020229634 | 3.82597023 | 9.213E-117 | 1.4729E-114 | 3.352633825 | 0.236403728 |
| Mrp4 | 3.134795307 | 9.63189E-20 | 3.48102E-18 | 0.51465798 | 0.05859375 | 1.319586659 | 1.13453E-56 | 1.32673E-55 | 0.229261296 | 0.091853472 | 2.745966281 | 8.18341E-15 | 1.42608E-13 | 0.315520073 | 0.04703366 |
| CG34347 | 1.839748973 | 3.07249E-23 | 1.62643E-21 | 3.201954397 | 0.89453125 | 1.222010822 | 5.68997E-86 | 1.03817E-84 | 0.420033469 | 0.180061965 | 0.737406472 | 6.46878E-27 | 1.97119E-25 | 10.51863087 | 6.309250164 |
| Invadolysin | 0.791843962 | 0.00120902 | 0.004627806 | 0.35504886 | 0.205078125 | 1.127479836 | 1.47727E-25 | 8.89644E-25 | 0.119053311 | 0.054492437 | 1.028825158 | 0.000532006 | 0.003505708 | 0.225944112 | 0.110737265 |
| Apc | 0.865455204 | 0.000198416 | 0.000939266 | 0.423452769 | 0.232421875 | 1.168445245 | 3.123E-44 | 2.93401E-43 | 0.230217547 | 0.102423911 | 2.329957776 | 0.00386537 | 0.021014736 | 0.040653179 | 0.008085508 |
| CG42232 | 1.699635138 | 3.84537E-23 | 2.02365E-21 | 2.423452769 | 0.74609375 | 1.902057123 | 0 | 0 | 1.256036338 | 0.336067068 | 1.085776197 | 9.53644E-12 | 1.38234E-10 | 0.724393623 | 0.341289883 |
| Spc25 | 1.761491975 | 2.40198E-13 | 4.85739E-12 | 0.602605863 | 0.177734375 | 1.711225628 | 9.0722E-157 | 3.5376E-155 | 0.473822615 | 0.144705668 | #NUM! | 0.008258629 | 0.040543365 | 0.01734094 | 0 |
| E(spl)m3-HLH | 1.525411653 | 1.62803E-33 | 1.95342E-31 | 15.10749186 | 5.248046875 | 1.020814861 | 5.8726E-219 | 3.6227E-217 | 6.061678221 | 2.987424822 | 1.027403306 | 3.66063E-21 | 8.72148E-20 | 2.120430198 | 1.040266871 |
| fru | 2.089486209 | 1.15255E-63 | 6.48238E-61 | 11.92833876 | 2.802734375 | 1.434818843 | 0 | 0 | 1.777193402 | 0.65737197 | 1.593749517 | 4.1787E-222 | 1.8438E-219 | 45.15173807 | 14.95918946 |
| tacc | 1.569757798 | 6.45273E-23 | 3.28068E-21 | 3.820846906 | 1.287109375 | 1.305470764 | 1.6524E-286 | 1.6054E-284 | 1.52163519 | 0.61563696 | 1.095542713 | 1.25627E-58 | 9.17742E-57 | 5.478384906 | 2.563665294 |
| Sulf1 | 1.295900608 | 2.81272E-07 | 2.52359E-06 | 0.517915309 | 0.2109375 | 1.793220254 | 3.70204E-72 | 5.60546E-71 | 0.206550323 | 0.059595407 | 1.062394467 | 1.26486E-17 | 2.49155E-16 | 2.018048712 | 0.966315717 |
| CG15546 | #NUM! | 7.54878E-05 | 0.000399362 | 0.091205212 | 0 | 4.877527052 | 9.1925E-215 | 5.6113E-213 | 0.444656945 | 0.015126663 | #NUM! | 1.09192E-05 | 9.24403E-05 | 0.049600709 | 0 |
| jumu | 1.589954578 | 1.05252E-23 | 5.74041E-22 | 2.32247557 | 0.771484375 | 1.834758238 | 1.546E-185 | 7.4483E-184 | 0.546736792 | 0.153271369 | 1.237977479 | 2.01578E-05 | 0.000164834 | 0.163003128 | 0.069107882 |
| Esyt2 | 3.662177828 | 2.1697E-43 | 4.88128E-41 | 2.175895765 | 0.171875 | 2.491892536 | 8.7553E-139 | 2.9417E-137 | 0.280898876 | 0.049936213 | 2.95757394 | 2.47558E-12 | 3.71539E-11 | 0.179156624 | 0.023062926 |
| Cep135 | 2.349834703 | 2.62049E-15 | 6.39073E-14 | 0.527687296 | 0.103515625 | 1.901792365 | 3.1872E-102 | 7.1875E-101 | 0.265598852 | 0.071077091 | 1.120615304 | 0.000908293 | 0.005683142 | 0.153617192 | 0.070648184 |
| Rbp6 | 2.340569657 | 1.5231E-06 | 1.1675E-05 | 0.267100977 | 0.052734375 | 1.772301362 | 5.16035E-51 | 5.4995E-50 | 0.149414296 | 0.043739748 | 1.459873249 | 1.04905E-32 | 3.79414E-31 | 1.978276996 | 0.719153276 |
| CG32354 | 1.534371761 | 1.29761E-06 | 1.01014E-05 | 0.32247557 | 0.111328125 | 1.677221135 | 2.2766E-60 | 2.86331E-59 | 0.254123835 | 0.079460543 | 1.818873013 | 7.43012E-11 | 1.01942E-09 | 0.353879258 | 0.100304351 |
| LanB2 | 0.853963284 | 3.88959E-11 | 6.23929E-10 | 4.045602606 | 2.23828125 | 0.805301119 | 1.1983E-183 | 5.6564E-182 | 2.929715515 | 1.67650811 | 0.80635719 | 1.39891E-31 | 4.83741E-30 | 8.103808615 | 4.633951398 |
| ko | 3.033985122 | 1.517E-50 | 5.46059E-48 | 3.423452769 | 0.41796875 | 1.824124021 | 5.0686E-123 | 1.4378E-121 | 0.328472388 | 0.092764717 | 2.505237771 | 3.0667E-128 | 5.6381E-126 | 4.057121757 | 0.714605453 |
| SP1173 | 2.567980153 | 1.97277E-10 | 2.85849E-09 | 0.938110749 | 0.158203125 | 4.282250508 | 1.01751E-46 | 9.97746E-46 | 0.063829787 | 0.003280481 | 1.394736368 | 2.14953E-05 | 0.000175251 | 0.210465517 | 0.08004302 |
| fax | 1.694157259 | 5.8704E-40 | 1.07812E-37 | 5.28990228 | 1.634765625 | 1.544387399 | 0 | 0 | 3.60148219 | 1.23473665 | 0.943313098 | 4.82601E-20 | 1.08866E-18 | 1.929544845 | 1.003435216 |
| dsx-c73A | 1.570177709 | 1.32516E-06 | 1.02891E-05 | 0.237785016 | 0.080078125 | 4.554709927 | 2.3193E-231 | 1.5452E-229 | 0.411188142 | 0.017495899 | 0.88384671 | 1.77648E-17 | 3.48689E-16 | 4.647385856 | 2.51851395 |
| CG14995 | 1.853382372 | 1.2719E-09 | 1.66849E-08 | 0.423452769 | 0.1171875 | 1.097488359 | 2.29187E-14 | 9.67447E-14 | 0.069806359 | 0.032622562 | 0.807666612 | 1.91272E-06 | 1.73513E-05 | 0.908748717 | 0.519172803 |
| sfl | 2.465439996 | 1.12187E-41 | 2.24348E-39 | 3.80781759 | 0.689453125 | 1.383457157 | 1.1176E-183 | 5.297E-182 | 0.771694956 | 0.295790049 | 1.389759967 | 3.1696E-107 | 4.3705E-105 | 9.270148536 | 3.537750948 |
| CG32485 | 2.326590741 | 4.89391E-20 | 1.83501E-18 | 1.263843648 | 0.251953125 | 3.262845355 | 0 | 0 | 1.443222568 | 0.150355385 | 2.880670814 | 0.000386936 | 0.002610577 | 0.052869709 | 0.007178581 |
| CG13252 | 1.103720138 | 4.56544E-26 | 2.99886E-24 | 6.006514658 | 2.794921875 | 1.547471345 | 0 | 0 | 2.087736075 | 0.714233643 | 1.419635656 | 3.3395E-121 | 5.8473E-119 | 12.15195559 | 4.542482452 |
| Baldspot | 0.924526445 | 7.0238E-07 | 5.79351E-06 | 0.778501629 | 0.41015625 | 0.702907909 | 9.58704E-23 | 5.37381E-22 | 0.253645709 | 0.155822854 | 0.58078532 | 0.000339318 | 0.002309079 | 0.715525995 | 0.478400488 |
| Hexo1 | 1.130222577 | 2.29945E-07 | 2.09441E-06 | 0.615635179 | 0.28125 | 0.919126563 | 8.00072E-35 | 6.06504E-34 | 0.253645709 | 0.134135229 | 1.554427794 | 2.20455E-07 | 2.22289E-06 | 0.233531608 | 0.079508998 |

| | | | | | | | | | | | | | | | |
|---|---|---|---|---|---|---|---|---|---|---|---|---|---|---|---|
| ome | 1.333557934 | 4.30676E-13 | 8.42533E-12 | 2.426710098 | 0.962890625 | 1.027710749 | 4.57164E-32 | 3.25401E-31 | 0.191728425 | 0.094040459 | 1.47794859 | 1.18503E-63 | 9.40436E-62 | 3.773977411 | 1.354853816 |
| CG32243 | 2.04635002 | 1.27232E-14 | 2.92828E-13 | 0.693811075 | 0.16796875 | 1.12382379 | 3.9091E-118 | 1.0682E-116 | 0.695433899 | 0.319117915 | #NUM! | 0.005231768 | 0.027652916 | 0.019820582 | 0 |
| sti | 4.222104678 | 3.39257E-28 | 2.70175E-26 | 0.765472313 | 0.041015625 | 3.212592424 | 0 | 0 | 0.554147741 | 0.059777656 | 3.049190764 | 2.94186E-15 | 5.24259E-14 | 0.201716758 | 0.024369359 |
| pbl | 1.826441829 | 3.58755E-16 | 9.41235E-15 | 1.094462541 | 0.30859375 | 2.750422515 | 1.9426E-209 | 1.1212E-207 | 0.434138178 | 0.064516129 | 2.170646675 | 7.94432E-36 | 3.23372E-34 | 0.733213064 | 0.162854752 |
| olf413 | 4.197336773 | 0.000266983 | 0.001222067 | 0.107491857 | 0.005859375 | 2.495816238 | 3.17265E-06 | 9.14459E-06 | 0.010279704 | 0.00182249 | 2.40165911 | 6.92021E-08 | 7.38268E-07 | 0.176002347 | 0.033307883 |
| CG7272 | 1.451600969 | 0.000543786 | 0.002290978 | 0.133550489 | 0.048828125 | 1.412505081 | 7.72414E-68 | 1.10227E-66 | 0.280898876 | 0.105522143 | #NUM! | 0.005589249 | 0.029303712 | 0.019990733 | 0 |
| CG32447 | #NUM! | 0.006998001 | 0.021225146 | 0.042345277 | 0 | 2.797928078 | 6.3114E-116 | 1.6762E-114 | 0.226870667 | 0.032622562 | 4.85201338 | 5.48316E-19 | 1.15871E-17 | 0.262931735 | 0.009104193 |
| fz2 | 3.846934562 | 1.13788E-69 | 9.3089E-67 | 5.817589577 | 0.404296875 | 2.765900104 | 0 | 0 | 1.532153956 | 0.225259705 | 2.511781618 | 3.1484E-271 | 2.3153E-268 | 11.19482565 | 1.962888925 |
| Tet | 4.395204597 | 1.65935E-33 | 1.9648E-31 | 1.931596091 | 0.091796875 | 2.177829935 | 8.9327E-148 | 3.2144E-146 | 0.350466173 | 0.077455805 | 2.672158288 | 8.0793E-191 | 2.7007E-188 | 5.261757724 | 0.825526558 |
| CG17732 | 1.247579528 | 5.94115E-05 | 0.000323439 | 0.273615635 | 0.115234375 | 1.010679448 | 3.80273E-22 | 2.08837E-21 | 0.127420512 | 0.063240386 | 1.010570986 | 0.001666328 | 0.009792894 | 0.224316807 | 0.111339594 |
| miple2 | 1.606157664 | 9.20316E-33 | 9.74344E-31 | 10.56026059 | 3.46875 | 1.333689465 | 0 | 0 | 8.066220416 | 3.200291598 | 1.550062667 | 4.32025E-86 | 4.62686E-84 | 4.495929855 | 1.535338601 |
| CG1136 | 3.05983325 | 1.43065E-20 | 5.64666E-19 | 2.491856678 | 0.298828125 | 4.055520139 | 0 | 0 | 0.748505857 | 0.045015491 | 3.752699651 | 2.0705E-111 | 3.1288E-109 | 2.911788168 | 0.216015352 |
| LanA | 1.222722092 | 4.58773E-23 | 2.38642E-21 | 7.703583062 | 3.30078125 | 0.836922398 | 9.0538E-252 | 6.9446E-250 | 5.09729859 | 2.853654091 | 0.621853635 | 2.44219E-12 | 3.67027E-11 | 4.95660327 | 3.220976544 |
| Ac76E | 1.69883559 | 2.10258E-10 | 3.02737E-09 | 0.830618893 | 0.255859375 | 2.234423257 | 4.81137E-58 | 5.76523E-57 | 0.122639254 | 0.0260616 | 1.861004563 | 1.53224E-43 | 7.89822E-42 | 1.787320922 | 0.492021798 |
| melt | 1.725305118 | 2.6296E-28 | 2.11283E-26 | 2.228013029 | 0.673828125 | 1.184130105 | 5.60259E-71 | 8.35302E-70 | 0.332536457 | 0.146345909 | 1.730688057 | 1.79812E-63 | 1.41679E-61 | 2.957062879 | 0.890987356 |
| Rbfox1 | 0.820496333 | 6.79952E-17 | 1.91215E-15 | 10.71335505 | 6.06640625 | 0.677751451 | 3.51106E-72 | 5.3232E-71 | 1.041357877 | 0.650993257 | 0.802057817 | 1.15407E-95 | 1.44665E-93 | 15.39294189 | 8.828322074 |
| CG10512 | 5.361341803 | 5.59981E-49 | 1.73768E-46 | 2.970684039 | 0.072265625 | 1.762911077 | 1.8518E-137 | 6.1336E-136 | 0.981592159 | 0.289229087 | 2.151383912 | 4.03862E-47 | 2.23869E-45 | 1.403168023 | 0.315848333 |
| form3 | 2.454112189 | 9.83651E-11 | 1.48273E-09 | 0.299674267 | 0.0546875 | 1.983496836 | 3.53142E-76 | 5.60175E-75 | 0.196031556 | 0.049571715 | 1.690675215 | 1.78046E-24 | 4.86145E-23 | 0.863845136 | 0.267603593 |
| ftz-f1 | 1.359393531 | 7.66576E-26 | 4.92744E-24 | 7.166123779 | 2.79296875 | 1.290796954 | 2.54301E-71 | 3.81582E-70 | 0.340664595 | 0.139238199 | 0.754598742 | 3.84576E-20 | 8.72894E-19 | 4.218364791 | 2.500272116 |
| CG12768 | 5.261467111 | 9.07587E-06 | 5.9922E-05 | 0.074918567 | 0.001953125 | 4.168460851 | 1.22228E-58 | 1.48135E-57 | 0.078651685 | 0.004373975 | 3.284131009 | 1.21864E-07 | 1.26819E-06 | 0.109520306 | 0.011242754 |
| p130CAS | 1.233070985 | 4.60542E-17 | 1.32834E-15 | 2.13029316 | 0.90625 | 1.125336687 | 1.84578E-88 | 3.47656E-87 | 0.480277313 | 0.220156734 | 0.734941256 | 2.0706E-18 | 4.21416E-17 | 3.274782469 | 1.967628451 |
| bab2 | 2.853382372 | 0.000421411 | 0.001827602 | 0.084690554 | 0.01171875 | 2.561404579 | 1.02651E-18 | 5.06694E-18 | 0.032273488 | 0.005467469 | 1.222229567 | 1.27592E-06 | 1.17681E-05 | 0.258255271 | 0.110693325 |
| Cyp305a1 | 1.372111174 | 6.69595E-08 | 6.73261E-07 | 0.48534202 | 0.1875 | 1.301394449 | 2.37801E-97 | 5.09654E-96 | 0.55653837 | 0.225806452 | 1.155793836 | 8.59524E-11 | 1.17344E-09 | 0.803341495 | 0.360555053 |
| corn | 3.345235468 | 1.31022E-20 | 5.24029E-19 | 0.635179153 | 0.0625 | 1.864250344 | 1.1399E-130 | 3.4881E-129 | 0.396127181 | 0.108802624 | 3.482695701 | 1.2843E-51 | 7.87059E-50 | 0.908948763 | 0.08130992 |
| Nrt | 1.790646617 | 8.99934E-11 | 1.37031E-09 | 0.635179153 | 0.18359375 | 2.190381935 | 4.6353E-97 | 9.8439E-96 | 0.22376285 | 0.049024968 | 1.50216961 | 8.84306E-13 | 1.37391E-11 | 0.486500211 | 0.171745324 |
| CG3961 | 1.639724761 | 5.05788E-05 | 0.000280962 | 0.231270358 | 0.07421875 | 0.648254993 | 3.0854E-18 | 1.50074E-17 | 0.263925412 | 0.168398032 | 1.021582403 | 1.43622E-06 | 1.3167E-05 | 0.557255108 | 0.27449037 |
| fng | 2.438763591 | 2.58485E-60 | 1.22427E-57 | 5.61237785 | 1.03515625 | 1.753521971 | 0 | 0 | 1.120248625 | 0.33223984 | 1.357619127 | 7.17447E-93 | 8.50985E-91 | 7.33826096 | 2.863578362 |
| Spc105R | 3.116416778 | 7.06936E-20 | 2.60726E-18 | 0.592833876 | 0.068359375 | 2.488247433 | 5.4783E-209 | 3.1464E-207 | 0.409036577 | 0.072899581 | 3.772029819 | 0.001155138 | 0.007055551 | 0.021238219 | 0.001554619 |
| Cad74A | 1.19537792 | 9.33863E-06 | 6.15215E-05 | 0.299674267 | 0.130859375 | 1.588837809 | 9.81839E-35 | 7.42365E-34 | 0.104709539 | 0.03480955 | 1.016451891 | 2.90432E-07 | 2.89148E-06 | 0.540554952 | 0.267212852 |
| bab1 | 5.322867655 | 0.007157106 | 0.021649344 | 0.078175896 | 0.001953125 | 2.279004849 | 1.02278E-05 | 2.84663E-05 | 0.008845326 | 0.00182249 | 0.794630282 | 7.08805E-05 | 0.000539975 | 0.52055257 | 0.300093815 |
| CG32845 | 4.545260077 | 0.00244985 | 0.008482571 | 0.045602606 | 0.001953125 | 3.166773291 | 2.60126E-15 | 1.13716E-14 | 0.021276596 | 0.002369236 | 2.888135795 | 2.61346E-05 | 0.000210278 | 0.080929266 | 0.01093176 |
| dlp | 1.511396625 | 1.1407E-12 | 2.12971E-11 | 1.302931596 | 0.45703125 | 1.096691121 | 4.20468E-80 | 7.13572E-79 | 0.497728903 | 0.232731912 | 0.885593744 | 4.96105E-34 | 1.92019E-32 | 4.636631363 | 2.509644969 |
| Pura | 1.554459014 | 1.04827E-09 | 1.39341E-08 | 0.768729642 | 0.26171875 | 2.467523759 | 5.06995E-94 | 1.02622E-92 | 0.195553431 | 0.035356297 | 1.318077718 | 3.54073E-29 | 1.13211E-27 | 1.630327167 | 0.653873649 |
| nuf | 2.474576291 | 3.55212E-49 | 1.14162E-46 | 5.319218241 | 0.95703125 | 1.943960602 | 4.6058E-238 | 3.2349E-236 | 0.766435573 | 0.199198105 | 1.077434794 | 2.13038E-50 | 1.25001E-48 | 7.262103293 | 3.441297591 |
| CG10479 | 2.06844638 | 1.65748E-17 | 5.0221E-16 | 1.368078176 | 0.326171875 | 1.853521465 | 6.1875E-108 | 1.4905E-106 | 0.330623954 | 0.091488974 | 1.495440013 | 1.71452E-18 | 3.5154E-17 | 1.120852017 | 0.397535554 |
| Klp61F | 2.457797235 | 5.7498E-17 | 1.64262E-15 | 0.912052117 | 0.166015625 | 2.514793637 | 0 | 0 | 0.720774564 | 0.126116275 | 2.805184944 | 1.12919E-05 | 9.53027E-05 | 0.069587837 | 0.009956083 |
| CG32264 | 1.403711012 | 1.94528E-27 | 1.47106E-25 | 6.48534202 | 2.451171875 | 1.230677353 | 1.5632E-219 | 9.6943E-218 | 1.344250538 | 0.572808456 | 0.691756738 | 9.78944E-33 | 3.56394E-31 | 10.95508253 | 6.782286412 |
| Exn | 2.388159116 | 9.48136E-23 | 4.74016E-21 | 1.104234528 | 0.2109375 | 0.605163091 | 7.28615E-10 | 2.54873E-09 | 0.125029883 | 0.082194277 | 0.700795259 | 9.00175E-07 | 8.43656E-06 | 1.760686962 | 1.083232682 |
| fz | 3.255574543 | 1.50023E-39 | 2.59626E-37 | 2.38762215 | 0.25 | 1.450718781 | 1.97569E-21 | 1.05711E-20 | 0.08319388 | 0.030435575 | 2.480569134 | 2.296E-264 | 1.4071E-261 | 6.623035972 | 1.186673921 |
| Tollo | 1.266153264 | 1.16251E-18 | 3.8603E-17 | 2.071661238 | 0.861328125 | 1.442492744 | 1.6083E-119 | 4.4329E-118 | 0.509203921 | 0.187351923 | 1.307825655 | 8.67928E-65 | 6.93776E-63 | 5.076286096 | 2.050460178 |
| Fili | 2.413902732 | 1.14117E-22 | 5.64253E-21 | 0.905537459 | 0.169921875 | 1.335172137 | 7.59148E-48 | 7.63668E-47 | 0.19220655 | 0.076180062 | 1.199243983 | 2.198E-42 | 1.07761E-40 | 4.082009535 | 1.777729191 |
| chn | 2.723047196 | 7.88609E-27 | 5.58794E-25 | 1.573289902 | 0.23828125 | 3.560537835 | 2.1535E-222 | 1.3646E-220 | 0.397800622 | 0.033716056 | 2.806222515 | 1.29532E-24 | 3.55439E-23 | 0.432460293 | 0.061828554 |
| CG30403 | 1.999460632 | 8.47056E-27 | 5.9552E-25 | 1.905537459 | 0.4765625 | 0.581272915 | 2.40517E-35 | 1.84381E-34 | 0.863016973 | 0.576817933 | #NUM! | 0.005666041 | 0.0295658 | 0.020230984 | 0 |
| edl | 3.13122759 | 7.55231E-40 | 1.33261E-37 | 3.508143322 | 0.400390625 | 2.127493212 | 3.6239E-165 | 1.4671E-163 | 0.603633756 | 0.138144706 | 1.10741265 | 2.68244E-10 | 3.49764E-09 | 0.960751505 | 0.445909421 |
| jbug | 1.119862707 | 6.9067E-09 | 8.09289E-08 | 1.638436482 | 0.75390625 | 0.904570783 | 5.90228E-52 | 6.44936E-51 | 0.498446091 | 0.266265719 | 1.206325074 | 4.33211E-40 | 2.00788E-38 | 5.076651581 | 2.20007419 |
| rgr | 1.874277596 | 1.91424E-15 | 4.75864E-14 | 1.661237785 | 0.453125 | 2.981474156 | 7.4624E-260 | 6.0842E-258 | 0.686588571 | 0.08693275 | 2.503885275 | 1.53366E-31 | 5.27033E-30 | 0.708750633 | 0.124953633 |
| Spn42Dc | 4.737905155 | 0.000225597 | 0.001055171 | 0.052117264 | 0.001953125 | 2.344886884 | 3.7523E-44 | 3.51673E-43 | 0.109251733 | 0.021505376 | 3.008145832 | 0.004522797 | 0.024242457 | 0.037913811 | 0.004712543 |
| Tret1-1 | 0.770729002 | 3.49852E-09 | 4.28926E-08 | 1.592833876 | 0.93359375 | 1.275585216 | 1.1502E-103 | 2.6347E-102 | 0.508725795 | 0.210133042 | 1.230922504 | 6.3775E-51 | 3.82338E-49 | 5.937047459 | 2.529448984 |
| uzip | 1.653292927 | 1.13107E-14 | 2.62334E-13 | 0.970684039 | 0.30859375 | 1.360794038 | 4.87299E-73 | 7.45592E-72 | 0.322495816 | 0.125569528 | 1.69853656 | 2.25299E-61 | 1.71398E-59 | 2.457477839 | 0.757145171 |
| Treh | 1.76893205 | 6.15089E-17 | 1.75164E-15 | 1.071661238 | 0.314453125 | 1.200281778 | 1.56168E-38 | 1.28767E-37 | 0.178819029 | 0.077820303 | 0.877981587 | 1.37553E-09 | 1.69536E-08 | 1.385231851 | 0.753743666 |
| CG42566 | 1.460446165 | 5.21341E-23 | 2.68089E-21 | 15.66775244 | 5.693359375 | 2.309495717 | 0 | 0 | 9.616065025 | 1.939857846 | 1.296072925 | 5.5371E-27 | 1.69196E-25 | 1.504281483 | 0.612593356 |
| otk | 2.717727273 | 7.58557E-09 | 8.85376E-08 | 0.231270358 | 0.03515625 | 1.704069808 | 8.25533E-34 | 6.1227E-33 | 0.092039206 | 0.028248588 | 3.511695193 | 8.61503E-48 | 4.82398E-46 | 0.889468135 | 0.077983874 |
| so | #NUM! | 0.000658438 | 0.002716773 | 0.048859935 | 0 | 0.909946667 | 0.004241461 | 0.009323378 | 0.012670332 | 0.006743211 | 2.592878626 | 6.84346E-05 | 0.000522062 | 0.083951016 | 0.013915273 |
| LRP1 | 2.280542246 | 1.49742E-25 | 9.22966E-24 | 1.859934853 | 0.3828125 | 1.143060153 | 6.84364E-22 | 3.72677E-21 | 0.11068611 | 0.050118462 | 0.876243213 | 4.40479E-16 | 8.1389E-15 | 1.721668488 | 0.937937802 |
| CG12769 | 0.785210869 | 0.004279922 | 0.01390936 | 0.302931596 | 0.17578125 | 0.803457277 | 1.54052E-05 | 4.24208E-05 | 0.037532871 | 0.021505376 | 0.901706383 | 0.000209907 | 0.0014786 | 0.613880312 | 0.328581446 |
| cnn | 1.512138007 | 9.98854E-10 | 1.33363E-08 | 0.596091205 | 0.208984375 | 1.806517077 | 4.46211E-66 | 6.1933E-65 | 0.179775281 | 0.051394204 | 2.020592454 | 0.001456206 | 0.008682503 | 0.046484993 | 0.011456549 |
| nord | 1.391299381 | 6.75591E-13 | 1.30465E-11 | 1.019543974 | 0.388671875 | 2.437698603 | 0 | 0 | 1.205594071 | 0.22252597 | 3.569188377 | 8.8779E-240 | 4.8966E-237 | 6.261176995 | 0.527500917 |
| LBR | 0.770205647 | 3.23152E-05 | 0.000188957 | 0.882736156 | 0.517578125 | 1.188689942 | 2.08318E-77 | 3.35467E-76 | 0.415013148 | 0.182066703 | 1.009220128 | 0.000157079 | 0.001127967 | 0.281796794 | 0.1400008 |
| mam | 1.712495991 | 3.34498E-30 | 2.98035E-28 | 4.583061889 | 1.3984375 | 1.565643351 | 6.082E-237 | 4.2461E-235 | 0.855606024 | 0.289046838 | 1.285305553 | 5.05091E-87 | 5.57166E-85 | 9.379087723 | 3.848088689 |
| mim | 1.626621821 | 4.44764E-21 | 1.85298E-19 | 1.381107492 | 0.447265625 | 1.328533233 | 1.3814E-133 | 4.4124E-132 | 0.557494621 | 0.221979224 | 1.042594258 | 2.91462E-14 | 4.93115E-13 | 1.224758876 | 0.594563782 |
| kcc | 1.421782681 | 6.74821E-11 | 1.04883E-09 | 0.811074919 | 0.302734375 | 1.785782385 | 4.1763E-130 | 1.2713E-128 | 0.412861583 | 0.119737562 | 0.839222776 | 1.03847E-06 | 9.65879E-06 | 0.650828507 | 0.363777085 |
| GLaz | 1.757270479 | 3.94496E-07 | 3.43999E-06 | 0.244299674 | 0.072265625 | 1.625360411 | 7.0957E-113 | 1.8142E-111 | 0.381783409 | 0.123747038 | 1.052178701 | 8.95883E-07 | 8.40347E-06 | 0.380994209 | 0.183730402 |
| CG8389 | 1.098652498 | 0.000100011 | 0.000516056 | 0.368078176 | 0.171875 | 0.847409508 | 1.89318E-24 | 1.10751E-23 | 0.196748745 | 0.109349371 | #NUM! | 0.004226178 | 0.022752062 | 0.01501905 | 0 |
| CG5522 | 1.319134936 | 1.65334E-16 | 4.53609E-15 | 1.25732899 | 0.50390625 | 1.016077854 | 4.68478E-65 | 6.39577E-64 | 0.401386565 | 0.198469109 | 0.791903908 | 0.007473231 | 0.037488501 | 0.289169644 | 0.167018997 |

| | | | | | | | | | | | | | | | |
|---|---|---|---|---|---|---|---|---|---|---|---|---|---|---|---|
| HmgD | 1.436094192 | 3.17333E-32 | 3.24509E-30 | 33.12052117 | 12.24023438 | 1.560329568 | 0 | 0 | 26.45852259 | 8.971386915 | 1.156954679 | 0.002805694 | 0.015750435 | 0.149458073 | 0.067025693 |
| Asph | 2.121915758 | 1.99672E-18 | 6.48682E-17 | 3.596091205 | 0.826171875 | 1.036484141 | 1.4816E-97 | 3.18122E-96 | 2.835524743 | 1.382358301 | 2.112125868 | 1.82171E-31 | 6.22146E-30 | 1.018792717 | 0.235652793 |
| Gbp2 | 3.261467111 | 6.5366E-05 | 0.000352444 | 0.074918567 | 0.0078125 | 0.810083692 | 3.43965E-05 | 9.26592E-05 | 0.032273488 | 0.018407144 | 2.67977168 | 0.002107699 | 0.012128338 | 0.039219843 | 0.006120885 |
| scra | 2.108128807 | 5.75677E-24 | 3.21771E-22 | 1.583061889 | 0.3671875 | 2.164449028 | 0 | 0 | 1.134831461 | 0.253143794 | 2.072193823 | 0.004352452 | 0.023397612 | 0.036138633 | 0.008593681 |
| Strn-Mlck | 1.960297576 | 1.02121E-09 | 1.36146E-08 | 0.319218241 | 0.08203125 | 0.679636936 | 1.28936E-11 | 4.8602E-11 | 0.113554865 | 0.070894842 | 1.966046462 | 1.41683E-15 | 2.56214E-14 | 0.433777654 | 0.111026901 |
| Hil | 2.386997993 | 1.45637E-08 | 1.62001E-07 | 0.2247557 | 0.04296875 | 1.069551483 | 4.25623E-12 | 1.63666E-11 | 0.059287593 | 0.028248588 | 1.310362568 | 4.29215E-08 | 4.68779E-07 | 0.379562427 | 0.153046985 |
| CG8306 | 1.224953899 | 1.08507E-12 | 2.03428E-11 | 1.159609121 | 0.49609375 | 0.701147158 | 1.33618E-47 | 1.33492E-46 | 0.574229022 | 0.353198469 | 0.949382698 | 2.81593E-07 | 2.80855E-06 | 0.615024281 | 0.31849278 |
| Pka-R2 | 2.029593942 | 1.37758E-45 | 3.44356E-43 | 8.381107492 | 2.052734375 | 2.537089083 | 0 | 0 | 2.91847956 | 0.502824859 | 1.291313767 | 8.14249E-95 | 1.00921E-92 | 14.98302924 | 6.121748022 |
| Magi | 1.180658032 | 1.83375E-15 | 4.57118E-14 | 1.540716612 | 0.6796875 | 1.251819274 | 3.6681E-114 | 9.4825E-113 | 0.493903897 | 0.207399307 | 1.071974305 | 0.000131362 | 0.000954582 | 0.254461505 | 0.121039096 |
| CG8547 | 3.595204508 | 4.11627E-35 | 5.61247E-33 | 1.723127036 | 0.142578125 | 2.134064244 | 6.2358E-131 | 1.9183E-129 | 0.327994262 | 0.07472207 | 2.15618381 | 9.01039E-34 | 3.42737E-32 | 0.861460693 | 0.193267782 |
| CG13506 | 2.786268176 | 4.00113E-13 | 7.87881E-12 | 0.592833876 | 0.0859375 | 1.391479578 | 3.28916E-18 | 1.59386E-17 | 0.065981353 | 0.025150355 | 1.857673244 | 8.38515E-25 | 2.33577E-23 | 0.978255183 | 0.269921071 |
| CG9815 | 2.655442994 | 2.11899E-15 | 5.23868E-14 | 0.442996743 | 0.0703125 | 2.50156903 | 9.9622E-110 | 2.466E-108 | 0.234281616 | 0.041370512 | 1.852158945 | 2.72109E-20 | 6.24041E-19 | 0.802483297 | 0.222269874 |
| apt | 1.430868613 | 3.41573E-35 | 4.72894E-33 | 5.328990228 | 1.9765625 | 1.058344354 | 1.9983E-160 | 7.9517E-159 | 1.195553431 | 0.574084199 | 0.712064882 | 1.89046E-52 | 1.19164E-50 | 10.91926868 | 6.665621727 |
| pk | 2.630989951 | 0.001674042 | 0.00609167 | 0.084690554 | 0.013671875 | 2.940818169 | 3.1256E-33 | 2.28043E-32 | 0.05737509 | 0.007472207 | 2.722501231 | 4.00142E-24 | 1.07658E-22 | 0.554836218 | 0.084064042 |
| PRAS40 | 1.382996431 | 2.69784E-19 | 9.30187E-18 | 6.423452769 | 2.462890625 | 1.703587624 | 4.0396E-223 | 2.5736E-221 | 1.015060961 | 0.311645708 | 0.684201137 | 3.97266E-22 | 9.82564E-21 | 7.682582526 | 4.781257885 |
| shg | 1.412494051 | 8.62245E-24 | 4.76033E-22 | 8.35504886 | 3.138671875 | 0.755452287 | 9.5868E-212 | 5.7615E-210 | 3.830743486 | 2.269181702 | 0.685715217 | 8.16047E-59 | 6.00121E-57 | 18.55753337 | 11.53717291 |
| sli | 4.142019005 | 3.16088E-13 | 6.29309E-12 | 1 | 0.056640625 | 1.938374038 | 1.94162E-54 | 2.20422E-53 | 0.134114272 | 0.034991799 | 2.990592043 | 2.96541E-47 | 1.65209E-45 | 0.906740465 | 0.114084091 |
| Spn42Dd | 3.828291391 | 1.46125E-26 | 1.01152E-24 | 0.804560261 | 0.056640625 | 2.031246515 | 4.0568E-127 | 1.1884E-125 | 0.557972747 | 0.136504465 | 3.415634178 | 3.29528E-31 | 1.11847E-29 | 0.57947957 | 0.054303746 |
| CG6044 | 6.492792657 | 1.75001E-06 | 1.32117E-05 | 0.175895765 | 0.001953125 | 5.19309813 | 1.33177E-57 | 1.58602E-56 | 0.120009563 | 0.003280481 | 3.920059552 | 5.36992E-18 | 1.06924E-16 | 0.259144644 | 0.017119329 |
| stl | 1.541239558 | 4.86313E-12 | 8.54752E-11 | 3.501628664 | 1.203125 | 1.411778073 | 0 | 0 | 2.617977528 | 0.983962092 | 0.901002096 | 1.92679E-45 | 1.0368E-43 | 13.63350314 | 7.300940443 |
| CG34207 | 2.875408678 | 1.46149E-19 | 5.1175E-18 | 0.573289902 | 0.078125 | 2.101689718 | 4.25624E-94 | 8.63017E-93 | 0.221372221 | 0.051576453 | 1.837802769 | 9.21924E-06 | 7.87742E-05 | 0.181770927 | 0.050849965 |
| lbk | 1.609675123 | 3.47637E-27 | 2.54341E-25 | 3.188925081 | 1.044921875 | 1.570730372 | 1.307E-144 | 4.6039E-143 | 0.55653837 | 0.187351923 | 0.676022101 | 5.30142E-18 | 1.0575E-16 | 5.592727826 | 3.500424819 |
| CG13516 | 1.647139691 | 3.07264E-11 | 5.00013E-10 | 0.501628664 | 0.16015625 | 1.410121984 | 6.2695E-131 | 1.9236E-129 | 0.502749223 | 0.189174412 | 1.050776435 | 0.001260857 | 0.007633655 | 0.199634061 | 0.09636502 |
| CG30069 | 3.578300338 | 2.05761E-21 | 8.90215E-20 | 0.863192182 | 0.072265625 | 3.287142918 | 3.7604E-285 | 3.6233E-283 | 0.535500837 | 0.054856935 | 2.443251337 | 9.90893E-45 | 5.22992E-43 | 1.170164368 | 0.215156728 |
| CycB | 0.654085164 | 9.12008E-11 | 1.38634E-09 | 3.267100977 | 2.076171875 | 3.077679044 | 0 | 0 | 1.217069089 | 0.144158921 | 1.143328869 | 0.00610462 | 0.031393966 | 0.121121235 | 0.054833252 |
| whd | 1.30130909 | 7.6475E-12 | 1.31336E-10 | 2.055374593 | 0.833984375 | 0.732050584 | 1.05756E-24 | 6.25256E-24 | 0.246712885 | 0.148532896 | 1.008238111 | 1.25657E-10 | 1.68833E-09 | 0.946155483 | 0.47038406 |
| babos | 2.354062277 | 3.94106E-56 | 1.61207E-53 | 11.5732899 | 2.263671875 | 1.668202128 | 0 | 0 | 5.073392302 | 1.596318571 | 1.694873556 | 1.4234E-103 | 1.9384E-101 | 5.202707413 | 1.607021265 |
| Fbl6 | 1.314285268 | 1.34502E-13 | 2.80181E-12 | 1.364820847 | 0.548828125 | 1.378304189 | 6.07426E-95 | 1.24567E-93 | 0.390867798 | 0.150355385 | 0.821539552 | 1.13359E-12 | 1.75135E-11 | 1.649887287 | 0.933568583 |
| Fas2 | 3.094067252 | 7.363E-107 | 3.313E-103 | 30.83713355 | 3.611328125 | 2.755222623 | 0 | 0 | 9.70714798 | 1.437761983 | 2.692845437 | 0 | 0 | 88.47363644 | 13.68316701 |
| fs(1)Yb | 2.159862068 | 3.51653E-17 | 1.02412E-15 | 0.680781759 | 0.15234375 | 1.242307007 | 5.1599E-129 | 1.5308E-127 | 0.634233803 | 0.268088208 | 1.855594392 | 7.09395E-15 | 1.23819E-13 | 0.503314893 | 0.139075365 |
| CG3038 | 2.015889902 | 2.95198E-14 | 6.64122E-13 | 0.631921824 | 0.15625 | 1.107888289 | 3.54785E-88 | 6.65022E-87 | 0.488644514 | 0.226717696 | 1.399514437 | 0.001869035 | 0.010828425 | 0.065168702 | 0.024702633 |
| elav | 1.644219709 | 1.20742E-14 | 2.78604E-13 | 1.019543974 | 0.326171875 | 1.063753638 | 3.03581E-49 | 3.11846E-48 | 0.284962945 | 0.136322216 | 0.713138158 | 0.000417492 | 0.00279451 | 0.701006273 | 0.427608105 |
| CG43759 | 1.027484859 | 1.27486E-18 | 4.21783E-17 | 7.146579805 | 3.505859375 | 1.523161428 | 8.4498E-282 | 7.8187E-280 | 1.27707387 | 0.444322945 | 1.04440201 | 2.51606E-93 | 3.04996E-91 | 25.48553282 | 12.35655439 |
| boi | 1.352251462 | 1.33562E-05 | 8.53036E-05 | 0.403908795 | 0.158203125 | 1.014458342 | 5.12152E-24 | 2.95603E-23 | 0.142481473 | 0.070530344 | 0.80184383 | 5.92677E-10 | 7.4975E-09 | 1.218740845 | 0.699088763 |
| rut | 0.835515951 | 0.000266663 | 0.001221846 | 0.348534202 | 0.1953125 | 1.778842119 | 2.01102E-52 | 2.20569E-51 | 0.156347119 | 0.045562238 | 1.495701527 | 4.81479E-15 | 8.48433E-14 | 0.912398647 | 0.323544192 |
| CG14223 | 1.803196615 | 7.72571E-05 | 0.000408482 | 0.368078176 | 0.10546875 | 1.664827492 | 7.17838E-76 | 1.13559E-74 | 0.261773847 | 0.082558775 | 1.793358365 | 0.000557357 | 0.003657469 | 0.069200918 | 0.019964443 |
| CG11380 | 3.973121616 | 2.54433E-10 | 3.64013E-09 | 0.368078176 | 0.0234375 | 3.996341636 | 3.73072E-63 | 4.94278E-62 | 0.087257949 | 0.005467469 | 3.232985541 | 6.99346E-17 | 1.34165E-15 | 0.273027664 | 0.029038958 |
| Klp10A | 1.28899279 | 6.87752E-13 | 1.32529E-11 | 1.508143322 | 0.6171875 | 1.280323135 | 1.3686E-115 | 3.5938E-114 | 0.510399235 | 0.210133042 | 0.6351436 | 8.72482E-06 | 7.47813E-05 | 1.141430754 | 0.734940693 |
| CG15784 | 0.882380191 | 4.40158E-09 | 5.27428E-08 | 2.905537459 | 1.576171875 | 0.685776554 | 1.74581E-29 | 1.16245E-28 | 0.432703801 | 0.268999453 | 1.395516628 | 9.40411E-05 | 0.000700451 | 0.178938882 | 0.068016198 |
| bou | 0.660336601 | 8.3821E-06 | 5.56683E-05 | 1.71009772 | 1.08203125 | 0.791938963 | 6.8878E-162 | 2.7691E-160 | 2.283289505 | 1.318753417 | 0.722103147 | 6.64695E-08 | 7.09802E-07 | 1.52860682 | 0.926661428 |
| CG5004 | 1.781282217 | 2.01237E-27 | 1.49955E-25 | 2.309446254 | 0.671875 | 1.902295187 | 6.1262E-240 | 4.3819E-238 | 0.70714798 | 0.189174412 | 1.951551121 | 3.063E-168 | 8.6635E-166 | 6.896714026 | 1.783063429 |
| meso18E | 5.604153766 | 7.00305E-34 | 8.51628E-32 | 1.140065147 | 0.0234375 | 4.182411101 | 0 | 0 | 0.605546259 | 0.033351558 | 3.93153525 | 2.2363E-101 | 3.0084E-99 | 1.738337008 | 0.113926285 |
| CG15365 | 2.794417176 | 9.33776E-35 | 1.23574E-32 | 2.628664495 | 0.37890625 | 2.33822151 | 1.3344E-117 | 3.6014E-116 | 0.311498924 | 0.061600146 | 0.728431323 | 9.82805E-07 | 9.15652E-06 | 0.888940754 | 0.536528923 |
| CG42258 | 2.078942072 | 1.0769E-31 | 1.06495E-29 | 2.351791531 | 0.556640625 | 1.337189422 | 5.15985E-80 | 8.74399E-79 | 0.324169257 | 0.128303262 | 1.011989698 | 6.94762E-23 | 1.77406E-21 | 1.934506408 | 0.959248035 |
| Bx | 2.468945915 | 3.45632E-28 | 2.72837E-26 | 1.319218241 | 0.23828125 | 2.830959218 | 1.8327E-182 | 8.5812E-181 | 0.360506813 | 0.050665209 | 2.365025498 | 6.5191E-101 | 8.6641E-99 | 3.564529801 | 0.691924382 |
| dome | 1.42764654 | 4.12719E-14 | 9.05867E-13 | 1.859934853 | 0.69140625 | 1.29185222 | 3.3463E-186 | 1.6324E-184 | 1.003107817 | 0.409695644 | 0.597054162 | 7.85609E-08 | 8.3488E-07 | 1.963391367 | 1.298002908 |
| N | 1.295795179 | 2.59366E-33 | 2.99235E-31 | 24.76221498 | 10.0859375 | 0.93805507 | 0 | 0 | 9.738943342 | 5.083105522 | 0.675712833 | 3.98488E-66 | 3.25609E-64 | 61.78525599 | 38.67898058 |
| CG9132 | 1.699180637 | 9.70977E-35 | 1.25251E-32 | 3.951140065 | 1.216796875 | 1.496962826 | 2.2614E-268 | 1.9676E-266 | 1.203681568 | 0.426462548 | 0.78563466 | 8.24594E-13 | 1.28295E-11 | 1.888881338 | 1.095733848 |
| CG4615 | 2.172842211 | 6.33248E-08 | 6.38141E-07 | 0.237785016 | 0.052734375 | 1.599587773 | 4.1319E-17 | 1.94014E-16 | 0.048051638 | 0.015855659 | 1.679230179 | 5.09057E-05 | 0.000395173 | 0.135769046 | 0.042393777 |
| Flo2 | 1.586963257 | 5.83304E-19 | 1.97337E-17 | 1.771986971 | 0.58984375 | 0.835523108 | 3.36962E-34 | 2.51998E-33 | 0.284245757 | 0.159285584 | 0.836079732 | 1.14412E-22 | 2.90802E-21 | 3.470111078 | 1.943830081 |
| CG1764 | 2.039369147 | 4.64607E-15 | 1.12091E-13 | 0.570032573 | 0.138671875 | 1.243945342 | 4.55184E-85 | 8.25348E-84 | 0.417403777 | 0.176234737 | 0.721715838 | 0.005326907 | 0.028088488 | 0.312432062 | 0.189451252 |
| if | 3.786268176 | 2.70993E-12 | 4.89692E-11 | 0.296416938 | 0.021484375 | 3.653494771 | 8.37183E-88 | 1.55922E-86 | 0.135309586 | 0.010752688 | 2.788888915 | 3.02505E-53 | 1.95142E-51 | 1.228935801 | 0.177823837 |
| CG42797 | 3.166481921 | 2.60145E-54 | 9.92064E-52 | 4.068403909 | 0.453125 | 2.445457199 | 0 | 0 | 2.708104231 | 0.497175141 | 1.834356217 | 6.44516E-52 | 3.97187E-50 | 1.962986818 | 0.550454136 |
| kirre | 1.445724403 | 0.000655285 | 0.002707486 | 0.159609121 | 0.05859375 | 2.776143428 | 7.89858E-18 | 3.79125E-17 | 0.033707865 | 0.004920722 | 1.121041736 | 2.81084E-05 | 0.000225501 | 1.18609578 | 0.545321439 |
| wgn | 2.162926742 | 1.59598E-20 | 6.2717E-19 | 0.944625407 | 0.2109375 | 1.801531644 | 3.911E-139 | 1.3255E-137 | 0.391345924 | 0.112265354 | 0.882821291 | 3.63087E-11 | 5.08275E-10 | 1.589259873 | 0.861864969 |
| CG12065 | 1.533243112 | 6.0698E-14 | 1.30675E-12 | 0.876221498 | 0.302734375 | 2.181410435 | 3.7992E-161 | 1.5222E-159 | 0.39349749 | 0.086750501 | 0.960305643 | 2.99623E-12 | 4.4785E-11 | 1.230274346 | 0.632297063 |
| baz | 0.847191913 | 1.86817E-10 | 2.71594E-09 | 5.491856678 | 3.052734375 | 0.903433432 | 7.6966E-186 | 3.7234E-184 | 1.747788668 | 0.934390377 | 0.657963508 | 2.08544E-38 | 9.20178E-37 | 18.45239379 | 11.6946159 |
| ec | 1.276931371 | 3.22274E-09 | 3.97281E-08 | 1.107491857 | 0.45703125 | 0.589817201 | 2.41421E-27 | 1.52726E-26 | 0.407602199 | 0.270821943 | 1.071755279 | 1.53581E-20 | 3.58171E-19 | 2.146797503 | 1.021317079 |
| Nrg | 1.100600072 | 5.51473E-20 | 2.05921E-18 | 5.377850163 | 2.5078125 | 0.812174712 | 1.66701E-88 | 3.14492E-87 | 0.977289027 | 0.5565883 | 0.631930093 | 2.2111E-20 | 5.12232E-19 | 8.693589541 | 5.610082119 |
| CG34417 | 2.157016915 | 1.11028E-66 | 6.66092E-64 | 7.700325733 | 1.7265625 | 0.944216633 | 1.7724E-148 | 6.4374E-147 | 1.407602199 | 0.731547294 | 1.321356443 | 8.43852E-76 | 7.95601E-74 | 6.522393478 | 2.609991367 |
| ct | 3.10700422 | 4.7216E-138 | 4.2489E-134 | 29.06188925 | 3.373046875 | 2.8607944 | 0 | 0 | 2.801338752 | 0.385638783 | 3.102231122 | 0 | 0 | 98.54060202 | 11.47494441 |
| mud | 2.165586863 | 4.90283E-22 | 2.20603E-20 | 1.244299674 | 0.27734375 | 2.413315379 | 1.2977E-210 | 7.68E-209 | 0.445613196 | 0.083652269 | 3.337346787 | 0.001051526 | 0.006476484 | 0.041629216 | 0.004118668 |
| Axs | 1.907830156 | 2.58394E-08 | 2.76649E-07 | 0.263843648 | 0.0703125 | 0.992727162 | 5.09318E-20 | 2.61823E-19 | 0.117140808 | 0.058866412 | 2.355699946 | 0.000457261 | 0.003040413 | 0.052424678 | 0.010242346 |
| drl | 3.213638586 | 7.3727E-17 | 2.06688E-15 | 0.57980456 | 0.0625 | 2.44297309 | 3.84145E-72 | 5.809E-71 | 0.170451829 | 0.03134682 | 3.341985138 | 2.91066E-41 | 1.38394E-39 | 0.860457626 | 0.08485779 |

| | | | | | | | | | | | | | | | |
|---|---|---|---|---|---|---|---|---|---|---|---|---|---|---|---|
| CG13101 | 2.091542109 | 4.55561E-06 | 3.18539E-05 | 0.2247557 | 0.052734375 | 1.718741117 | 6.51828E-23 | 3.67134E-22 | 0.064786039 | 0.019682887 | 1.892094774 | 4.38991E-13 | 7.00797E-12 | 0.437814625 | 0.117954084 |
| CG10026 | 4.010923649 | 1.08762E-08 | 1.23736E-07 | 0.188925081 | 0.01171875 | 2.795201764 | 1.42569E-18 | 7.01059E-18 | 0.030360985 | 0.004373975 | 2.403794223 | 0.002501742 | 0.01416669 | 0.051363784 | 0.009706057 |
| piwi | 1.546061797 | 4.82514E-33 | 5.36067E-31 | 8.794788274 | 3.01171875 | 1.490884815 | 0 | 0 | 1.888596701 | 0.671951886 | 1.073411213 | 1.13686E-46 | 6.23918E-45 | 3.336703028 | 1.585581547 |
| CG9967 | 2.835515951 | 1.01625E-22 | 5.05261E-21 | 1.394136808 | 0.1953125 | 3.099908808 | 0 | 0 | 1.168778389 | 0.136322216 | 2.08280383 | 1.6382E-119 | 2.738E-117 | 4.370552587 | 1.031691666 |
| Idgf3 | 3.112188655 | 6.01515E-27 | 4.29606E-25 | 1.553745928 | 0.1796875 | 1.814305879 | 1.94426E-88 | 3.65614E-87 | 0.325603634 | 0.092582468 | 2.692434738 | 4.70292E-35 | 1.85942E-33 | 0.747323123 | 0.115612503 |
| eya | 1.570223847 | 1.55545E-48 | 4.51532E-46 | 7.45276873 | 2.509765625 | 1.932068494 | 0 | 0 | 3.053789147 | 0.800255149 | 2.403115797 | 0 | 0 | 20.24758796 | 3.827924439 |
| tsh | 1.795620652 | 7.06771E-07 | 5.82439E-06 | 0.332247557 | 0.095703125 | 2.757317371 | 1.2635E-197 | 6.8517E-196 | 0.42266316 | 0.062511391 | 2.861380192 | 4.36279E-54 | 2.86464E-52 | 1.317226394 | 0.181258824 |
| CadN | 2.663661183 | 1.9371E-103 | 5.8107E-100 | 19.30618893 | 3.046875 | 1.462720304 | 2.4884E-280 | 2.2666E-278 | 1.215634712 | 0.441042464 | 3.017358159 | 0 | 0 | 26.64883062 | 3.291264935 |
| Df31 | 1.581339661 | 7.60392E-47 | 1.95508E-44 | 69.10749186 | 23.09375 | 1.273811355 | 0 | 0 | 61.45374133 | 25.41516311 | 1.248111708 | 0.00548054 | 0.028779912 | 0.142793151 | 0.060115756 |
| CG31705 | 2.853382372 | 6.77875E-29 | 5.54563E-27 | 1.143322476 | 0.158203125 | 1.681060375 | 8.8225E-154 | 3.3505E-152 | 0.51661487 | 0.161108074 | 1.422749106 | 2.23682E-28 | 7.07003E-27 | 1.303657892 | 0.486265526 |
| Fas3 | 2.009928344 | 2.57603E-33 | 2.99235E-31 | 9.439739414 | 2.34375 | 1.059878557 | 1.8187E-251 | 1.3859E-249 | 2.707865169 | 1.298888281 | 1.119386128 | 1.5377E-131 | 2.9758E-129 | 28.58145586 | 13.15574799 |
| CG31886 | 1.270573148 | 2.41174E-07 | 2.19224E-06 | 0.48534202 | 0.201171875 | 1.650289398 | 1.03375E-53 | 1.16001E-52 | 0.16304088 | 0.051940951 | 1.431300809 | 2.27852E-15 | 4.06704E-14 | 0.783004527 | 0.29033527 |
| CG9932 | 2.377443046 | 3.34666E-80 | 3.76458E-77 | 9.915309446 | 1.908203125 | 2.050327092 | 0 | 0 | 2.355964619 | 0.568798979 | 2.002929328 | 0 | 0 | 28.34050473 | 7.070754743 |
| CG31710 | 0.928558385 | 1.06085E-08 | 1.20996E-07 | 1.684039088 | 0.884765625 | 1.736523922 | 1.6566E-276 | 1.4744E-274 | 1.022710973 | 0.306907235 | 1.07258808 | 1.06788E-30 | 3.54812E-29 | 3.576205007 | 1.700361338 |
| CG17544 | 3.569589406 | 4.35308E-31 | 4.05408E-29 | 0.973941368 | 0.08203125 | 1.683762211 | 5.316E-130 | 1.6057E-128 | 0.414535023 | 0.129032258 | 1.545759508 | 6.22834E-12 | 9.12415E-11 | 0.410810956 | 0.140709053 |
| lok | 0.688768376 | 2.48453E-07 | 2.25159E-06 | 2.068403909 | 1.283203125 | 1.139467367 | 2.5494E-116 | 6.8016E-115 | 0.670093235 | 0.304173501 | 1.759171346 | 0.000808403 | 0.00510538 | 0.103759958 | 0.030652538 |
| Sema1a | 2.696955526 | 6.37583E-31 | 5.8547E-29 | 1.697068404 | 0.26171875 | 1.335944901 | 1.63131E-73 | 2.5158E-72 | 0.328950514 | 0.130308001 | 1.553387224 | 4.6948E-111 | 6.9984E-109 | 5.899671107 | 2.010072176 |
| bdl | 5.036458741 | 2.41082E-47 | 6.57423E-45 | 1.794788274 | 0.0546875 | 3.163392237 | 2.0189E-271 | 1.7698E-269 | 0.509442984 | 0.056861673 | 3.827751711 | 2.0705E-123 | 3.7442E-121 | 2.307420115 | 0.162501956 |
| Spn28Dc | 4.159368923 | 1.23081E-11 | 2.07806E-10 | 0.244299674 | 0.013671875 | 3.94606843 | 1.22694E-63 | 1.63672E-62 | 0.08988764 | 0.005831966 | 2.662880736 | 8.37696E-08 | 8.85415E-07 | 0.137025312 | 0.021636839 |
| Nhe2 | 2.502501868 | 1.38353E-34 | 1.75358E-32 | 2.003257329 | 0.353515625 | 1.608423749 | 1.3877E-175 | 6.152E-174 | 0.631843175 | 0.207217059 | 2.146100408 | 1.71E-114 | 2.6567E-112 | 3.693513274 | 0.834447596 |
| ds | 1.960297576 | 0.009394665 | 0.027140478 | 0.045602606 | 0.01171875 | 0.629423003 | 0.000161796 | 0.000410529 | 0.040879751 | 0.026426098 | 1.188500426 | 5.02107E-05 | 0.000390328 | 0.212873675 | 0.093400173 |
| glu | 1.695911066 | 2.73619E-15 | 6.65486E-14 | 1.543973941 | 0.4765625 | 1.520805546 | 0 | 0 | 1.505617978 | 0.524694733 | 0.703602826 | 0.001050926 | 0.006476407 | 0.284604118 | 0.17475742 |
| Uvrag | 2.644460591 | 1.0508E-41 | 2.14912E-39 | 3.504885993 | 0.560546875 | 2.31721522 | 0 | 0 | 2.353334927 | 0.472207035 | 1.38027237 | 6.68982E-25 | 1.87665E-23 | 1.359470434 | 0.522235489 |
| neb | 1.124928278 | 1.24056E-05 | 8.01423E-05 | 0.38762215 | 0.177734375 | 1.282773318 | 7.74322E-47 | 7.63129E-46 | 0.192445613 | 0.079096045 | 0.955927303 | 4.88282E-05 | 0.000380384 | 0.386737353 | 0.199367025 |
| CG31808 | 4.690732034 | 6.39609E-39 | 1.08601E-36 | 3.429967427 | 0.1328125 | 2.898529217 | 0 | 0 | 1.012431269 | 0.135775469 | 1.822439243 | 2.1477E-16 | 4.02228E-15 | 0.497329657 | 0.14061624 |
| spi | 2.081582833 | 4.34444E-36 | 6.30575E-34 | 2.61237785 | 0.6171875 | 1.912287773 | 1.846E-204 | 1.0448E-202 | 0.62562754 | 0.166211044 | 0.842067565 | 1.11349E-18 | 2.30883E-17 | 2.703437193 | 1.508094961 |
| stai | 2.36830412 | 8.43224E-58 | 3.61342E-55 | 17.33550489 | 3.357421875 | 3.031853174 | 0 | 0 | 4.104948601 | 0.501913614 | 1.152766291 | 0.000700719 | 0.004478352 | 0.107080313 | 0.048160657 |
| dnt | 2.971395285 | 1.83575E-29 | 1.52962E-27 | 1.133550489 | 0.14453125 | 2.558465701 | 1.5545E-140 | 5.3149E-139 | 0.293091083 | 0.049753964 | 2.522799543 | 1.70529E-83 | 1.77463E-81 | 2.129590899 | 0.37055941 |
| cmet | 3.05983325 | 2.7208E-13 | 5.47751E-12 | 0.553745928 | 0.06640625 | 3.610888141 | 0 | 0 | 0.538847717 | 0.044104246 | 2.469495989 | 2.84737E-05 | 0.000228266 | 0.062461599 | 0.011277705 |
| E23 | 1.416662588 | 1.06188E-19 | 3.792E-18 | 1.371335505 | 0.513671875 | 2.363303083 | 8.5867E-230 | 5.6561E-228 | 0.654554148 | 0.127209769 | 2.39302554 | 2.1368E-70 | 1.90089E-68 | 1.679805791 | 0.319805999 |
| LanB1 | 1.273818413 | 2.53434E-22 | 1.17559E-20 | 18.50814332 | 7.654296875 | 0.811801878 | 0 | 0 | 6.407363137 | 3.650082012 | 1.032515808 | 1.3393E-88 | 1.49231E-86 | 15.49755744 | 7.57608826 |
| tkv | 1.519724985 | 1.25501E-22 | 6.07194E-21 | 2.234527687 | 0.779296875 | 2.474158577 | 4.9024E-210 | 2.8579E-208 | 0.467846044 | 0.084199016 | 0.880732312 | 2.12762E-30 | 7.0059E-29 | 6.842277552 | 3.715984065 |
| CG5390 | 2.513689998 | 3.23153E-11 | 5.23031E-10 | 0.368078176 | 0.064453125 | 2.347325248 | 9.8007E-182 | 4.5525E-180 | 0.523069567 | 0.102788409 | 1.500678089 | 3.53028E-18 | 7.11928E-17 | 0.769658927 | 0.271987655 |
| Pax | 2.107390502 | 3.63645E-27 | 2.63907E-25 | 1.557003257 | 0.361328125 | 1.51888964 | 7.8106E-128 | 2.2938E-126 | 0.517571121 | 0.180608711 | 0.932825951 | 6.59138E-15 | 1.15229E-13 | 2.104331068 | 1.102314386 |
| Idgf1 | 5.782299274 | 4.72876E-06 | 3.28857E-05 | 0.107491857 | 0.001953125 | 1.342569977 | 0.003483133 | 0.007738157 | 0.006932823 | 0.002733734 | 3.28946142 | 1.46629E-05 | 0.000121889 | 0.113604776 | 0.011619034 |
| robo2 | 1.550535019 | 3.98704E-07 | 3.46996E-06 | 0.446254072 | 0.15234375 | 0.955623607 | 9.80471E-18 | 4.69651E-17 | 0.123356443 | 0.063604884 | 1.994015907 | 5.37594E-53 | 3.44779E-51 | 2.25125121 | 0.565152118 |
| ab | 1.236259817 | 2.23975E-13 | 4.54979E-12 | 2.263843648 | 0.9609375 | 1.317478996 | 3.04057E-94 | 6.17595E-93 | 0.5041836 | 0.202296337 | 1.156896858 | 4.58061E-44 | 2.39473E-42 | 5.059201889 | 2.268931342 |
| bowl | 2.483561709 | 1.40788E-48 | 4.22317E-46 | 3.615635179 | 0.646484375 | 1.535255467 | 7.2992E-181 | 3.3505E-179 | 0.650251016 | 0.22434846 | 1.057688977 | 4.67994E-34 | 1.81776E-32 | 4.226305672 | 2.03032142 |
| CG9328 | 1.394603409 | 2.46928E-12 | 4.47104E-11 | 0.872964169 | 0.33203125 | 1.801325654 | 5.1612E-108 | 1.2484E-106 | 0.331580206 | 0.095133953 | 0.773833474 | 6.16454E-07 | 5.90287E-06 | 1.050286673 | 0.614272068 |
| CG5888 | 3.754206967 | 2.44263E-11 | 4.02586E-10 | 0.28990228 | 0.021484375 | 4.624140335 | 1.93876E-19 | 9.77681E-19 | 0.02247191 | 0.000911245 | 3.31030321 | 9.1269E-24 | 2.42016E-22 | 0.508598137 | 0.051271285 |
| CG5758 | 1.578838669 | 1.15859E-17 | 3.59521E-16 | 1.592833876 | 0.533203125 | 1.097634025 | 2.99011E-79 | 4.99452E-78 | 0.476213244 | 0.22252597 | 0.875954334 | 1.32814E-23 | 3.50497E-22 | 3.071911039 | 1.673863754 |
| CG15629 | 1.640607953 | 0.000487128 | 0.002076582 | 0.140065147 | 0.044921875 | 2.824828955 | 5.88556E-82 | 1.02418E-80 | 0.176906526 | 0.024968106 | 1.900253905 | 0.000155066 | 0.001115806 | 0.086984146 | 0.023302723 |
| CG31915 | 1.479712012 | 1.12844E-10 | 1.68965E-09 | 0.648208469 | 0.232421875 | 1.056974798 | 3.20825E-95 | 6.62035E-94 | 0.693521396 | 0.333333333 | 0.829020412 | 5.87347E-09 | 6.89258E-08 | 0.912224128 | 0.513501296 |
| Pen | 1.605284848 | 1.52733E-32 | 1.59819E-30 | 7.273615635 | 2.390625 | 2.279855577 | 0 | 0 | 6.529046139 | 1.344450519 | 1.44468939 | 0.006615613 | 0.03364538 | 0.072473676 | 0.026624744 |
| IFT52 | 2.283873524 | 1.7109E-08 | 1.8822E-07 | 0.237785016 | 0.048828125 | 0.878666863 | 1.65743E-06 | 4.85406E-06 | 0.039206311 | 0.021323127 | 2.601367806 | 0.005062203 | 0.026833812 | 0.053660747 | 0.008842335 |
| tutl | 4.010175363 | 6.27681E-68 | 4.345E-65 | 4.185667752 | 0.259765625 | 2.999674385 | 0 | 0 | 0.897920153 | 0.112265354 | 1.915566345 | 6.2693E-299 | 6.287E-296 | 14.10115674 | 3.737763436 |
| DIP-epsilon | 3.773529064 | 1.82283E-05 | 0.000113128 | 0.133550489 | 0.009765625 | 4.321639953 | 7.5792E-98 | 1.63338E-96 | 0.149414296 | 0.007472207 | 4.682252963 | 1.4284E-68 | 1.24068E-66 | 1.303478157 | 0.050769767 |
| Snoo | 2.582816081 | 1.25265E-37 | 2.04956E-35 | 4.563517915 | 0.76171875 | 2.080459961 | 1.3741E-211 | 8.2154E-210 | 0.622041597 | 0.147074904 | 0.821453445 | 1.4002E-34 | 5.45779E-33 | 16.91296205 | 9.570564343 |
| Atg18b | 2.126470443 | 4.64598E-23 | 2.40283E-21 | 0.938110749 | 0.21484375 | 1.379983939 | 8.00324E-17 | 3.7013E-16 | 0.059765718 | 0.022963368 | 0.708223572 | 0.00012999 | 0.000946479 | 0.971464276 | 0.594607381 |
| ci | 1.692332982 | 5.36311E-32 | 5.42276E-30 | 2.537459283 | 0.78515625 | 1.527014895 | 8.5523E-188 | 4.2072E-186 | 0.770499641 | 0.267359213 | 1.091343815 | 6.7316E-71 | 6.03709E-69 | 9.644216051 | 4.526262187 |
